# Inferring the multi-host fitness landscape of endive necrotic mosaic virus from cross-inoculation experiments

**DOI:** 10.64898/2026.03.18.712764

**Authors:** Lionel Roques, Julien Papaïx, Guillaume Martin, Raphaël Forien, Thomas Lenormand, Samuel Soubeyrand, Karine Berthier, Benoît Moury

## Abstract

Fitness landscapes offer a compact representation of adaptation, yet are rarely inferred from sparse multi-environment data. We present a Bayesian approach to infer an effective multi-host phenotypic fitness landscape from cross-inoculation assays by linking successful infection probabilities to Fisher’s geometrical model and to an explicit decomposition of establishment routes. The model estimates (i) the distance matrix among host-specific phenotypic optima, (ii) target-host permissiveness through the widths of fitness peaks, (iii) target-specific differences in the efficiency with which phenotypic suitability translates into successful infection, and (iv) the conditional probabilities that successful infections are attributed to direct establishment, rescue from standing variation in the source inoculum, or *de novo* rescue in the target host. We apply the approach to an experimental evolution dataset for endive necrotic mosaic virus evolved on five Asteraceae hosts and challenged in a full cross-inoculation design. The inferred landscape can be visualized as a phenotypic map of the host community, revealing pronounced heterogeneity in target-host permissiveness and a geometry broadly concordant with host phylogeny. By grounding assay-derived distances in an explicit mechanistic model, the approach provides a parsimonious representation of multi-host constraints that can be used to discuss establishment barriers and potential springboard hosts in heterogeneous communities. More broadly, it offers a general method for inferring effective fitness landscapes from sparse multi-environment data.

## 1 Introduction

Following the seminal work of Wright (1932), fitness landscapes have become a pivotal framework for understanding adaptation: they represent fitness as a function of genotype or phenotype, and they offer a simple language to describe evolutionary constraints in terms of peaks, ridges, and valleys (Gavrilets, 2004; Svensson and Calsbeek, 2012; Fragata et al., 2019). Beyond metaphor, landscapes can be coupled to mutation and other evolutionary forces to yield quantitative predictions about adaptive trajectories and evolutionary outcomes (Kauffman and Levin, 1987; de Visser and Krug, 2014). Empirically, genotype-to-fitness reconstructions in which many genotypes are constructed and measured systematically in microbes and molecular systems have shown how epistasis can constrain mutational paths and influence the repeatability of adaptation (Weinreich et al., 2006; Poelwijk et al., 2007; de Visser and Krug, 2014). However, such systematic reconstructions are feasible only in a limited set of systems where many genotypes can be constructed and fitness (or well-defined fitness proxies) can be measured directly (de Visser and Krug, 2014; Fragata et al., 2019). In most settings, available data provide only sparse and indirect readouts of performance. This motivates parsimonious landscape representations that can be inferred from tractable measurements collected across contrasted environments.

When a population encounters multiple environments, fitness is better represented not by a single surface but as a set of environment-specific fitness functions, all defined over the same trait space (Martin and Lenormand, 2015; Fragata et al., 2019). The geometry of this collection of fitness functions (how far the optima are from each other, and how sharply fitness declines away from each optimum) shapes trade-offs and helps determine how easily adaptation in one environment translates into performance in another. Pathogens circulating in multi-host communities provide a natural case where such geometry may differ among environments, since each host species represents a distinct selective environment with its own cellular constraints, immune system, and resource availability, and can therefore be viewed as imposing its own selective optimum.

Predicting pathogen emergence in such communities therefore requires linking landscape geometry to the probability that a population establishes after a host shift. Host shifts make the consequences of this geometry immediate. Establishment in a novel host often requires rapid adaptation; otherwise the pathogen population fails to persist. This is the setting of evolutionary rescue (Gomulkiewicz and Holt, 1995; Bell, 2013; Carlson et al., 2014), where persistence occurs because genotypes with higher fitness arise and increase quickly enough to reverse demographic decline. Rescue can rely on standing genetic variation, on *de novo* beneficial mutations, or on both (Barrett and Schluter, 2008; Messer and Petrov, 2013), and theory shows that rescue probability depends on mutation supply and on the initial fitness deficit in the new environment (Bell, 2013; Carlson et al., 2014). In a landscape representation, this deficit depends on the distance between the inoculum phenotype and the region of positive growth in the target environment, together with how tolerant that environment is to phenotypic mismatch.

In heterogeneous host assemblages, host diversity can either reduce or enhance emergence risk. Increasing diversity may reduce transmission by lowering the density or competence of the most permissive hosts (dilution effects; Keesing et al., 2006; Rohr et al., 2020). Conversely, adding host species that are permissive or occupy intermediate positions in the pathogen fitness landscape may sustain pathogen populations, increase mutational input, and promote host shifts, thereby acting as springboards for subsequent spillover (Woolhouse et al., 2001; Longdon et al., 2014). Such questions arise in agroecosystems, where host diversification is often proposed to limit epidemics (Mundt, 2002; Caquet et al., 2020), but it can also reshape the selective regime experienced by the pathogen and the amount of variation produced, thereby affecting evolutionary outcomes. Addressing these issues requires representations that summarize multi-host constraints in a form that is both interpretable and linked to mechanisms of adaptation.

Here we infer a multi-host fitness landscape from cross-inoculation experiments by grounding successful infection probabilities in Fisher’s geometrical model (FGM) (Tenaillon, 2014; Martin and Lenormand, 2015). In FGM, genotypes map to an *n*-dimensional phenotype, and each host corresponds to a quadratic fitness peak centered on a host-specific optimum; distances between optima quantify the phenotypic displacement required for adaptation, while peak width captures the strength of stabilizing selection and thus host permissiveness. In this setting, an inoculum sampled from a source host has an effective phenotype whose absolute growth rate in a target host is determined by (i) its distance to the target optimum and (ii) the height and width of the target peak. Under a weak-mutation-relative-to-selection approximation, referred to here as SSWM in the sense of Martin and Roques (2016) and Anciaux et al. (2018), the source population is represented by a dominant effective type accompanied by low-frequency single-step mutants. Rescue is restricted to mutant lineages one mutational step from this dominant type; sequential multistep rescue is not modeled. Establishment can then be described by explicit evolutionary-rescue formulas that account for direct establishment of the source phenotype, rescue by mutants already present in the inoculum, and rescue by mutants arising *de novo* in the target host (Martin et al., 2013; Anciaux et al., 2018). We exploit this link to turn cross-inoculation outcomes into information about landscape geometry: here, successful infection probability on a target host becomes a mechanistic function of the distance between source and target optima and of a target-specific permissiveness parameter.

We apply the approach to an experimental evolution dataset for endive necrotic mosaic virus (ENMV; species *Potyvirus chichorii*) evolved on five Asteraceae hosts and then tested in a full cross-inoculation design. Using a Bayesian approach grounded on a Markov chain Monte Carlo (MCMC) algorithm, we estimate (i) the distance matrix among host-specific phenotypic optima (including the clonal ancestor), (ii) host-specific permissiveness through fitness-peak widths on target hosts, (iii) target-specific differences in the efficiency with which phenotypic suitability translates into successful infection and, (iv) the conditional probabilities that a successful infection is attributed to direct establishment, rescue from source standing variation, or *de novo* rescue. This yields a compact map that separates geometric constraints on host shifts from target-dependent permissiveness, and provides a mechanistically interpretable summary of multi-host adaptation. It also offers a basis to discuss constraints on establishment across hosts and potential springboard hosts in heterogeneous communities. More generally, the present approach provides a general method for inferring effective fitness landscapes from sparse multi-environment data, beyond the specific system considered here.

## 2 Data

### 2.1 Study system: endive necrotic mosaic virus (ENMV)

Endive necrotic mosaic virus (ENMV; genus *Potyvirus*, family *Potyviridae*) is a positive-sense single-stranded RNA virus that infects Asteraceae hosts. The experimental study used the ENMV isolate ENMV-FR (GenBank accession KU941946), collected from lettuce (*Lactuca sativa*) in Montfavet (France) in November 2012. To start the experimental evolution from a genetically homogeneous inoculum, the field isolate was subjected to a single *Myzus persicae* aphid transmission to lettuce, yielding a clonal-derived strain (ENMV-7098MP1; hereafter ‘clonal strain’, CL) that served as the ancestor for all lineages. A thorough description of the biological system, experimental design, plant material, inoculation procedures, and additional phenotypes measured during the study is provided in the companion experimental study (Moury et al., 2026). Absolute within-host ENMV population sizes were not measured in the present experiment, although relative viral population sizes across viral populations and host species are analysed in the companion study. ENMV infections persisted over the experimental observation period, and no symptom recovery was observed.

### 2.2 Experimental design

#### Host plant species and code names

All hosts belong to the family Asteraceae. The five hosts are CH (endive, *Cichorium endivia*, tribe Cichorieae), LA (lettuce, *Lactuca sativa*, tribe Cichorieae), SA (wild salsify, *Tragopogon pratensis*, tribe Cichorieae), SO (field marigold, *Calendula arvensis*, tribe Calenduleae) and ZI (zinnia, *Zinnia elegans*, tribe Heliantheae).

The experiment consisted of two successive steps (see Moury et al., 2026 for full experimental details and greenhouse conditions).

#### Step 1: experimental evolution on each host

Experimental evolution consisted of serial infections on each of the five host species. Three-week-old seedlings were mechanically inoculated on the first two true leaves using infected-leaf material from the ancestral strain. For each host, 8 independent viral lineages were maintained through serial passages on the same host species (six infection cycles in total), yielding 5 × 8 evolved viral populations in addition to the ancestor. In the first infection cycle, 13-14 seedlings were inoculated per host species (CH, LA, SA and ZI), whereas 98 seedlings were inoculated for SO because this host is highly resistant; SO seedlings were reinoculated at 8 days post inoculation in cycle 1 because no symptoms were observed at that time. At the end of each cycle (typically 26 days post inoculation), for each lineage, the inoculum for the next cycle was prepared from 1 g of leaves harvested from one randomly chosen infected plant.

#### Step 2: cross-inoculation assay

Each of the 5 ×8 evolved viral populations was inoculated on each of the five hosts, with 12 replicate plants per (lineage, target) pair, leading to 5 ×8 ×5 ×12 inoculated plants. In addition, the clonal strain was inoculated on each host with host-dependent replication numbers, denoted *N*_CL,*k*_ (CH: 20, LA: 20, SA: 40, SO: 60, ZI: 20). Mock inoculations were included as negative controls. Infection success was assessed as systemic infection and recorded as a binary outcome (success/failure) based on serological detection (DAS-ELISA) in non-inoculated leaves at 21 days post inoculation. The model developed below is fitted exclusively to this binary systemic-infection endpoint.

### 2.3 Observed outcomes

The observed outcome is a binary infection data point. Successful infection is defined as systemic infection in an inoculated plant (ELISA-positive); failure corresponds to the absence of systemic infection. For the clonal strain, we denote by 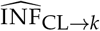 he number of observed successful infections obtained after inoculation of host *k* with the clonal strain among *N*_CL,*k*_ plants, with *N*_CL,*k*_ host-dependent.

For evolved sources, we denote by 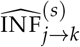 the number of successes on host *k* among 12 plants inoculated with lineage *s* ∈ {1, …, 8} evolved on host *j*. We denote by 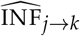 the observed number of successful infections aggregated over the eight evolved lineages. Thus, for evolved sources, with the clonal counts 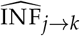, are summarized in Figure 1a, whereas Figure 1b summarizes the *N*_*jk*_ = 8 × 12 = 96 for each source-target pair *j* → *k*. The aggregated values of 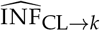, together corresponding between-lineage variability for each evolved source-target pair.

**Figure 1:**
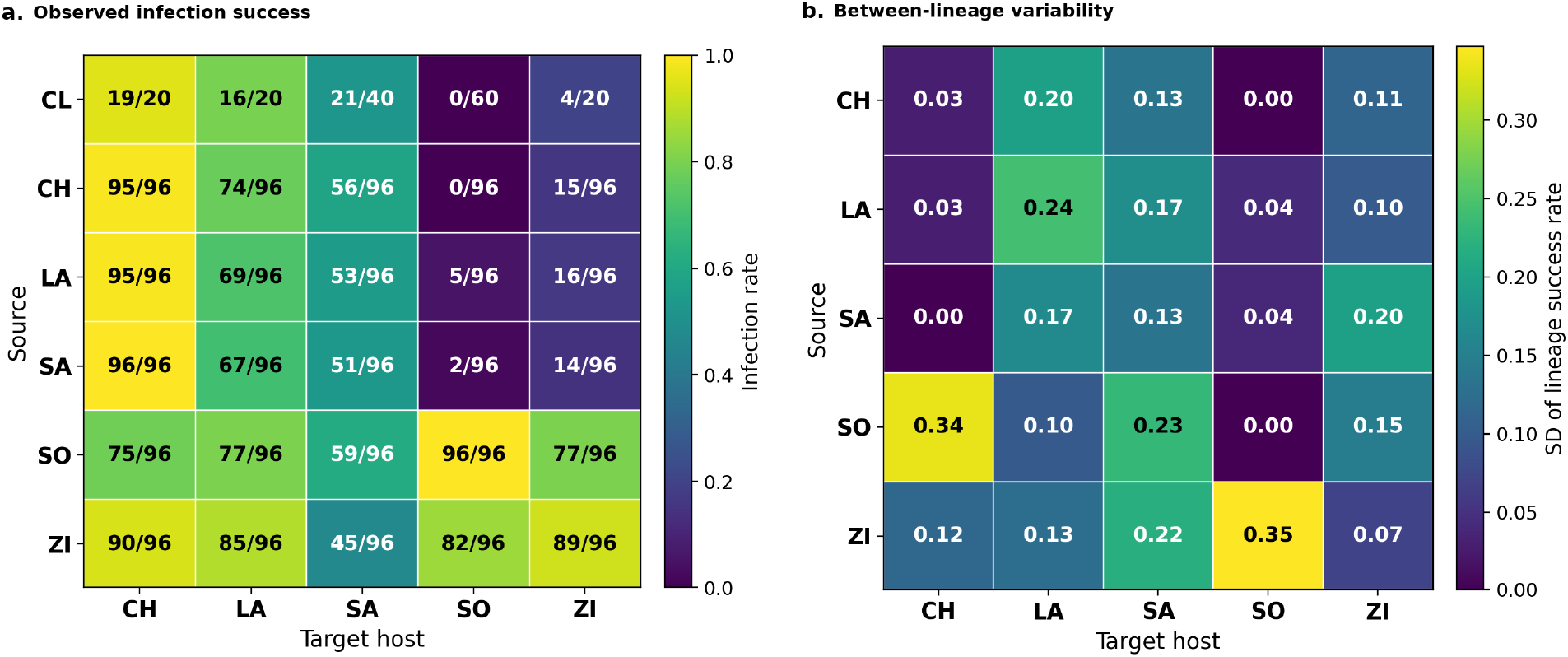
Descriptive structure of the cross-inoculation data. (a) Observed infection rates in the cross-inoculation assay. Entries are 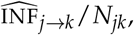, where 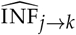 is the number of observed successful infections and *N*_*jk*_ is the number of inoculated plants. Rows correspond to the source host, or to the initial clonal strain CL, and columns to the target host. (b) Between-lineage variability in infectivity for evolved sources. Entries are the standard deviations of lineage-level infection success rates across the eight independently evolved lineages for each source-target combination; see Appendix E.

## 3 Mechanistic model

### 3.1 Overview, assumptions and notation

Our aim is to infer a mechanistic landscape from binary cross-inoculation outcomes. To make this inference tractable, we use a deliberately parsimonious model that reduces within-host dynamics to the components needed to compute a source-to-target infection probability. The model combines Fisher’s geometrical model (FGM) with a stochastic establishment model (Figure 2). FGM defines a shared phenotype space, on which a host-specific single-peak fitness surface determines the growth rate of all possible genotypes (and their associated breeding value for phenotype), in each host species. Given this geometric structure, successful infection can occur through three routes: direct establishment of the dominant source strain (EST), establishment of a mutant already present in the source inoculum (SV), or establishment of a mutant that arises *de novo* in the target host (DN). This construction yields closed-form or easily evaluated expressions for successful infection probabilities, allowing the landscape parameters to be estimated in a Bayesian framework. Table A1 lists the main quantities used below.

**Figure 2:**
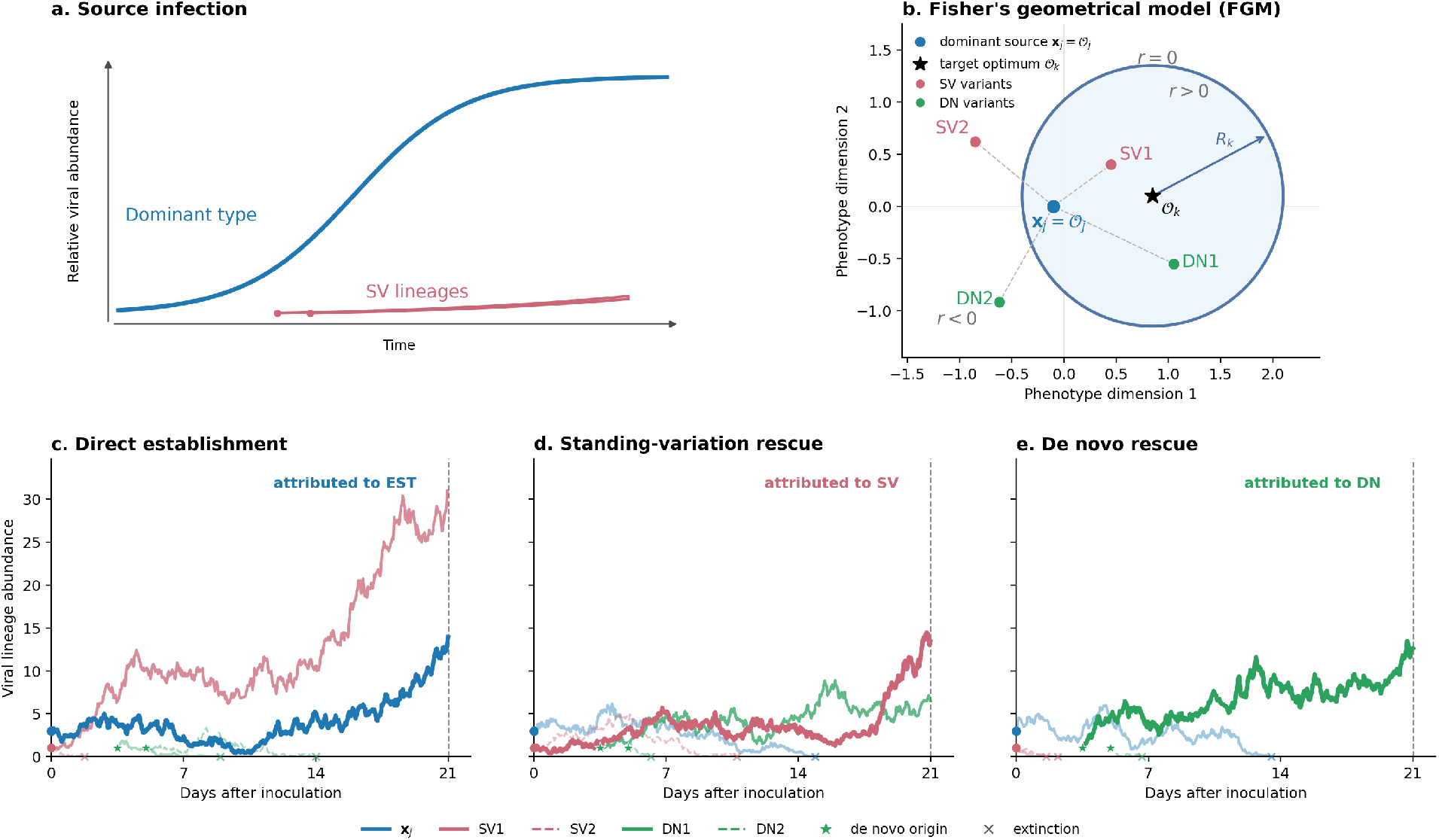
Schematic representation of the mechanistic model. (a) Source infection: the dominant type expands and saturates, while rare standing-variation (SV) lineages may arise during the early growth phase. (b) Illustrative two-dimensional FGM configuration for a particular source–target combination. The circle defines the viability boundary *r* = 0 in the target host; phenotypes inside have *r >* 0 and those outside have *r <* 0. The dominant type (in blue) is located at the source-host phenotypic optimum *O*_*j*_. Magenta and green points illustrate possible SV and DN variants, respectively. Their positions are illustrative: in a given inoculum or target-host realization, such variants may be absent or may fail to establish. (c–e) Illustrative stochastic realizations in the target host. The mechanisms are not mutually exclusive; successful infections are attributed hierarchically to EST, then SV, then DN. Panel (c) illustrates an infection attributed to direct establishment even though an SV lineage also establishes; panel (d) illustrates an infection attributed to SV although a DN lineage also establishes; and panel (e) illustrates DN rescue after failure of the dominant type and SV lineages.

### 3.2 Fitness landscape and mutational scale

Let **x** ∈ ℝ^*n*^ denote the heritable phenotype of a viral strain, with *n* treated as an unknown model dimension and compared across *n* = 1, 2, 3 below. The growth rate of strain *j* on host species *k* is assumed to depend quadratically on the distance between its phenotype **x**_*j*_ and a host-specific optimum *O*_*k*_ ∈ *ℝ*^*n*^ (Martin, 2014; Tenaillon, 2014; Martin and Lenormand, 2015):

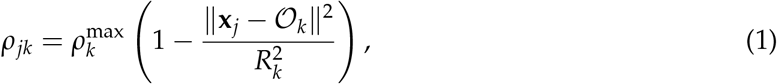

where 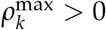 is the maximum growth rate in host *k* (peak height), attained at the optimum, and *R*_*k*_ *>* 0 is the width of the viable region around that optimum. Larger *R*_*k*_ values correspond to broader viable regions and therefore more permissive target hosts. Without loss of generality, we fix the clonal strain at *O*_CL_ = **0**, which sets the origin of phenotype space.

For the establishment–extinction calculations, time can be rescaled by a common multiplicative factor without changing asymptotic lineage-fate probabilities. We assume that the establishment or extinction fate relevant to the binary endpoint is effectively resolved within the 21-day observation window. We thus rescale time by 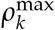 (differently in each target host species), and define the corresponding host-normalized growth rate

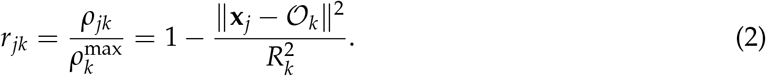

Thus, distances and radii are interpreted on this normalized growth scale. Variation in absolute peak heights 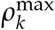 is not modeled explicitly here and does not enter the cross-inoculation probability. Biologically, this normalization does not assume equal absolute peak heights across hosts. It assumes instead that, within host *k*, 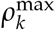 acts as a common multiplicative timescale for the demographic rates of all genotypes. After time rescaling, hosts may still differ in their optima *O*_*k*_ and permissiveness *R*_*k*_.

Mutations are modeled as isotropic Gaussian steps in phenotype space. A mutation from phenotype **x** produces

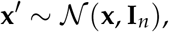

so the scale of phenotype space is set, without loss of generality, by the standard deviation of one-step mutational effects. Mutations occur at effective rate *U* per individual per unit of rescaled time. We treat *U* as common across hosts and viral lineages after rescaling by the host-specific growth timescale. Details of the mutation process can be found in Appendix B. Throughout, viruses are assumed to reproduce asexually.

### 3.3 Source inocula from Step 1: a Luria-Delbrück construction for viral source populations around the source optimum

To model establishment versus extinction in target hosts, we use an effective stochastic description of the viral population sampled from source plants at the end of Step 1 and used to inoculate target plants during Step 2. We do not model the complete within-source eco-evolutionary dynamics because their duration in viral generations is unknown. Step 1 comprised six infection cycles of approximately 26 days each and therefore likely spanned many viral generations. As a parsimonious null model, we assume that adaptation proceeded long enough for the dominant effective phenotype **x**_*j*_ to approach the source-host optimum, so that **x**_*j*_ ≈ *O*_*j*_.

In a bacterial evolution experiment along an antibiotic-dose gradient, fitness profiles measured after approximately 400 generations were consistent with populations having approached their environment-specific phenotypic optima (Harmand et al., 2018). This supports the plausibility of the near-optimum assumption but does not demonstrate that it holds in our experiment, which would require direct fitness measurements across hosts. We refer to the end-of-Step 1 state as a “mature infection” to indicate prolonged within-host growth. It consists of a dominant source type near the source-host optimum and rare segregating mutants. Mutant death during source growth is neglected; robustness to nonzero death is examined in Appendix D.

Rather than assuming mutation-selection balance, we account for the fact that the mean phenotype can approach the optimum faster than the higher moments of the phenotypic distribution equilibrate (Martin and Roques, 2016). Full equilibration may require tens of thousands of generations even in large populations evolving in stable, homogeneous environments, and possibly longer in spatially heterogeneous plant hosts subject to recurrent bottlenecks. We therefore describe source polymorphism through mutations arising during growth of a dominant type located near the source optimum.

This corresponds to a Luria-Delbrück construction, a framework that is well characterized theoretically and supported empirically (Robeva and Jungck, 2023). We adapt the large-population, rare-mutation approach of Lea and Coulson (1949) to viral burst reproduction using the stochastic pure-burst model of Hubbarde et al. (2007), while allowing for an arbitrary mutant cost. Each reproductive event replaces one particle by *B* + 1 particles, where *B*≥ 1 is the net burst increment and *B* = 1 corresponds to binary division. As *B* becomes large, the model yields a simple limit that differs from the classical Lea-Coulson result and is independent of the mutant cost. Individual-based simulations validate the analytical results and assess their robustness to nonzero viral death under a burst-death model; see Appendix D.

Finally, the branching model neglects density dependence during mature source infections. For binary division, corresponding here to *B* = 1, an appropriate change of time shows that the Lea-Coulson result can remain valid under certain forms of density-dependent growth when death is neglected (Houchmandzadeh, 2015). Whether an analogous result holds for *B >* 1 remains conjectural.

### 3.4 Stochastic demography and establishment during Step 2: a Feller diffusion approximation to early viral infection dynamics

During Step 2, the inoculum enters a new and potentially stressful target host to which its dominant source type may be maladapted. We refer to the initial phase as a “new infection”. Mortality may be substantial during this phase, but susceptible host resources are treated as effectively non-limiting, so density dependence is neglected over the early establishment-extinction timescale.

A discrete burst-death process (Hubbarde et al., 2007), or a more detailed spatial infection model, would provide a more mechanistic description of viral demography but would be less tractable. Because our main interest is whether a lineage ultimately establishes or becomes extinct, we use Feller’s diffusion approximation (Feller, 1951). After appropriate rescaling, it summarizes a broad class of branching processes through two parameters: a growth rate and a measure of demographic noise. In Appendix D, we derive the corresponding Feller coefficients from the moment dynamics of a viral burst-death process and use individual-based simulations to assess how accurately the diffusion approximates the resulting population-size distributions.

As in Step 1, we represent the population by a dominant source type, assumed to lie near the optimum of the host from which it was sampled, together with single-step mutants. These mutants may either pre-exist in the sampled inoculum or arise *de novo* in the target host. Combining the Strong Selection Weak Mutation (SSWM, in the weak-mutation-relative-to-selection sense described above) and Feller diffusion approximations makes the resulting multitype invasion process analytically tractable; see Appendix B. Using results from Feller diffusions (Martin et al., 2013) and Fisher’s geometrical model (Anciaux et al., 2018), we derive establishment probabilities for the dominant source type and its possible rescue mutants across the target-host fitness landscape.

The resulting probability of successful cross-inoculation from source *j* to target *k* depends mainly on (i) the distance between the dominant source phenotype and the target optimum, together with the target-host radius *R*_*k*_, and (ii) a composite target-specific parameter *q*_*k*_. This parameter combines the effective number of source virions available for inoculation, assumed constant across source species, the target-specific probability that an inoculated virion enters the infection process, and the scale governing early demographic stochasticity. These components are not estimated separately. The parameter *q*_*k*_ therefore measures how efficiently phenotypic suitability in host *k* is translated into detectable systemic infection. Larger values of *q*_*k*_ imply a higher establishment probability for a given growth rate of the dominant source type. Details are provided in Appendix B.

Mechanical inoculation can impose a severe host-dependent bottleneck. In other plant-virus systems, the inferred number of primary infection founders ranged from no detectable founder to approximately one hundred, depending on the host genotype (Tamisier et al., 2017). These counts provide a plausible biological order of magnitude, but cannot be mapped directly onto *q*_*k*_, which also incorporates target entry in the plant circulation system and early stochastic establishment processes. Any source-to-source heterogeneity not captured by this target filter or by the inferred source phenotype is represented statistically by a Beta-Binomial observation model (Section “Observation model and likelihood“).

### 3.5 Rescue routes and infection probability

Let *p*_*jk*_ denote the probability that inoculum from source host *j* yields detectable infection in target host *k*. If the dominant source strain has *r*_*jk*_ *>* 0, it may establish directly. If it fails to establish, or if *r*_*jk*_ *<* 0, infection may still occur through evolutionary rescue by a single-step mutant. We consider three routes to infection and denote their corresponding events by EST, SV, and DN: direct establishment of the dominant source strain, rescue from source standing variation, and *de novo* rescue in the target host, respectively.

Let INF denote successful infection and 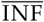 its complement. Infection fails if none of the three routes succeeds. Because SV occurs in the source host, independently of the target-host branching process, and because DN depends on the fate of the parental lineage in the target host, the failure probability factorizes as

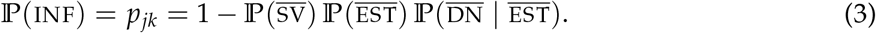

Let 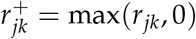 and let

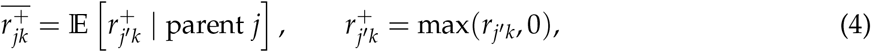

where the expectation is over random single-step mutants *j*′ from parent strain *j*. Thus, 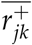 is the expected positive part of the scaled growth rate of one-step mutants in target host *k*. Define the scaled rescue-mutant production rate

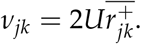

Under the Feller and single-step rescue approximations derived in Appendix B, the three failure terms are

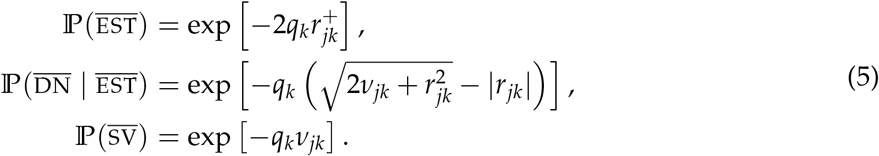

The first term describes failure of direct establishment by the dominant strain. The second term describes the absence of *de novo* rescue during the lifetime of a parental lineage that ultimately goes extinct. The third term describes the absence of rescue by mutants already present in the sampled inoculum.

Combining the three failure probabilities gives

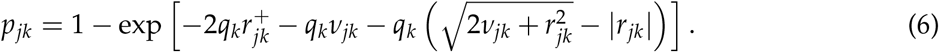

The mechanisms contributing to this probability need not be mutually exclusive. To avoid double counting, we use the following disjoint decomposition of infection outcomes:

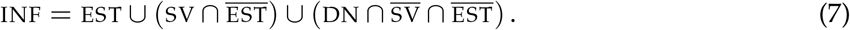

Under this decomposition, an infection is attributed to direct establishment whenever EST succeeds, to standing variation when EST fails but SV succeeds, and to *de novo* rescue when both EST and SV fail but DN succeeds.

The corresponding route-attribution probabilities, conditional on infection, are

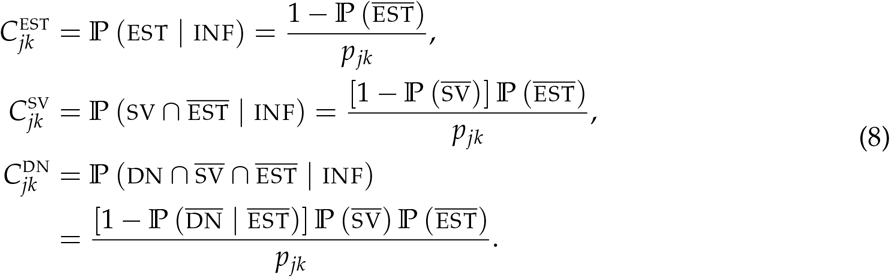

We set 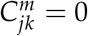 when 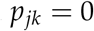. When *p*_*jk*_ *>* 0, these probabilities satisfy

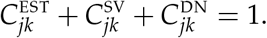

They quantify the probability that a successful infection is attributed to each route under the hierarchy EST, then SV, then DN.

The exact expression for 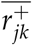 and the numerical validation of its approximation are given in Appendix C. Although 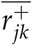 can be computed exactly under the FGM, repeated evaluation of the exact expression, which involves nested integrals, is computationally demanding and can be numerically unstable during MCMC exploration. We therefore use a matched analytic approximation,

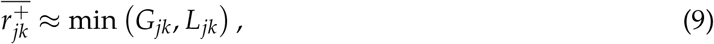

where *G*_*jk*_ and *L*_*jk*_ are Gaussian and Laplace approximations, respectively. The Gaussian approximation is accurate near the bulk of the mutant-growth distribution, whereas the Laplace approximation better captures the tail contribution when viable mutants are rare. Taking their minimum provides a stable approximation over the posterior parameter range. Their analytic expressions and accuracy assessments are reported in Appendix C.

### 3.6 How the cross-inoculation matrix informs the parameters

The relative success and failure of each source-target combination informs the distances between source and target optima. The target radius *R*_*k*_ is informed by how broadly host *k* can be infected by source phenotypes at different inferred distances: broad success across several mismatched sources supports a large *R*_*k*_, whereas success restricted to closely matched or rescued inocula supports a small *R*_*k*_. By contrast, *q*_*k*_ is a target-specific scaling factor in the exponents of all three route-specific failure probabilities. It is informed by target-column success patterns that remain after accounting for geometric mismatch: high success for phenotypically matched or nearly matched inocula supports a large *q*_*k*_, whereas incomplete infection even for well-matched inocula supports a lower *q*_*k*_. Finally, *U* is informed by the extent to which mutation-mediated rescue is required to explain infections for which the dominant source phenotype is poorly matched to the target host.

## 4 Observation model and likelihood

### Clonal inoculations

For host *k*, the number of successful infections out of *N*_CL,*k*_ inoculated plants is modeled as

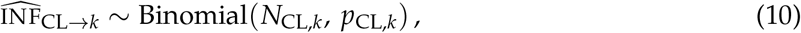

where *p*_CL,*k*_ is obtained from (6) by taking *j* = CL and **x**_*j*_ = **0**.

### Lineage-level cross-inoculations

For each source host *j* ∈ {CH,LA,SA,SO,ZI}, evolved lineage *s* ∈ {1, …, 8}, and target host *k* ∈ {CH,LA,SA,SO,ZI}, we observe 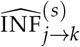 successful infections among 12 inoculated plants. Across many (*j, k*) pairs, infection outcomes vary substantially across lineages (Figure 1b and Figure E1), indicating extra-binomial variability relative to a model with a single success probability per (source, target) pair. This motivates an explicit overdispersion component in the observation model used for mechanistic inference.

We quantified this extra-binomial variability by computing pairwise dispersion factors *ϕ*_*jk*_ and found clear overdispersion relative to a Binomial model (pooled estimate 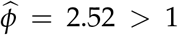; Appendix E). A natural mechanistic explanation is that independently evolved lineages from the same source host do not sit exactly at **x** = *O*_*j*_, but rather occupy a distribution of effective phenotypes around *O*_*j*_. This source-lineage heterogeneity is distinct from the SV term in the mechanistic model: SV represents rare single-step mutants present within a sampled inoculum and able to rescue infection in a target host, whereas the overdispersion term captures residual variation among independently evolved lineages. Explicitly modeling lineage-specific source phenotypes would substantially complicate the likelihood and increase computational cost. Instead, we retain the mechanistic infection probability *p*_*jk*_, derived from the effective distance *d*_*jk*_ = ∥*O*_*j*_ −*O*_*k*_∥, and account for residual lineage-level heterogeneity using a Beta-Binomial observation model.

Specifically, each lineage has its own infection probability 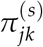, assumed to vary around the mechanistic mean *p*_*jk*_ according to a Beta distribution:

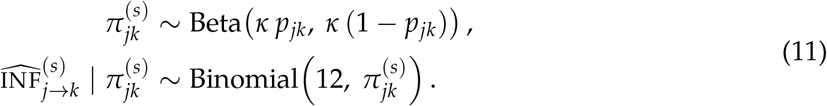

Marginally, lineage-level counts follow a Beta-Binomial distribution,

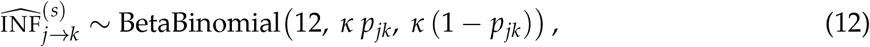

where the concentration parameter *κ >* 0 controls overdispersion (large *κ* recovers the Binomial model; smaller *κ* implies stronger lineage-to-lineage heterogeneity). We estimated *κ* from the lineage-level data prior to mechanistic inference (Appendix E) and fixed *κ* = 10 in subsequent analyses.

Assuming independence across plants and lineages, the global likelihood is the product of the five clonal Binomial terms (10) and the 5 ×8 ×5 lineage-level Beta-Binomial terms (12), where *p*_*jk*_ is defined by (6).

## 5 Estimation procedure

We perform Bayesian inference by sampling parameters in their joint posterior distribution using a Markov chain Monte Carlo (MCMC) algorithm with a Metropolis-Hastings sampler.

### 5.1 Parameters and priors

The parameters inferred in the mechanistic model are (i) the mutation parameter Λ = log_10_(*U*), (ii) the host optima *O*_*k*_ ∈ ℝ^*n*^ (with *O*_CL_ = **0** fixed), (iii) the target-specific characteristic radii *R*_*k*_ governing the width of the fitness peak for each host *k*, and (iv) the infection-efficiency parameters *q*_*k*_. We assign independent, weakly informative uniform priors: Λ ~ Uniform(−4, 4) and, for each host *k* ∈ {CH, LA, SA, SO, ZI},

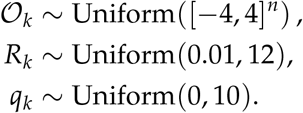

### 5.2 MCMC algorithm

We sample the posterior distribution using a random-walk Metropolis algorithm on the full parameter vector

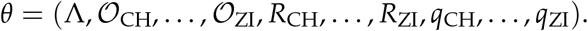

At each iteration we propose *θ*′ = *θ* + *γ*, where the components of *γ* follow independent, centered Gaussian distributions with fixed proposal standard deviations, tuned in preliminary runs to yield acceptance rates around 0.3. The likelihood is evaluated analytically using (6), with the closed-form approximation (9) of 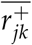.

To initialize chains near a high-probability region, we first perform a search over 200 random parameter draws and retain the best point (maximum log-likelihood) as a starting value. We then run 10 independent MCMC chains of 100,000 iterations each, discard the first 50,000 iterations as burn-in, and subsample the remaining 50,000 iterations by keeping one sample every 10 iterations. This yields 5,000 posterior samples per chain. Convergence is assessed by inspecting trace plots and verifying consistency across independent chains. We repeat the full inference for *n* = 1, *n* = 2 and *n* = 3 trait dimensions.

### 5.3 Visualization

Because absolute coordinates of optima are not identifiable up to global rotations (and reflections), we report inference in terms of the posterior distribution of the distance matrix *M* with entries *M*_*jk*_ = ∥*O*_*j*_ − *O*_*k*_∥ (including CL). To visualize the inferred geometry, we embed each posterior draw of *M* into ℝ using classical multidimensional scaling (classical MDS; Borg and Groenen, 2005, also known as principal coordinates analysis, PCoA). Classical MDS constructs a configuration of points whose Euclidean inter-point distances approximate *M* by eigen-decomposition of the double-centered matrix 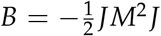, where *M*^2^ denotes the matrix with entries 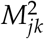 and 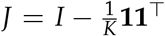 is the centering matrix (**1** is the *K*-vector of ones), and *K* = 6 (number of hosts + clonal strain CL). Let *λ*_*a*_ denote the eigenvalues of the double-centered matrix, ordered decreasingly, with associated orthonormal eigenvectors *v*_*a*_ of length *K*. Retaining the two largest positive eigenvalues, the two-dimensional MDS coordinates are obtained by scaling each eigenvector by the square root of its eigenvalue: for object *i*, the coordinate along axis *a* is 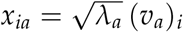, where (*v*_*a*_)_*i*_ denotes the *i*-th component of eigenvector *v*_*a*_.

Since each MDS embedding is defined only up to rotation, reflection and translation, we align the MDS coordinates across posterior samples using an orthogonal Procrustes alignment (Gower, 1975). Specifically, we select a reference configuration obtained by applying classical MDS to the posterior mean distance matrix and, for each posterior draw, apply the least-squares orthogonal Procrustes transformation consisting of a rotation or reflection followed by a translation. This yields an interpretable posterior distribution of host positions in phenotype space, while preserving the identifiability of distances.

### Software implementation and numerical reproducibility

Most analyses are implemented in Python using Jupyter notebooks and standard scientific libraries (NumPy, SciPy, pandas). The individual-based validation simulations are implemented separately in Wolfram Mathematica.

Likelihoods are evaluated using SciPy implementations of the Binomial and Beta-Binomial log-likelihoods. MCMC chains designed to draw posterior samples of parameters are run in parallel using Python `multiprocessing` with fixed random seeds for reproducibility. Posterior predictive checks are performed by simulating replicated datasets from the fitted observation model (10)-(12). Classical MDS and Procrustes alignment used for visualization are implemented directly from linear-algebra operations (eigen-decomposition and orthogonal Procrustes). Model comparison across phenotype dimensions is performed using pointwise log-likelihood contributions to compute WAIC together with approximate leave-one-out cross-validation (PSIS-LOO). The accuracy of the closed-form approximation is assessed by comparison with numerical simulations.

All data and notebooks required to reproduce the analyses are available in the public repository described in the Data Availability statement.

## 6 Results

### 6.1 Descriptive structure of the cross-inoculation matrix

Before fitting the mechanistic model, we first examined the aggregated cross-inoculation matrix of Figure 1a with descriptive analyses. We fitted binomial generalized linear models to the evolved-source counts, using source identity, target identity, and a diagonal, or sympatric, indicator as explanatory variables. The diagonal term tests for excess infection success when the source and target hosts are the same. A target-only model explained 37.6% of the null deviance, whereas a source-only model explained 19.2%. The additive source-plus-target model explained 61.2%, and adding the diagonal term increased this value to 69.7% (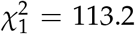, *P <* 10^−25^). Thus, the raw matrix has a strong target-host component, but this column-wise structure is not sufficient to explain the data: source identity and excess diagonal success also contribute. These descriptive patterns indicate that the cross-inoculation matrix contains both target-specific variation and source-target matching, which the mechanistic analysis must disentangle.

We next tested whether the CL, CH, LA and SA source rows had distinguishable infectivity profiles after accounting for target-host effects, since these rows appear broadly similar in Figure 1a. For these four sources, adding source identity to a binomial model with target effects did not improve the fit (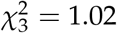, *P* = 0.80). Accordingly, we found little evidence that CL, CH, LA and SA have distinct source infectivity profiles. This result motivates a cautious interpretation of CH, LA and SA source adaptation in the mechanistic analyses below.

As another descriptive summary of Figure 1a, we treated each row of the aggregated cross-inoculation matrix as a five-dimensional vector of observed infectivity across target hosts. After empirical-logit transformation of the counts, we computed Euclidean distances between rows and applied classical multidimensional scaling (MDS; Appendix F). The first two axes separated SO and ZI from the remaining hosts, consistent with the main contrast visible in the raw infectivity profiles.

### 6.2 Phenotype dimension

We first assessed the phenotypic dimensionality required by the mechanistic model to reproduce the cross-inoculation structure using posterior predictive checks (PPC, see Appendix G for details) and predictive information criteria (Appendix G).

Posterior predictive performance is broadly similar across the three candidate phenotype dimensions. With *n* = 1, posterior predictive means already lie close to the identity line (Supplementary Figure S1a), and the PPC root-mean-square error (RMSE) is 0.05, indicating that the one-dimensional model captures the main structure of the observed infection patterns. Increasing the dimension to *n* = 2 slightly improves predictive accuracy, with posterior predictive means remaining close to the identity line (Figure 3a) and the RMSE decreasing to 0.04. Further increasing dimensionality to *n* = 3 does not lead to a substantial additional improvement: the PPC scatter remains very similar to the *n* = 2 case (Supplementary Figure S2a) and the RMSE remains essentially unchanged (RMSE = 0.04). Overall, PPC diagnostics indicate that all three models provide a reasonable description of the aggregated successful infection patterns, with only modest differences in predictive accuracy across phenotype dimensions.

**Figure 3:**
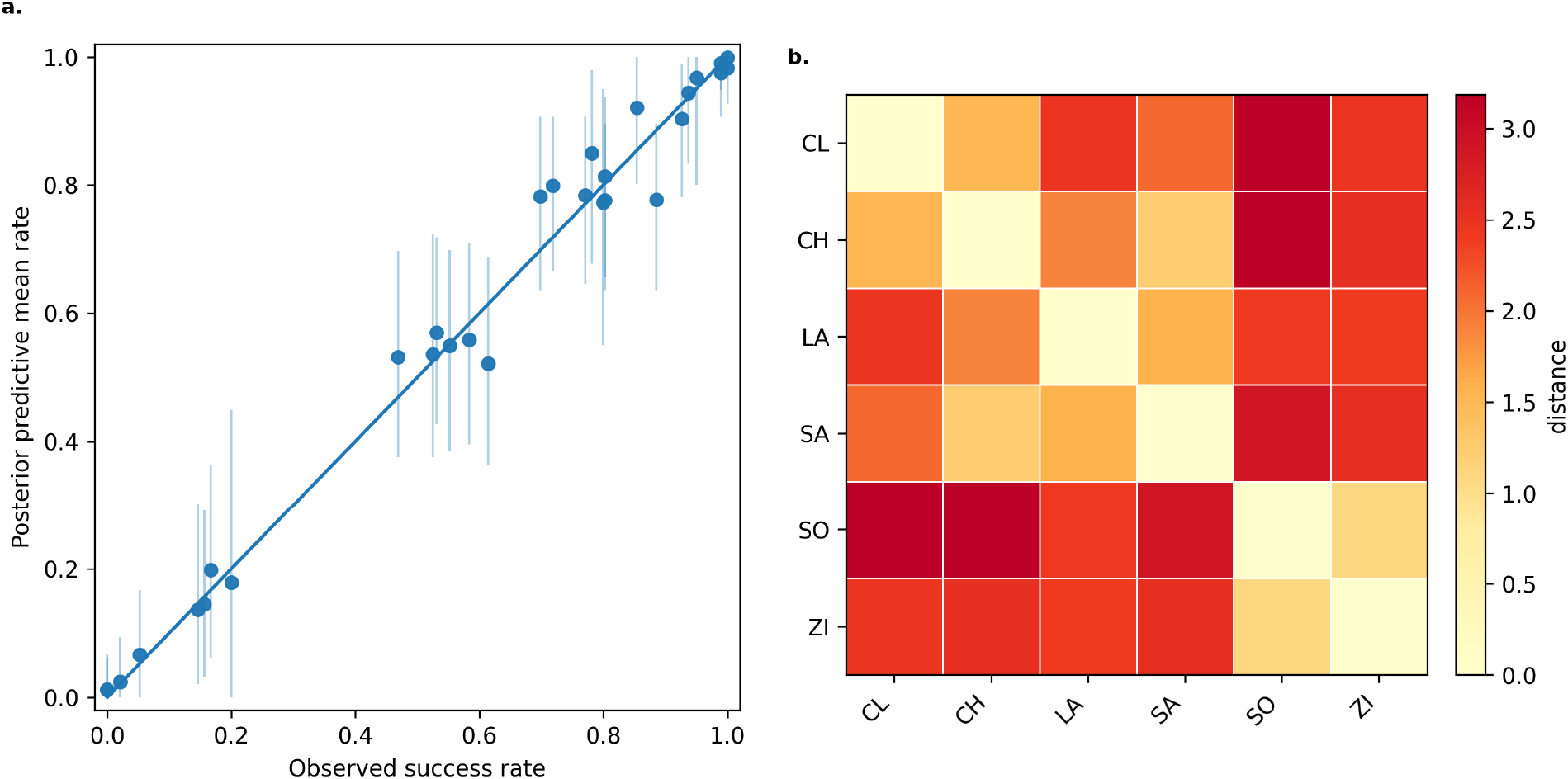
Posterior inference under the mechanistic model with *n* = 2 trait dimensions. (a) Posterior predictive check (PPC) for aggregated successful infection rates. Points show posterior predictive means versus observed rates; vertical bars denote 95% posterior predictive intervals. The diagonal line indicates perfect agreement. (b) Posterior mean distance matrix among host optima and the clonal ancestor; CL denotes the clonal strain.

To complement PPC diagnostics, we computed predictive information criteria. The Widely Applicable Information Criterion (WAIC Watanabe, 2010) strongly favors both higher-dimensional models over *n* = 1, with WAIC decreasing from 978.18 for *n* = 1 to 718.75 for *n* = 2 and 732.35 for *n* = 3. In terms of expected log predictive density, this corresponds to 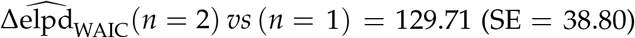 and 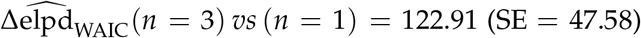. By contrast, the difference between *n* = 2 and *n* = 3 is small, with 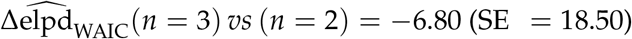, providing little support for the additional complexity of the three-dimensional model. Approximate leave-one-out cross-validation leads to the same qualitative ranking, with LOOIC values of 863.71 for *n* = 1, 708.05 for *n* = 2, and 744.75 for *n* = 3.

Taken together, PPC diagnostics and predictive criteria indicate that *n* = 2 provides the best compromise between predictive adequacy and parsimony. We therefore focus on *n* = 2 in the remainder of the Results section, and report *n* = 1 and *n* = 3 outputs in Supplementary File 1, Section S1 for completeness.

### 6.3 Posterior inference in *n* = 2 dimensions

Figure 3 summarizes posterior inference for *n* = 2 phenotypic dimensions. The posterior mean distance matrix among host optima (Figure 3b) reveals a structured multi-host geometry, with some host pairs inferred to be close in phenotype space and others strongly separated.

To visualize the inferred geometry, Figure 4 shows a two-dimensional MDS embedding of posterior distance matrices. The posterior cloud reveals well-separated regions of trait space occupied by the different host optima, confirming that the inferred landscape is genuinely two-dimensional. The spread of points around each host reflects posterior uncertainty in the relative placement of optima, while the global structure of the configuration is stable across posterior draws. Hosts that exhibit similar cross-infection profiles tend to cluster in this embedding, whereas hosts associated with strong incompatibilities are located further apart. In particular, SO and ZI form a cluster that is clearly separated from the three Cichorieae hosts.

**Figure 4:**
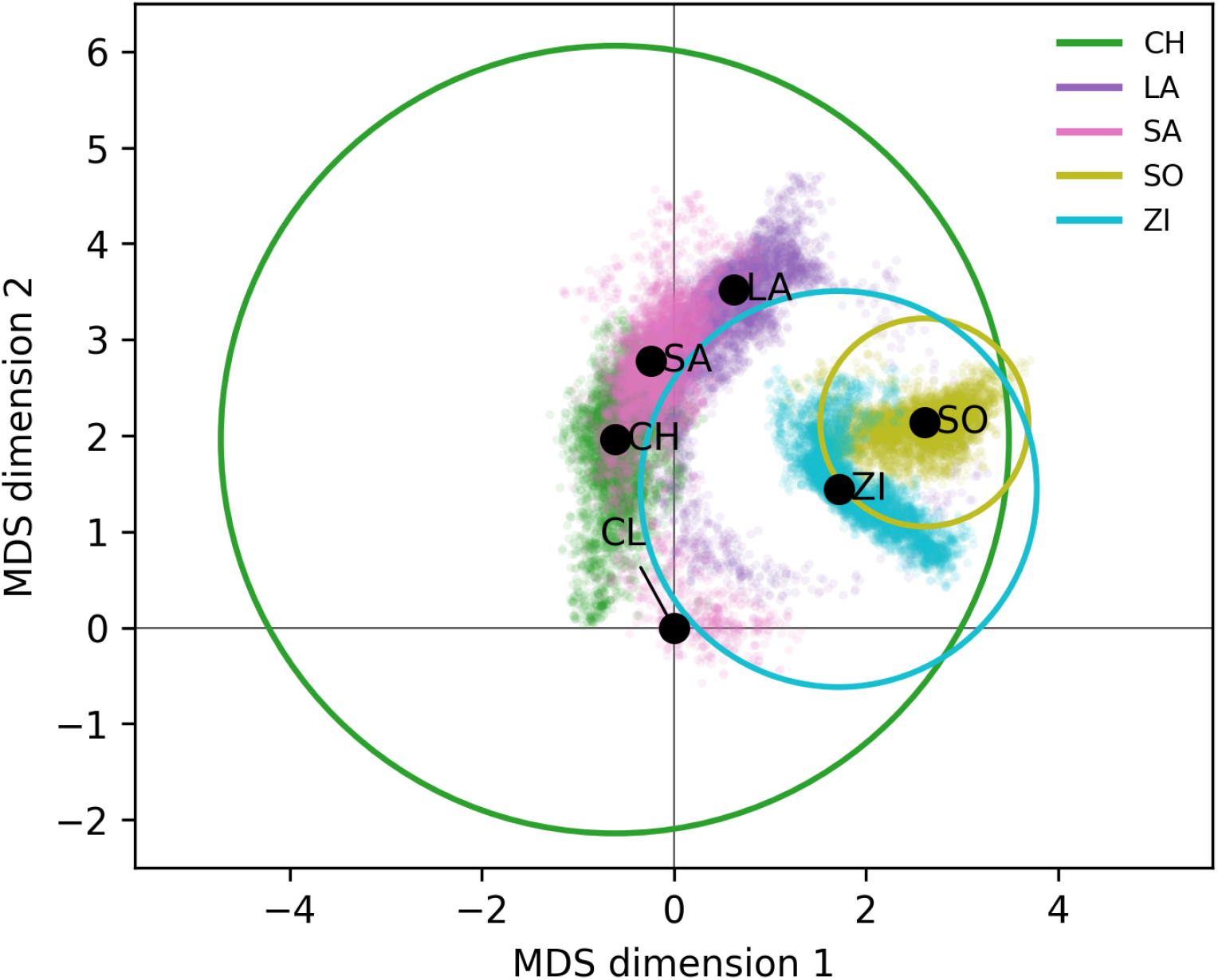
MDS visualization of the inferred host landscape with phenotypic dimension *n* = 2. Points show MDS embeddings of posterior distance matrices after Procrustes alignment; black dots indicate posterior modes of host optima. Circles represent posterior mean target radii *R*_*k*_ (permissiveness). The mean radii for LA and SA are not displayed because they extend beyond the plotting range. Coordinates are translated so that the clonal strain (CL) is located at the origin.

In addition to distances between optima, the model infers host-specific permissiveness through the characteristic radii *R*_*k*_ (Figure 5a). Posterior distributions of *R*_*k*_ reveal clear heterogeneity among hosts. SO has the smallest radius, followed by ZI, indicating restrictive targets that require a close phenotypic match for successful infection, whereas CH shows an intermediate radius. By contrast, the posterior distributions for LA and SA are shifted toward large values and accumulate near the upper bound of the prior. This pattern is expected from the model structure: for fixed phenotypic distance *d*_*jk*_, the infection probability satisfies 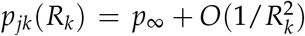 as *R*_*k*_ → ∞, with 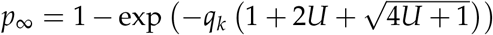. Thus, once *R*_*k*_ is sufficiently large, further increases have only a weak effect on infection probability, so the upper tail of *R*_*k*_ is only weakly identifiable from the data. Accordingly, the posteriors for LA and SA should be interpreted as evidence that these hosts are highly permissive, rather than as precise estimates of their radii. Differences in *R*_*k*_ nonetheless provide a mechanistic explanation for directional cross-inoculation effects: even when *d*_*jk*_ = *d*_*kj*_, the probability of successful infection can differ markedly between directions. For example, CH →ZI is rare (15/96), whereas ZI → CH is almost always successful (90/96; Figure 1a). Likewise, successful infection on SO is very rare for lineages evolved on the Cichorieae hosts CH, LA and SA (0/96, 5/96, and 2/96, respectively), whereas viruses evolved on SO infect CH, LA and SA readily (75/96, 77/96, and 59/96).

**Figure 5:**
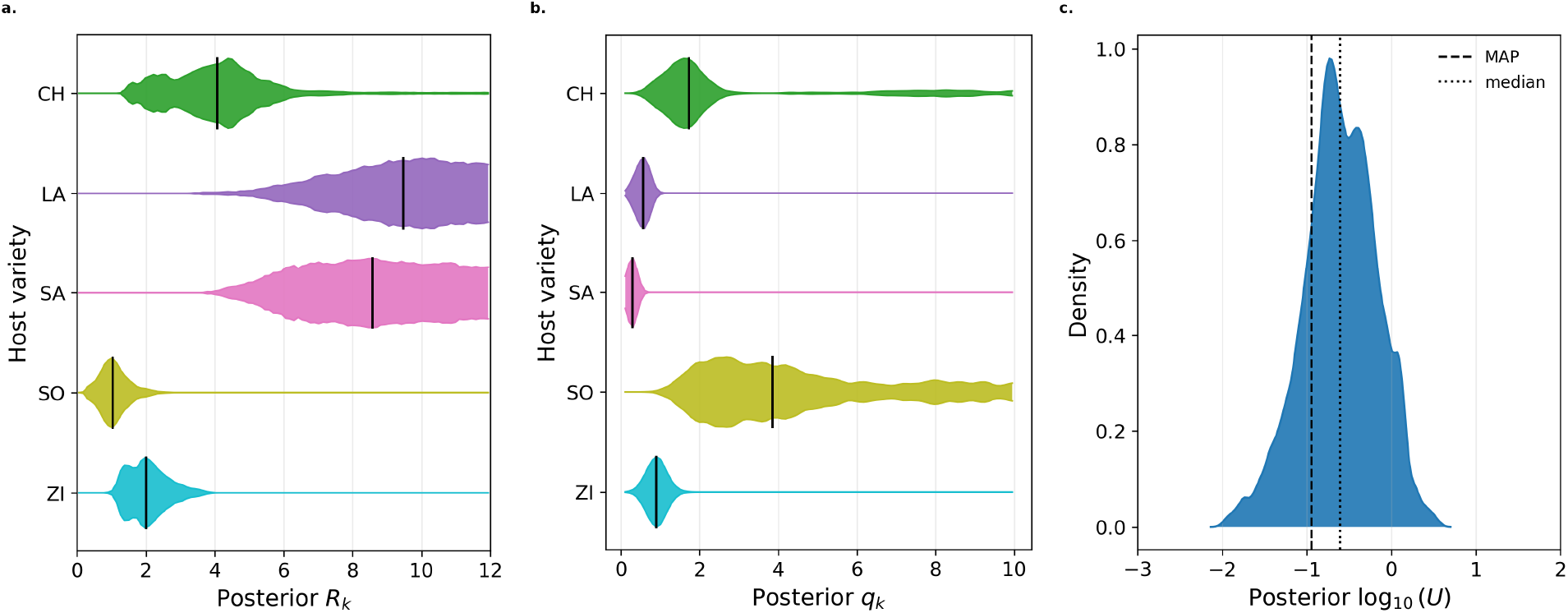
Marginal posterior distributions of model parameters for *n* = 2 trait dimensions. (a) Posterior distributions of host-specific permissiveness parameters *R*_*k*_; vertical segments indicate posterior medians. (b)Posterior distributions of target-specific infection-efficiency parameters *q*_*k*_, with the same representation. (c)Posterior distribution of the mutation parameter Λ = log_10_(*U*); dashed and dotted vertical lines indicate the maximum a posteriori (MAP) value of Λ under the joint posterior distribution of all parameters, and the posterior median, respectively.

The inferred radii also clarify when evolutionary rescue is required. Because direct establishment is possible whenever the source optimum lies within the viable region of the target, that is, when *d*_*jk*_ *< R*_*k*_, the large radii of LA, SA and CH imply that rescue is generally not required for lineages originating from CL and the other hosts to establish on them. By contrast, the smaller radii inferred for SO and ZI place the optima of CH, LA, SA and CL outside their viable regions, so that adaptation to SO or ZI from these sources typically requires rescue. However, adaptation to ZI from SO does not require rescue.

The posterior distributions of the *q*_*k*_ are shown in Figure 5b. The posterior medians are 1.7 for CH, 0.6 for LA, 0.3 for SA, 3.9 for SO, and 0.9 for ZI. These parameters complement the interpretation based on the radii *R*_*k*_: whereas *R*_*k*_ determines how far from the target optimum direct establishment remains possible, *q*_*k*_ controls how efficiently phenotypic suitability is translated into successful infection on host *k*. In this sense, SO appears as a narrow but highly effective target: its small inferred radius implies that close phenotypic matching is required, but its large *q*_*k*_ indicates that once this condition is met, successful infection is highly likely. This is consistent with the very high success of matched or nearly matched inocula on SO, including self-inoculation from SO (96/96) (Figure 1a). By contrast, LA and SA combine large radii with relatively small *q*_*k*_ values, suggesting broad viable regions but less efficient establishment conditional on phenotypic match. This is consistent with the fact that even self-inoculations on LA and SA reach only 69/96 and 51/96, respectively. CH occupies an intermediate position, combining a broad viable region with a moderately large *q*_*k*_, which is consistent with its near-perfect self-inoculation success (95/96). Overall, the posterior variation in *q*_*k*_ indicates that host-specific differences in successful infection cannot be explained by geometry alone: both the extent of the viable region (*R*_*k*_) and the local infection efficiency captured by *q*_*k*_ are needed to account for the observed asymmetries in cross-inoculation success.

Regarding the mutation parameter Λ = log_10_(*U*), the marginal posterior has median − 0.61 with a 95% credible interval from − 1.53 to 0.17; the MAP value remains of order 10^−1^. Because *U* is defined per capita and per unit of rescaled time in the rescue model, it should be interpreted as an effective mutation-supply parameter, rather than as a per-site molecular mutation probability. The marginal posterior distribution of Λ is presented in Figure 5c.

### 6.4 Contributions of direct establishment and evolutionary rescue

We next quantified how successful infections are attributed among the three infection routes by applying the decomposition in Eq. (7) and the conditional probabilities in Eq. (8) to every posterior draw of the *n* = 2 model. Figure 6 shows the posterior mean total infection probabilities *p*_*jk*_ together with the posterior mean conditional attribution probabilities 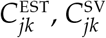, and 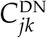.

**Figure 6:**
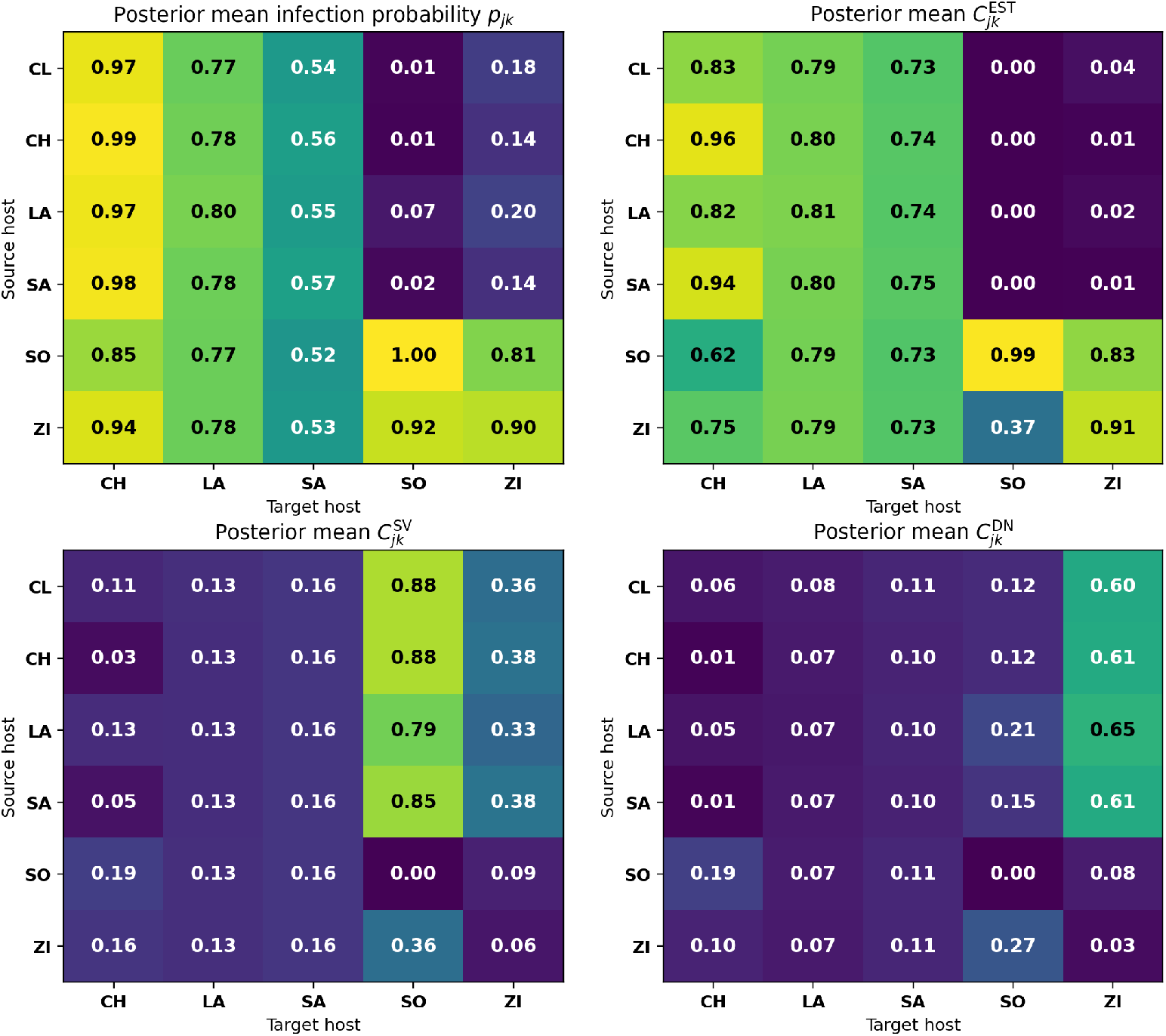
Posterior decomposition of infection probabilities under the *n* = 2 model. The top-left panel shows the posterior mean total infection probability *p*_*jk*_. The remaining panels show the posterior means of the conditional route-attribution probabilities 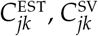, and 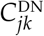, defined by the decomposition in Eq. (8). For each posterior draw and each source-target pair with *p*_*jk*_ *>* 0, the three attribution probabilities sum to 1. Rows correspond to source hosts and columns to target hosts.

The main qualitative pattern is that infection of the relatively permissive Cichorieae targets CH, LA, and SA is predominantly attributed to direct establishment. Across most source-target combinations involving these three targets, the posterior mean value of 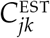 lies between 0.73 and 0.96. The lowest value is obtained for infection of CH by the SO-derived inoculum, for which direct establishment still accounts for 0.62 of the conditional attribution probability. This is consistent with the inferred geometry: among evolved source optima, SO is the most distant from CH, so the SO-derived dominant phenotype is the least well matched to the CH target despite the relatively broad CH viable region. The corresponding attribution probabilities for standing variation and *de novo* rescue are limited for these permissive targets.

A different pattern emerges for the more restrictive ZI and SO targets. For infection of ZI from CL-, CH-, LA-, and SA-derived inocula, the posterior mean total infection probabilities are 0.18, 0.14, 0.20, and 0.14, respectively. Conditional on infection, *de novo* rescue is the dominant attributed route, with posterior mean values of 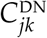 equal to 0.60, 0.61, 0.65, and 0.61, respectively. Source standing variation accounts for most of the remaining attribution probability, with 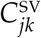 ranging from 0.33 to 0.38, whereas direct establishment contributes little.

The SO target imposes an even sharper filter on these same source inocula. Posterior mean infection probabilities from CL-, CH-, LA-, and SA-derived inocula are 0.01, 0.01, 0.07, and 0.02, respectively. The attribution probability for direct establishment is close to zero, and the rare predicted infections are predominantly attributed to source standing variation, with posterior mean values of 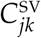 equal to 0.88, 0.88, 0.79, and 0.85, respectively. The remaining attribution probability is assigned to *de novo* rescue. The contrast with ZI follows directly from the rescue formulas. For a subcritical parental type (*r*_*jk*_ *<* 0), the *de novo* term contains 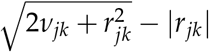, which, for fixed *ν*_*jk*_, decreases as the magnitude of maladaptation |*r*_*jk*_| increases, because a more strongly declining parental lineage accumulates less replication before extinction. The standing-variation term, *q*_*k*_*ν*_*jk*_, does not contain this additional target-host persistence penalty because those mutants are already present at inoculation. Thus, the sharper SO filter shifts the attribution toward standing variation, whereas on ZI the parental lineage has more opportunity to generate rescue mutants *de novo*.

The reciprocal SO-to-ZI and ZI-to-SO inoculations illustrate that similar total infection probabilities can arise through different combinations of routes. SO-to-ZI infection has a high posterior mean probability of 0.81 and is predominantly attributed to direct establishment 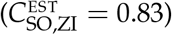. ZI-to-SO infection also has a high posterior mean probability of 0.92, but its attribution is more evenly distributed among direct establishment 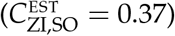, source standing variation 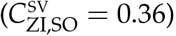, and *de novo* rescue 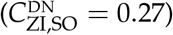. This directional difference is consistent with the broader viable region inferred for ZI than for SO: the SO-adapted phenotype lies within the ZI viable region, whereas the ZI-adapted phenotype lies closer to the narrower SO viability boundary, leaving a larger role for stochastic loss and rescue. Thus, the mechanistic model adds information beyond the raw MDS map by distinguishing permissive targets that accept many source phenotypes through direct establishment from restrictive targets for which infection is primarily attributed to evolutionary rescue.

## 7 Discussion

Our goal was to infer a compact, mechanistically interpretable multi-host fitness landscape from cross-inoculation outcomes. We represent each host as a fitness peak in a shared phenotypic space and estimate the geometry of host-specific optima (including the clonal ancestor), the characteristic radii *R*_*k*_ that determine the extent of each host’s viable region, and the target-specific infection-efficiency parameters *q*_*k*_ that control how efficiently phenotypic suitability is translated into successful infection. Establishment probabilities are linked to this landscape using Fisher’s geometrical model (FGM) together with a weak-mutation, single-step rescue approximation. The resulting phenotypic map of the host community (Figure 4) provides a parsimonious summary of multi-host constraints on successful infection. Below, we discuss the biological interpretation of this map, its implications for host shifts and potential springboard effects, and the key modeling assumptions and limitations.

### 7.1 A phenotypic map of host community structure

Figure 4 condenses a high-dimensional dataset (lineage-resolved cross-inoculation outcomes) into a low-dimensional object that can be read as a map of host-associated selective constraints. The positions of the optima encode which host shifts require large phenotypic change (large inter-optimum distances) and which host environments are close in the mechanistic sense relevant to rescue. The radii *R*_*k*_ quantify how tolerant each target host is to phenotypic mismatch, thereby separating two conceptually distinct drivers of infection success: where the optima are located (geometry) and how wide each peak is (permissiveness).

This mechanistic map is consistent with the descriptive MDS of the raw infectivity profiles, which also separates SO and ZI from the remaining sources (Appendix F). The mechanistic model adds interpretation by decomposing this pattern into target geometry, permissiveness, infection efficiency and rescue routes, while the posterior clouds show the uncertainty associated with the inferred host positions.

This representation is reminiscent of antigenic cartography, where multidimensional scaling is used to map assay-derived relationships and visualize cross-protection (Smith et al., 2004; Bedford et al., 2014). Here, geometric distances are grounded in an evolutionary-rescue approximation under the FGM, and can be interpreted in terms of maladaptation and target-specific viability thresholds.

### 7.2 Host relatedness and clustering: a phylogenetic signal hypothesis

The inferred landscape reveals a structure that is broadly concordant with host taxonomy. Posterior clouds form two clearly separated clusters: CH, LA and SA, all belonging to the Cichorieae, and a second cluster formed by SO (Calenduleae) and ZI (Heliantheae). This separation is consistent with the deeper phylogenetic split between tribe Cichorieae (belonging to subfamily Cichorioideae) and the tribes Calenduleae and Heliantheae, belonging to subfamily Asteroideae, which are more closely related to each other than to Cichorieae (Mandel et al., 2019). Within the Cichorieae cluster, inferred optima fall within one another’s *R*_*k*_ neighbourhoods, indicating substantial phenotypic compatibility across these hosts: the phenotypic displacement associated with successful infection on one Cichorieae host is largely sufficient for success on the others. This interpretation is qualitatively consistent with Moury et al. (2026), where little or no nonsynonymous change was detected in the VPg (viral protein genome-linked) cistron, a genome region of potyviruses essential for infectivity and adaptation to host plants (Pollari et al., 2024), of ENMV for populations evolved on Cichorieae hosts.

By contrast, the SO/ZI cluster is separated from the Cichorieae cluster, and their neighbourhoods largely exclude the Cichorieae optima, implying limited cross-compatibility between these environments and a larger effective shift across tribes. This is consistent with the strong molecular signature of adaptation in Asteroideae, where parallel amino-acid substitutions in VPg at codons 122–125 and 165 were repeatedly selected in ZI and shown (alone or in combination) by reverse genetics to increase infectivity and/or systemic load (Moury et al., 2026). Yet Moury et al. (2026) also report frequent VPg-mediated cross-adaptation across tribes, with no detectable enrichment within tribes. This suggests that the relationship between host geometry and mutational routes should be interpreted with caution.

More generally, the geometry is consistent with the hypothesis that distances between inferred optima, together with the clustering of posterior support, may carry phylogenetic signal while allowing for heterogeneity among closely related hosts (see discussion of phylogenetic signal in host-pathogen associations Gilbert and Webb, 2007). Expanding the host panel within Asteraceae would allow formal tests of correlations between inferred geometry and phylogenetic distance, and could help relate deviations to measurable host traits or resistance mechanisms.

### 7.3 Permissiveness as a driver of asymmetry, and alternative mechanisms

Directional asymmetries in cross-inoculation success are clearly visible in the data (Figure 1a). In our model, two target-dependent components contribute to such asymmetries, but act at different levels. The characteristic radius *R*_*k*_ determines the geometric extent of the viable region around the optimum of host *k*, and therefore whether direct establishment is possible without rescue. By contrast, *q*_*k*_ controls how efficiently phenotypic suitability is translated into successful infection once the lineage is in that viable region.

The posterior results show that differences in host permissiveness cannot be reduced to geometry alone. Hosts differ not only in the width of their viable regions, but also in the local efficiency of establishment, as captured by *q*_*k*_. However, *q*_*k*_ alone cannot explain the strongest directional patterns in the data. For example, successful infection from CH to SO was never observed, whereas the reverse direction was frequently successful, despite the fact that SO is inferred to be highly favorable to establishment once close phenotypic matching is achieved. This indicates that the major asymmetries arise from the joint action of the two components: *q*_*k*_ modulates infection efficiency, but the geometry encoded by *R*_*k*_ remains essential because it determines whether the target host is reachable without rescue.

Several alternative, and potentially concurrent, mechanisms could also generate such asymmetries. One possibility is source-dependent inoculum quality, such that the effective inoculum size differs among source populations. Another is variation in the mutational input available for rescue. Biologically, this input may reflect both the baseline mutation rate and the genetic basis of adaptation to the target host, i.e., the number of sites that could mutate to improve fitness on the new host. In our approach, these components are not disentangled and are effectively absorbed into the rescue process. The inferred landscape should therefore be interpreted as an effective landscape for successful infection, rather than a direct representation of phenotypic trade-offs among potential hosts. This is both a strength and a limitation: it captures the overall constraints governing successful infection, but does not isolate phenotypic mismatch from mutational accessibility or other sources of asymmetry. Disentangling these effects would require additional observations, such as the identification of mutations involved in adaptation to novel hosts, direct fitness assays, within-host viral load trajectories, and between-host transmission.

From a statistical standpoint, our lineage-resolved Beta-Binomial likelihood already acknowledges that a single deterministic mapping (*j, k*) ↦*p*_*jk*_ is an approximation, by allowing among-lineage heterogeneity around the mechanistic mean. This relaxes the distance-based approximation without forcing the landscape geometry or the target-specific parameters to account for all lineage-level variation.

### 7.4 What the map suggests about host shifts and springboard effects

The phenotypic map provides a way to discuss constraints on infection success across hosts and potential springboard effects in a multi-host community. In this approach, a host can facilitate emergence on other hosts through two non-exclusive mechanisms: by being readily infected from diverse sources, which allows pathogen populations to persist and diversify there; and second, by occupying a geometric position such that adaptation on this host brings lineages closer to other host optima.

These mechanisms imply that hosts with broad viable regions and efficient establishment should be readily infected from diverse sources and may amplify pathogen diversity, whereas restrictive hosts act as filters: successful infection is unlikely unless the source phenotype lies close to the target viability region or has evolved toward it. The ENMV data illustrate this pattern: successful infection on SO and ZI is rare for lineages evolved on Cichorieae hosts, yet viruses evolved on SO and ZI show broad cross-infection success (Figure 1a).

### 7.5 Applications: designing rugged fitness landscapes to slow pathogen evolution

This springboard perspective connects directly to debates on host diversification: mixtures can reduce disease spread via dilution, but can also promote evolutionary emergence under some community structures (Mundt, 2002; Keesing et al., 2006; Caquet et al., 2020; Rohr et al., 2020). A phenotypic landscape representation offers a way to articulate when diversification is expected to be ‘safe’ (e.g. hosts far apart and restrictive, limiting successful host shifts) versus potentially risky (e.g. inclusion of permissive or central hosts that sustain populations and generate variants). More generally, our approach for inferring a pathogen fitness landscape across multiple hosts has direct implications for designing interventions that deliberately increase landscape complexity and thereby impede general adaptation.

In plant pathology, host genetic heterogeneity created by cultivar mixtures, multilines, or landscape-scale mosaics can simultaneously reduce epidemics and alter adaptation in pathogen populations, with strong empirical support and long-standing theory on mixture-based disease management and resistance durability (Wolfe, 1985; Mundt, 2002, 2014). A field demonstration showed that increasing within-field host diversity can provide substantial disease control (Zhu et al., 2000), while more recent evolutionary-epidemiological work emphasizes that deployment strategies should be evaluated for both epidemiological efficiency and evolutionary robustness (McDonald and Linde, 2002; Zhan et al., 2015; Rimbaud et al., 2021). Our estimated fitness landscape provides a data-driven way to select cultivar sets (or crop mixtures) that maximize reciprocal constraints - e.g., combining peaks separated by deep valleys - thereby making evolutionary escape less likely under realistic dispersal and mutation.

Analogously, in clinical settings, combination and sequential antibiotic therapies can be interpreted through the same lens as our host-based analysis: they are attempts to engineer a rugged fitness landscape across drugs, so that adaptation to one environment incurs predictable costs in others (Yeh et al., 2009; Bollenbach, 2015; Tyers and Wright, 2019; Römhild et al., 2022). In this view, drug-drug interactions and resistance trade-offs shape the height and connectivity of peaks, while collateral sensitivity creates exploitable valleys whereby evolutionary escape from drug A entails increased susceptibility to drug B (Imamovic and Sommer, 2013; Pál et al., 2015; Römhild and Andersson, 2021). Sequential or cycling protocols then aim to push populations toward states from which the next drug is especially constraining (Imamovic and Sommer, 2013; Nichol et al., 2015). This motivates fitness-landscape-informed designs that explicitly quantify which adaptive transitions are likely under a given treatment schedule, and that use principled combination design to reduce evolutionary escape routes (Nichol et al., 2019; Bognár et al., 2024).

### 7.6 Phenotypic dimensionality and robustness of the inferred landscape

A methodological result of this study is that *n* = 2 provides the best-supported compromise between predictive adequacy and parsimony. This connects our approach to broader theory showing that trait dimensionality can strongly affect the extent of local adaptation across heterogeneous environments. In particular, MacPherson et al. (2015) showed, using a multivariate evolutionary model and reciprocal-transplant data, that the number of traits under spatially variable selection can explain substantial variation in local adaptation. Their framework also emphasizes that multi-dimensional structure can be inferred from performance matrices by estimating the positions of population phenotypes and local optima in a latent trait space. Our approach is related in spirit, but differs in that the matrix entries are cross-inoculation outcomes and the latent geometry is linked mechanistically to target permissiveness, stochastic establishment and evolutionary rescue. While the one-dimensional model (*n* = 1) captures part of the cross-infection structure, predictive criteria favor two phenotypic dimensions (Section “Phenotype dimension“). Increasing dimensionality beyond two yields only marginal improvements in posterior predictive checks, and the *n* = 3 model is not favored by WAIC. Moreover, the qualitative geometry inferred for *n* = 3 remains close to that obtained for *n* = 2 when projected onto two dimensions (Supplementary File 1, Section S1). This suggests that the main geometric features recovered under *n* = 2 are not merely artifacts of constraining the landscape to a plane, but reflect structure that is robustly supported by the data.

Beyond statistical support, two-dimensional maps are also easier to communicate and compare across systems, and may help guide experimental design, for example by suggesting additional hosts expected to lie between already inferred optima.

### 7.7 Model assumptions and limits of interpretation

#### Peak-height normalization and host resistance

The model does not estimate host-specific absolute peak heights 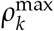. We assume an unnormalized quadratic form, but the likelihood is written in terms of the normalized growth rate 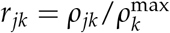 so that *r*_*kk*_ = 1 at the target optimum. Under this normalization, variation in 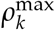 is not identifiable from binary endpoint infection data. A lower absolute peak height, stronger innate resistance, slower systemic movement, or reduced access of inoculated virions to susceptible tissue would reduce the probability that a viable phenotype yields detectable systemic infection, but these effects are not separated in the present likelihood. Their effects on binary infection success are absorbed into the composite establishment-efficiency parameter *q*_*k*_ and, for mutation-mediated routes, into the effective timescale implicit in *U*. Quantitative viral-load time series or direct within-host growth-rate measurements would be required to estimate 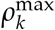 separately. This point is especially relevant for SO, which was the most resistant host at the start of the experiment: many more SO seedlings were inoculated during the first cycle and the clonal strain produced no systemic infections in SO in the cross-inoculation assay. In the inferred landscape, this initial resistance is represented primarily by a very small target radius *R*_SO_, combined with a large *q*_SO_: SO is therefore interpreted as a narrow target that is difficult to enter from mismatched source phenotypes, but highly efficient once a compatible phenotype is present. If SO resistance instead reflected only a lower absolute peak height, the present binary data would not distinguish that mechanism from a change in *q*_*k*_; this is why we interpret the geometry as an effective establishment landscape rather than a direct measurement of all within-host resistance components.

#### Source-population heterogeneity

The raw profile tests show that the CL, CH, LA and SA source rows cannot be distinguished with the aggregate data once target-host effects are accounted for (Section “Descriptive structure of the cross-inoculation matrix“). This result calls for a conservative interpretation of source adaptation on these hosts. In particular, the fitted landscape should not be read as strong evidence that CH-, LA- or SA-evolved inocula reached distinct adapted source phenotypes during Step 1. Rather, the data are compatible with the simpler possibility that CL-like phenotypes already had sufficient infectivity on these targets, either because the corresponding target optima are close to CL, because their viable regions are broad, or both.

To assess this point directly, we fitted an alternative model with *n* = 2 in which the effective source phenotypes for CH, LA and SA inocula were set to the clonal phenotype, **x**_*j*_ = *O*_CL_ = **0** for *j* ∈{CH, LA, SA}, while CH, LA and SA were retained as distinct target hosts with their own optima, radii and infection-efficiency parameters (Supplementary File 1, Section S2). This alternative model achieved essentially the same fit as the baseline model, with a slightly lower maximum log-likelihood (Δ log *L* = − 1.3 relative to the baseline model). Thus, the data do not require CH, LA and SA source inocula to be treated as distinct adapted phenotypes.

However, the alternative model produced a less interpretable landscape, with broader posterior uncertainty on the optimum positions (Supplementary Figure S3c). In this model, CH-, LA- and SA-derived inocula are no longer represented by their corresponding host optima but are constrained to be CL-like. Consequently, the points labeled CH, LA and SA in the inferred map can only be interpreted as target-host optima, not as evolved source phenotypes. Distances involving these points therefore no longer measure similarity among source inocula. Moreover, because CH, LA and SA are relatively permissive targets, the data provide little leverage to localize their optima: different positions can be compensated by broad viable regions and yield similar infection probabilities. The main conclusions concerning the restrictive SO and ZI targets and target-specific permissiveness nevertheless remain unchanged.

#### Mutants generated in mature source infections

Viral accumulation in inoculated leaves may be approximated by an initial population expansion followed by saturation (Jayet et al., 2026). Accordingly, the Luria–Delbrück argument used for the SV term should be interpreted as an early-growth and rare-mutant approximation: parental growth is approximated as exponential during the initial expansion phase, rather than over the complete infection trajectory (Appendix B). If the source population reaches saturation rapidly, the production of late-arising source mutants would be reduced relative to this approximation. In addition, mutant lineages generated in the source host may be lost stochastically, remain at low abundance, or fail to contribute to the sampled inoculum.

To assess the impact of this assumption, Supplementary File 1, Section S3 refits the *n* = 2 model after introducing a fixed attenuation factor *α*_SV_ in the source-standing-variation failure probability: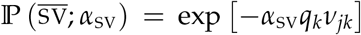. The main model corresponds to *α*_SV_ = 1, whereas *α*_SV_ = 0.5 reduces the contribution of source standing variation and *α*_SV_ = 0 removes it. The alternative fits have maximum log-likelihoods very close to that of the main model. Their posterior predictive checks, distance matrices, MDS maps and target-radius distributions preserve the same qualitative structure: SO and ZI remain separated from the CL/Cichorieae group, SO remains a restrictive target, and SO- and ZI-derived inocula retain broad infectivity (Supplementary Figures S4 and S5). Thus, the effective magnitude of the SV term affects the mechanistic attribution of some infections, but the main inferred host-shift constraints are not driven by this source-standing-variation assumption.

#### Scope of the SSWM approximation and clonal interference

Viruses can evolve with substantial standing diversity and clonal interference (Miralles et al., 1999; Desai and Fisher, 2007). We therefore do not claim that ENMV populations as a whole evolved as strictly monomorphic populations through non-overlapping selective sweeps.

We use SSWM in the specific FGM sense of a weak-mutation-relative-to-selection approximation (Martin and Roques, 2016; Anciaux et al., 2018): on the relevant rescaled timescale, the mutation rate is assumed to be small relative to a characteristic magnitude of the selection effects of random one-step mutations. This condition does not require the population-wide mutation supply to be smaller than one and therefore does not exclude the simultaneous presence of multiple low-frequency mutant lineages. Operationally, each source inoculum is represented by a dominant effective type and low-frequency single-step mutants. Rescue is restricted to mutant lineages one mutational step from the dominant type. The companion molecular results provide qualitative support for a single-step approximation because adaptation to restrictive hosts involved a small number of recurrent amino-acid substitutions in VPg with detectable effects (Moury et al., 2026). These observations do not, however, demonstrate that beneficial lineages never segregated simultaneously or that the weak-mutation-relative-to-selection condition was quantitatively satisfied.

The transition from mutation-limited evolution to a regime with concurrent beneficial lineages depends jointly on effective population size, beneficial mutation input, and selection coefficients (Desai and Fisher, 2007; Martin and Roques, 2016). The present binary cross-inoculation data do not identify these quantities separately. Although census abundances can reach 10^11^–10^12^ virus particles per infected leaf in some plant-virus systems (Harrison, 1956), census abundance cannot be equated with the effective number of competing lineages because within-host colonization involves recurrent bottlenecks (Gutiérrez et al., 2012).

The fitted *U* is an effective mutation-supply parameter per capita and per unit of rescaled time, estimated conditionally on the single-step model. It cannot therefore be used as an independent test of the SSWM assumption and is not directly comparable with a per-site molecular mutation rate. No direct molecular mutation-rate estimate is available for ENMV. For Turnip mosaic virus, a member of the same genus, a rate of order 10^−6^ substitutions per nucleotide per replication has been reported (de la Iglesia et al., 2012). This value provides a molecular order of magnitude for a related virus but cannot be mapped directly onto *U*, which also incorporates the demographic turnover and the rescaled time available for mutation and rescue.

### 7.8 Applicability of the approach

Although we developed the method for host shifts in a plant virus, it could be applied much more broadly. The environmental axis need not correspond to host species: it could represent any set of selective conditions for which cross-environment performance can be measured, such as different antibiotic doses for bacteria (Harmand et al., 2018), or different temperature regimes for pathogens. Likewise, the workflow is not restricted to binary establishment outcomes. It could be adapted to other response variables, including direct fitness measurements or quantitative viral-load measurements such as ELISA absorbance values, provided that an appropriate mechanistic model and observation model are specified. In the ENMV dataset, ELISA absorbance values combine two components: whether systemic infection was established and, among infected plants, how much virus accumulated in systemic tissue. Because the latter component reflects post-establishment processes, including within-host replication, movement, accumulation, and measurement variation, incorporating viral load would require a two-stage model for infection success and viral accumulation conditional on infection, together with additional biological and observation parameters. This illustrates a broader point: the response variable should match the biological process represented by the inferred landscape. In this sense, the present method can be viewed as a template for inferring effective fitness landscapes from sparse multi-environment data, beyond the specific system considered here.

## Data availability

The lineage-resolved cross-inoculation dataset analyzed in this study, together with the code required to reproduce all analyses, figures, and Appendix results, are available at Zenodo: https://doi.org/10.5281/zenodo.21728298.

## Acknowledgments

We are grateful to the editor and the anonymous reviewers for their careful reading and constructive comments, which helped to substantially improve the manuscript.

## Author contributions

All authors designed the study. K.B. and B.M. conducted the experimental work. L.R. and G.M. developed the mechanistic model. L.R., J.P., and S.S. developed the statistical inference. L.R. wrote the code and performed the analyses. All authors interpreted the results. L.R. wrote the first draft of the manuscript with substantial input from J.P. and G.M. All authors contributed to revising the manuscript.

## A Notation used in the main text

**Table A1:**
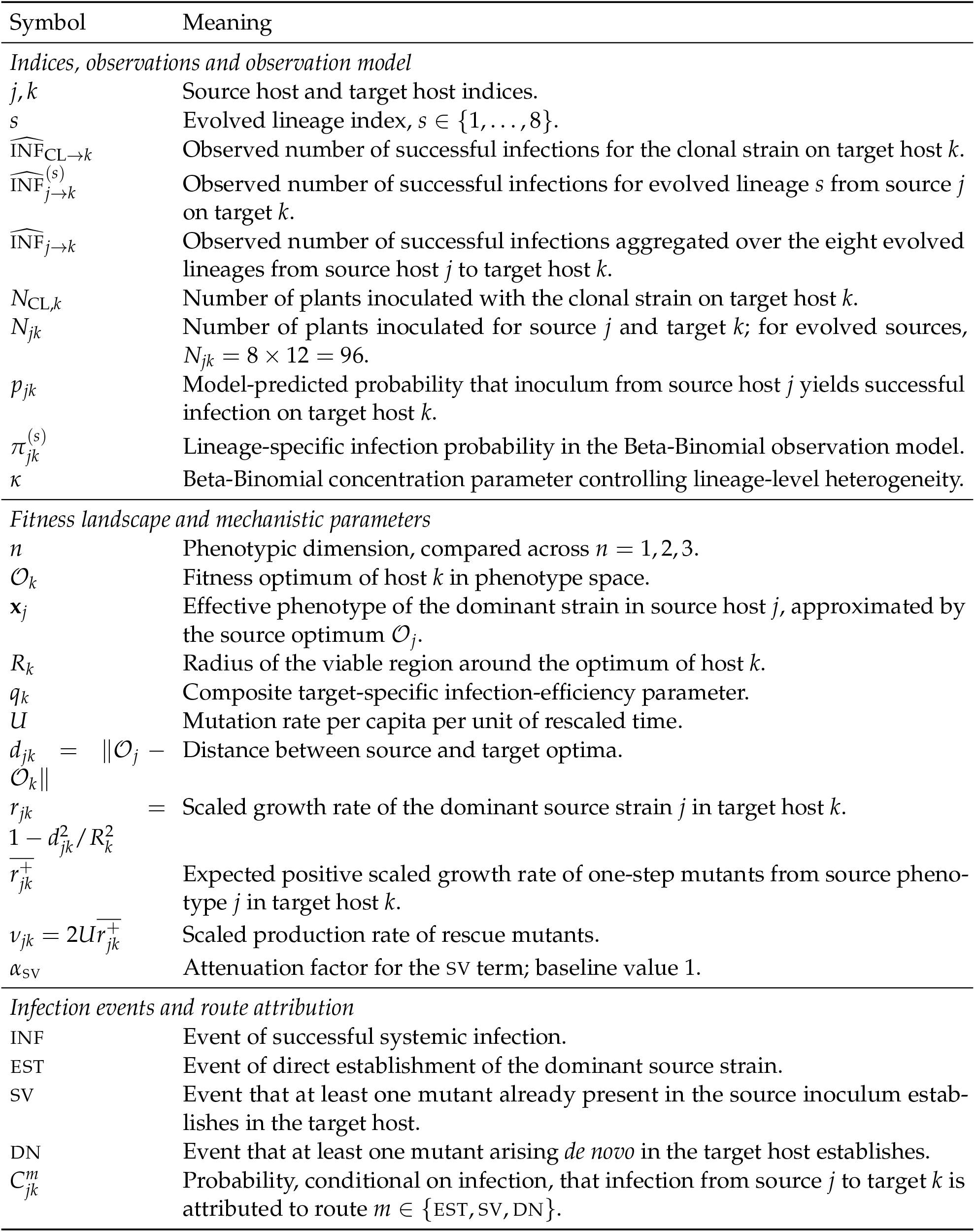
Notation used in the main text.

## B Probability of successful infection under Feller diffusions and a single-step SSWM approximation

We start by detailing the notations and recalling some useful properties of Feller diffusions. Most are provided, with various others and in much more detail, in section 2.3.2 of (Lambert, 2008), except for the Inverse Gaussian result (which was hinted at by Amaury Lambert in 2013). Several of these properties have been used to model evolutionary rescue (ER) in previous work, *e*.*g* (Martin et al., 2013; Gomulkiewicz et al., 2017; Anciaux et al., 2018, 2019; Osmond et al., 2020). We then state all distributional assumptions and definitions needed for the derivation; the corresponding biological discussion is provided in the main text Sections “Source inocula from Step 1: a Luria-Delbrück construction for viral source populations around the source optimum” and “Stochastic demography and establishment during Step 2: a Feller diffusion approximation to early viral infection dynamics“. We derive the three failure probabilities 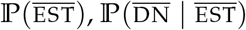,and 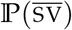, and combine them to obtain the successful-infection probability *p*_*jk*_.

### Notations and useful properties of Feller diffusions

In all our derivations, E_*X*_[*f*(*X*)] is the expectation of any function *f* (*X*) taken over the distribution of a random variable *X*, which may be multivariate or conditional on another variable (*X* | *Y*). We denote the Laplace transform of a positive random variable *X* by *M*_*X*_(*z*) = E[*e*^−*zX*^], *z* ∈ ℝ^+^. Similarly, the probability generating function (PGF) of a discrete nonnegative random variable *X* is denoted by *P*_*X*_(*z*) = E[*z*^*X*^], *z* ∈ [0, 1], and we have *M*_*X*_(*z*) = *P*_*X*_(*e*^−*z*^).

We denote the fact that a process *X*(*t*) follows a Feller diffusion with growth rate *r* and variance coefficient *σ* by:

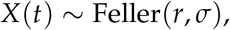

with corresponding stochastic differential equation (SDE):

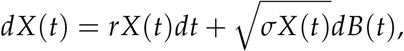

where (*B*(*t*), *t* ≥ 0) is standard Brownian motion. The survival (non-extinction) probability of such a process, started at *X*(0) = 1, is:

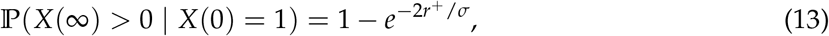

where *r*^+^ = max(*r*, 0).

Consider a supercritical Feller diffusion (*r >* 0) started from *X*(0) = *X*_0_ *>* 0, and fix a target level *X*_*f*_ *> X*_0_. Let

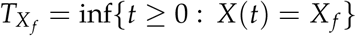

denote the first hitting time of *X*_*f*_. Then the cumulative size accumulated up to that hitting time,

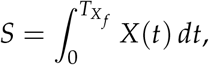

has an Inverse Gaussian distribution:

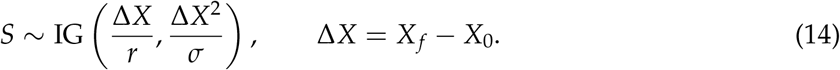

Throughout this appendix, IG(*µ, λ*) denotes the Inverse Gaussian distribution with mean parameter *µ* and shape parameter *λ*, whose Laplace transform is

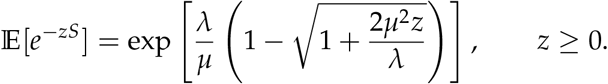

This result was derived in (Martin et al., 2013). Briefly, it stems from the fact that a Feller diffusion *X*(*t*) has Lamperti transforms ℒ(*X*(*t*)), defined below, that is a drifted Brownian motion, denoted BM(*r, σ*):

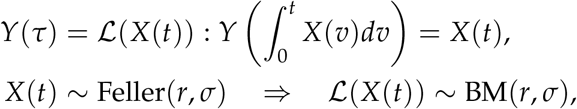

with SDE:

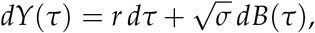

(see definitions 2.2.3.1 p. 92 and section 2.3.2 p. 106 of (Lambert, 2008)). This means that the cumulative size 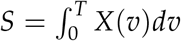 of *X*(*t*) over *t ∈*[0, *T*] is also the first hitting time of the Brownian motion with drift from *Y*(0) = *X*_0_ to *Y*(*S*) = *X*_*f*_. Without loss of generality, we can shift the starting point of the Brownian motion to go from *Y*(0) = 0 to *Y*(*S*) = Δ*X*. We can then use classical results (*e*.*g* (Whitmore and Seshadri, 1987)) on the first hitting time of a drifted Brownian motion started at 0 and hitting Δ*X >* 0 to obtain Eq. (14).

We obtain a similar result for a Feller process doomed to extinction, from some initial size *X*(0) = *X*_0_ *>* 0 up to extinction at some finite time *T*_ext_:

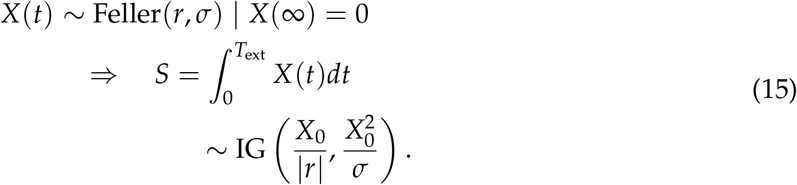

This result applies both to a subcritical process (*r <* 0) and to a supercritical process (*r >* 0) conditioned on ultimate extinction. In the latter case, the conditioned process is a Feller diffusion with growth rate −|*r*| (proposition 2.2.3.1 p. 107 in Lambert, 2008). Under the Lamperti transform, its cumulative size up to extinction is the first hitting time of 0 by a Brownian motion with drift −|*r*|, started at *X*_0_, which gives Eq. (15). The critical case (*r* = 0) is obtained as a limit and corresponds to a Lévy distribution.

#### General assumptions of the model

We first summarize how we conceptualize the whole experimental protocol in terms of demographic and evolutionary model and give some definitions used throughout the derivation. The appendix contains all definitions required for the calculations below; further biological discussion of the assumptions is given in the main text (Section “Mechanistic model“).

In this appendix, we will model the growth process in the source plants that generates the viral population being sampled (end of Step 1). We will also model the process of early growth or decay in target plants (beginning of Step 2) leading to extinction or establishment (successful cross-infection) of the viral population.

For each host *k*, let *ρ*_*ik*_ denote the absolute Malthusian growth rate of clone *i*, and let 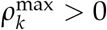 denote the maximal growth rate attainable in that host. Time in host *k* is rescaled by this maximal growth rate: if *τ* denotes absolute time, the rescaled time is 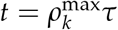. Growth rates in the Feller processes are therefore the host-normalized rates

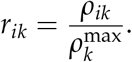

Under the quadratic FGM landscape,

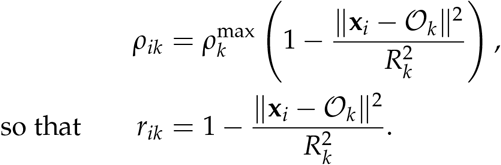

In particular, the optimal phenotype in host *k* has normalized growth rate *r*_*kk*_ = 1. The mutation parameter *U* is defined per capita and per unit of this rescaled time. It is therefore an effective mutation-supply parameter on the host-normalized demographic timescale, rather than a molecular mutation probability per replication.

We use a strong-selection weak-mutation approximation in the specific sense adopted in previous FGM rescue models (Martin and Roques, 2016; Anciaux et al., 2018). Formally, the asymptotic ordering is

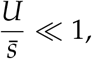

where 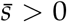 denotes a characteristic magnitude of the selection coefficients of random one-step mutations on the relevant rescaled demographic timescale. The contrasting high-mutation FGM limit, 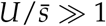, is treated in Anciaux et al. (2019).

This ordering is distinct from requiring the population-wide mutation supply *NU* to be small. A large population may therefore contain several distinct low-frequency mutant lineages. In the calculations below, the weak-mutation-relative-to-selection ordering is implemented by retaining a dominant effective phenotypic background and single-step mutant classes as the leading mutational contribution. Multiple mutant types may be present or arise, but competition and other interactions among mutant lineages are not represented. Sequential accumulation of multiple mutations is also excluded.

The additional assumption that the dominant source phenotype lies near the source-host optimum, **x**_*j*_ ≈ *O*_*j*_, does not follow from SSWM alone. It is a separate assumption concerning adaptation during Step 1 and is discussed in Section “Source inocula from Step 1: a Luria-Delbrück construction for viral source populations around the source optimum“.

For simplicity, clone *j* denotes the dominant effective clone (“wild-type”) in source host *j* at the end of Step 1. Upon entering target host *k*, its normalized growth rate is

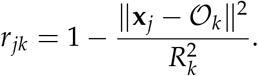

Single-step mutants *j*′ have phenotypes distributed as 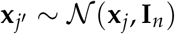 and normalized growth rate 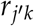 defined by the same fitness function. We use *r*^+^ = max(*r*, 0) and define

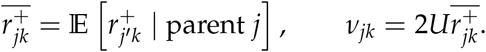

Successful cross-infections from source *j* to target *k* may stem from three (not mutually exclusive) causes:

- EST: establishment of the dominant clone *j* in the target *k*,
- DN: establishment of a mutant *j*′ from clone *j* arising in the target *k*,
- SV: establishment of a mutant *j*′ from clone *j* arising in the source *j*, before sampling.

Let INF denote successful systemic infection and let *p*_*jk*_ = P(INF) be the probability that inoculum from source *j* yields infection in target *k*. Because source standing variation is generated before inoculation, whereas *de novo* rescue depends on the extinction of the dominant clone in the target, we use the factorization

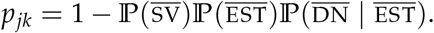

We use Feller diffusions to approximate demographic processes when they involve long timescales, because they approximate discrete branching processes after rescaling of the population size and time (Feller, 1951). This includes the dynamics of all genotypes in the target, as we only wish to describe long-term loss or establishment there. This also includes the growth of the dominant clone *j* in the source *j*, from some initial size *O*(1) to *N*≫ 1 at the time of sampling, where *N* denotes the effective size of the source population available for sampling before the target-entry filter. This *N* is an effective model scale rather than a directly measured within-plant census size. We denote *n*_*ik*_(*t*) the stochastic process describing the number of copies of any clone *i* (clone size) in any host *k*. We write *n*_*ik*_~ Feller(*r*_*ik*_, 1/*ξ*_*k*_) to state that the clone size is approximated by a Feller diffusion with growth rate *r*_*ik*_ and variance coefficient 1/*ξ*_*k*_ in that host species. For target host *k, p*_*k*_ denotes the probability that a virion from the effective source population enters the target-host branching phase. We define the composite parameter

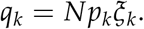

The entry probability *p*_*k*_ is distinct from the overall successful-infection probability *p*_*jk*_.

We distinguish between the status of ‘new’ and ‘mature’ infections, whatever the viral strains or host species considered. A ‘new infection’ is the early infective process of an otherwise uninfected plant: it describes the demographic process at the beginning of Step 2, in the target plants. Stochastic extinction is possible in new infections, for any lineage, even optimal ones. The coefficient *ξ*_*k*_ is the inverse demographic-variance scale in target host *k*, so larger *ξ*_*k*_ corresponds to weaker stochasticity in the Feller approximation. A ‘mature infection’ describes the demographic process in an already fully infected plant, at the end of Step 1, in the source plants. In the viral extension of the Luria-Delbrück model that we use to quantify mutants in that phase (see Appendix B), death is neglected.

This is an effective retention assumption, rather than a claim that biological extinction is impossible. We denote by 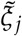 the inverse demographic-variance scale in mature infections in source plants of species *j*. In these mature infections, time is rescaled by the maximal growth in this context, and clone *j*, being near optimal in that host has growth rate 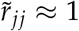 1, in these rescaled time units. The same is true (*r*_*jj*_ ≈ 1) for clone *j* in new infections when the target is of the same species as the source (*k* = *j*). We assume 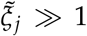, so that stochastic loss after appearance is neglected in the source-growth approximation.

There is one class of processes that we do not model by a Feller diffusion: the clone size of mutants *j*′ in the source plant. These mutant clone sizes grow over short timescales (before being sampled) up to a few copies. Their distribution remains *a priori* discrete and there is no natural rescaling under which the diffusion approximation emerges. Rather, we use a Luria-Delbrück type of model that we accommodate to the specificities of viral demography by using the simplest tractable stochastic model of viral demography: the pure burst model (Hubbarde et al., 2007). This model has two parameters: a burst size *B* and a rate of bursting *λ*. We write 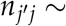 PBM 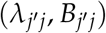 to state that a clone size process is approximated by a pure burst model in that case, for any mutant clone *j*′ of strain *j*, in source *j*. This model is detailed in Section “Luria-Delbrück model under the ‘pure burst model’“.

Below is a summary of the various stochastic processes considered here, in the source *j* and target *k* for the dominant clone *j* and its mutants *j*′:

- wild type *j* in mature infections in source *j*: *n*_*jj*_ ~ Feller(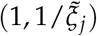) with 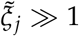,
- wild type *j* in new infections in target *k*: *n*_*jk*_ ~ Feller(*r*_*jk*_, 1/*ξ*_*k*_) with *ξ*_*k*_ ≪ 1,
- mutant *j*′ in mature infections in source 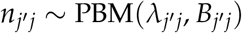 with 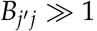,
- mutant *j*′ in new infections in target *k*: 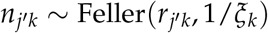 with *ξ*_*k*_ ≪ 1.

#### Fate of the dominant clone *j* **from source host** *j* **to target host** *k*

In this section, we describe the properties of the dominant clone *j* over the course of the experiment, from the late phase of Step 1 (up to inoculation) to the end of Step 2 (extinction or establishment).

##### Step 1

Clone *j* grows from a small initial size *n*_0_ = *O*(1) to some large final size (*O*(*N*)) at the time *T* of sampling. By this time, the population consists of *N*(1 − *F*) virions of type *j* plus a stochastic but small fraction (*F* ≪ 1) of single-step mutants from clone *j*, under the dominant-background, low-frequency-mutant approximation. As *N* ≫ 1, the rescaled process *x*_*jj*_(*t*) = *n*_*jj*_(*t*)/*N* is a rescaled Feller diffusion: *x*_*jj*_(*t*) ~ Feller 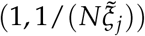. This process grows from *x*_*jj*_(0) = *n*_0_/*N* to *x*_*jj*_(*T*) = *N*(1 − *F*)/*N*≈ 1, by the time *T* of sampling. Hence, it grows over *t* ∈ [0, *T*] by some increment Δ*x*_*jj*_ = 1− *n*_0_/*N* ≈1. The (unscaled) cumulative size of clone *j* is thus, by Eq. (14) and scaling properties of the Inverse Gaussian:

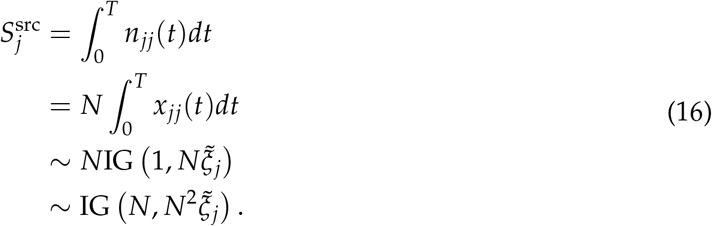

Note that here, we use the assumption in Section “Source inocula from Step 1: a Luria-Delbrück construction for viral source populations around the source optimum“, that in mature infections (the source plant), stochastic extinction can be neglected. This means we do not need to condition the Feller process *n*_*jj*_(*t*) on having avoided extinction initially when it started in *n*_0_ = *O*(1) copies.

##### Step 2

The inoculation process consists in three phases: sampling from the source host, then entry into the plant system of the target host, then early (branching) growth in that host. We ignore differences in the average number of virions sampled from different source hosts (see Section “Source inocula from Step 1: a Luria-Delbrück construction for viral source populations around the source optimum“). Each virion from the source population then has probability *p*_*k*_ to enter the plant system, in species *k*. This gathers the probability to be sampled in the source and then to successfully infect a first cell in the target. Assuming *N* ≫ 1 while *p*_*k*_ ≪ 1, a Poisson limit of the binomial sampling arises. A number *V*_*jk*_ ~ Poisson(*N*(1 − *F*)*p*_*k*_) of virions of type *j* enter host *k* to initiate infections. Because *F* ≪ 1, we use 1 − *F* ≈ 1, so that:

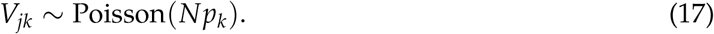

Clone *j* is now present at abundance *n*_*jk*_(0) = *V*_*jk*_ in host *k* (we reset time upon entry). Each of its copies may establish or be lost independently (by the branching property during early infection). The probability that none of them establishes is (using Eq. (13)):

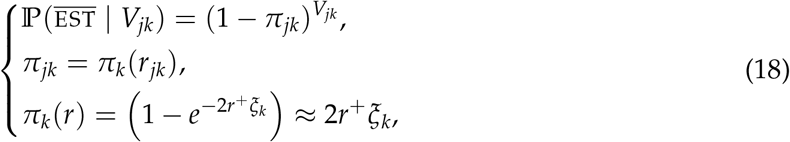

with *r*^+^ = max(*r*, 0). Deconditioning on the distribution of *V*_*jk*_ gives the probability:

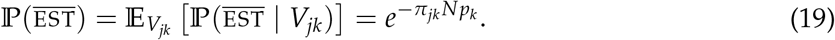

Using the linear approximation for establishment probability in Eq. (18), then yields:

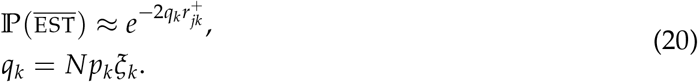

It is this expression that is plugged into Eq. (3) to compute *p*_*jk*_ in Eq. (6). The linearized approximation for *π*_*k*_(*r*) in Eq. (18) reduces the number of parameters in the statistical inference ({*N, ξ*_*k*_, *p*_*k*_} ↦*q*_*k*_), but requires *ξ*_*k*_ ≪ 1 in all hosts, so that it applies to ‘re-infection’ controls (*k* = *j, r*_*jk*_ = *r*_*jj*_ = 1). We note that the estimated *q*_*k*_ are of order 1 in all hosts: as we expect that a substantial number of viruses do enter the plant system (*Np*_*k*_ ≫ 1) these estimates are consistent with the assumption *ξ*_*k*_ ≪ 1.

Conditional on not establishing (given 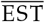), clone *j* must decay to extinction. By Poisson thinning, the numbers of establishing and ultimately doomed founding copies are independent Poisson random variables. Consequently, conditional on 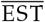, the number of doomed founders is

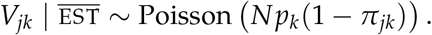

Below, *V*_*jk*_ denotes this conditional number of doomed founders. We need the cumulative size of this doomed lineage because, in Section “Fate of mutants arising *de novo* in the target host: 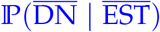“, the number of *de novo* mutants produced before extinction is proportional to the total amount of parental replication in the target host. The clone size process then goes from *n*_*jk*_(0) = *V*_*jk*_ to *n*_*jk*_(*T*) = 0 over some finite time *T*. We can apply a similar scaling argument as in Eq. (16) to obtain a Feller approximation (*n*_*jk*_ ↦ *x*_*jk*_ = *n*_*jk*_/*V*_*jk*_). We have *x*_*jk*_(*t*) | *x*_*jk*_(*T*) = 0 ~ Feller(−|*r*_*jk*_|, 1/(*V*_*jk*_*ξ*_*k*_)) and the (unscaled) cumulative size of the clone, up to extinction is (by Eq. (15)):

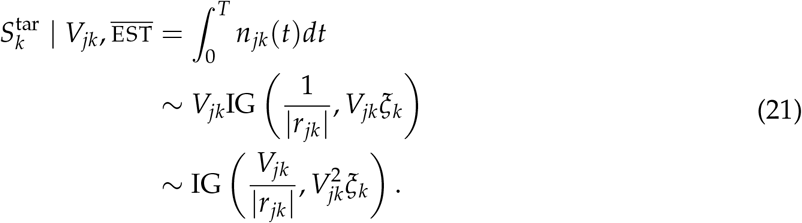

#### Mutation process

In any host *k* at any time *t* the rate of production of new mutants is assumed to be proportional to (i) a mutation probability *u*∈ [0, 1] per RNA copy, (ii) a viral equivalent of birth rate *b*_*jk*_ for strain *j* in host *k* (rate of production of new RNA copies per virus particle per arbitrary rescaled unit time) and (iii) the stochastic size *n*_*jk*_(*t*) of a strain *j* clone in a host *k*. We assume that *u* is an inherent property of the viral species and thus independent of the host, strain or time. We assume that the viral birth rate *β*_*jk*_, per arbitrary unit time, scales with the maximal growth rate in host *k* (in the same unit time): 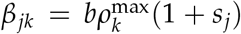 with some constant *b* and some *relative* effect *s*_*j*_ of genotype *j*, independent of the host. Consistently with the time rescaling defined in Section “General assumptions of the model“, the viral birth rate per unit of rescaled time is 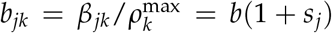 which is independent of the host species. Finally, we assume that relative mutation effects on birth rates are small *s*_*j*_ = *O*(*ϵ*) and that the mutation probability per viral copy is also small (*u* = *O*(*ϵ*)), for some *ϵ*≪ 1. We then get, to leading order, a constant mutation rate across hosts and viral genotypes:

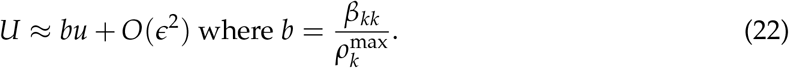

Notice that the factor *b* is a ratio of birth rate over growth rate: it may be substantially larger than one if there is a large demographic turnover (high birth and death rates of similar order). Therefore, even if the mutation probability per copy *u* is small, we may get a substantial *U* in the relevant timescale.

#### Fate of mutants arising *de novo* in the target host: 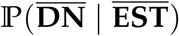

We consider only *de novo* mutations in those cases where the parent clone *j* ultimately goes extinct in the target host, at some finite time *T*. In that case, there is a finite production of mutants by that clone up to its extinction, and we wish to compute the probability 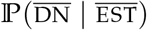 that none of these mutants manage to establish and rescue the population. We first derive the process for an arbitrary mutant type *j*′ produced by clone *j* and we denote 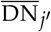 the event that no mutant of this type yields a rescue.

Mutants of type *j*′ arise in single copy according to a Poisson process with intensity 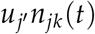 where 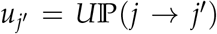 is the rate of mutations to type *j*′ from the dominant type *j*, and *U* is the genomic mutation rate, per rescaled time per capita, detailed in Section “Mutation process”. Conditional on appearing, this single copy produces a whole ‘lineage’, that may establish with probability *π*_*j′k*_ = *π*_*k*_(*π*_*j′k*_ given by Eq. (18). By the thinning property of Poisson processes, the number 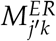 of such rescue mutant lineages, produced until time *T*, is:

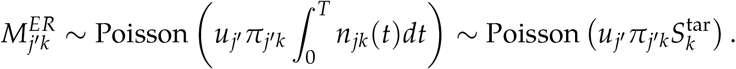

The probability that no successful rescue lineage of type *j*′ arises is therefore:

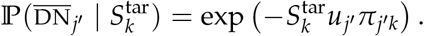

Appearance and establishment of mutants of different types are independent so the probability that no mutant of any type produces an ER is the product:

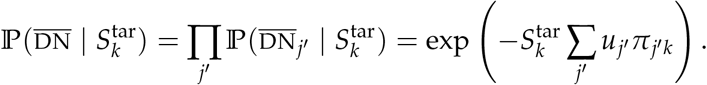

Taking the continuous types limit we have: 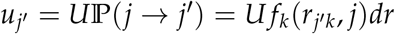 where *f*_*k*_(*r, j*) is the probability density function (pdf) of the growth rates in host *k*, among random mutants from clone *j*. Using the establishment probability function *π*_*k*_(*r*) in Eq. (18) yields:

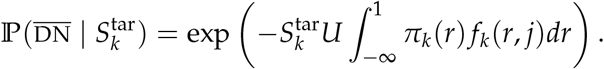

We then use the linearized approximation for *π*_*k*_(*r*) in Eq. (18) which gives:

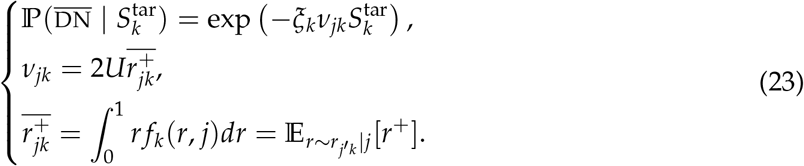

The term 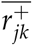 is the expected positive part of the scaled growth rate in host *k* among random mutants from clone *j*. Taking successive expectations over (i) the distribution of 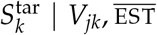, EST in Eq. (21), then (ii) the conditional distribution of 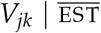 given above, yields the probability that no *de novo* rescue arises:

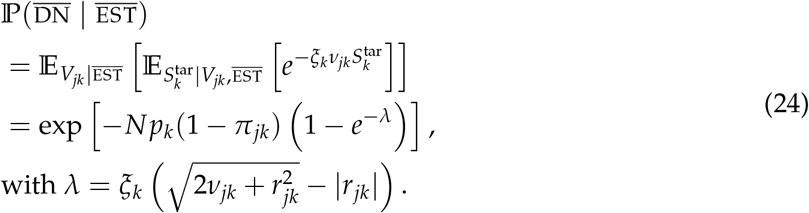

The linear approximation used in Section “Fate of the dominant clone *j* from source host *j* to target host *k*” requires *ξ*_*k*_ ≪ 1. Hence *π*_*jk*_ = *O*(*ξ*_*k*_) and *λ* = *O*(*ξ*_*k*_), so that

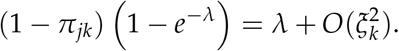

Since *Np*_*k*_ = *q*_*k*_/*ξ*_*k*_, the corresponding correction in the exponent is *O*(*q*_*k*_*ξ*_*k*_) and vanishes in the small *ξ*_*k*_ limit with *q*_*k*_ fixed. This yields the simpler first-order formula:

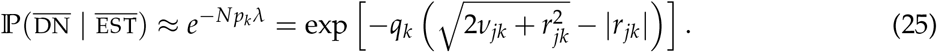

with 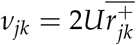. This expression was plugged into Eq. (3) to compute *p*_*jk*_ in Eq. (6). It is continuous with respect to clone *j*’s growth rate over its full possible range: *r*_*jk*_ ∈ [−∞, 1].

#### Fate of mutants arising in the source host: P(SV)

Next, we must consider mutants that may be segregating in the sampled inoculum, to compute 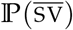. Similarly to Section “Fate of mutants arising *de novo* in the target host: 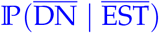”, we denote 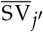 the event that no segregating mutant of type *j*′ establishes in the target host.

This time, we must consider the appearance of mutants in the source host, then their fate (establishment or loss) in the target host. A mutant of type *j*′ from the dominant clone *j* segregates at some random frequency 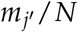 in the source population of size *N*, at the time of sampling. The total number of virions that enter the plant system is *V* ~ Poisson(*Np*_*k*_). Among these, a number 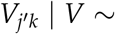 Binomial(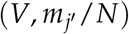 are of type *j*′. The number of mutants *j*′, from the source, that enter the target, is: By Poisson thinning, conditional on 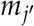,

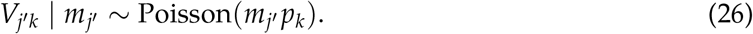

Each mutant copy establishes with probability 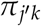 given by Eq. (18), so the probability that no such establishment occurs is

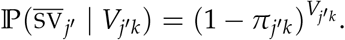

Deconditioning on 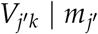 yields:

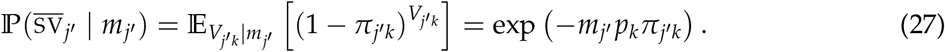

It now remains to describe the distribution of 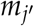 produced during the growth of clone *j* in the source. If no such lineage appeared at any time before sampling, then 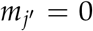. Otherwise, the final size of clone *j*′ follows some random discrete distribution D, independent of type *j*′ and of the cumulative size 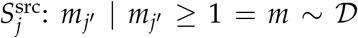. We justify these assumptions and detail the distribution D in the next section.

At this point, we use the baseline source-growth approximation stated above: once a mutant lineage is produced in the source, its stochastic loss before sampling is neglected. Thus, 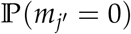 is approximated by the probability that no mutation event of type *j*′ occurred during source growth. By a similar argument as in Section “Fate of mutants arising *de novo* in the target host: 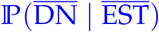, 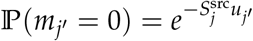, where 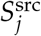 is the cumulative size of clone *j* over its growth in the source plant from a low initial number. This approximation is relaxed in the sensitivity analysis of Supplementary File 1, Section S3. The stochastic representation of the mutant clone size, in the source, is therefore:

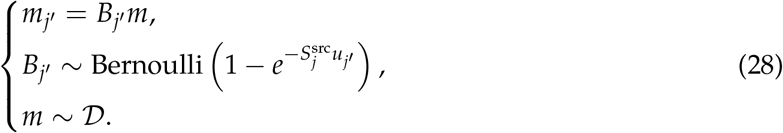

Taking the expectation of 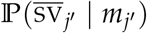 in Eq. (27) over the distributions of 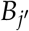 yields:

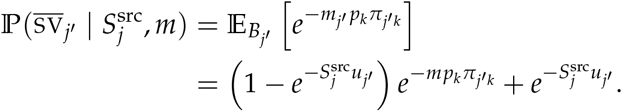

We can now take the expectation over 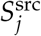, whose distribution is given by Eq. (16), yielding the expression:

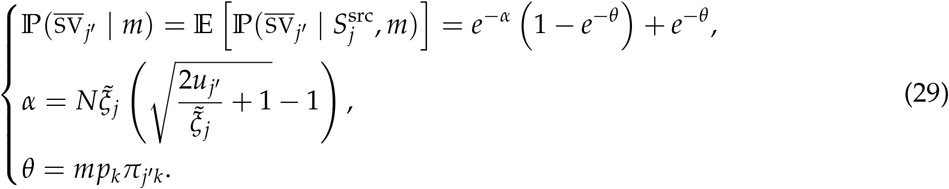

From there, we take the continuous types limit, as in Section Fate of mutants arising *de novo* inthe target host: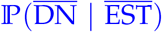)“, by setting 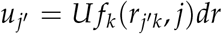 and letting *dr* → 0, so that 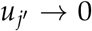. Equivalently we could take a limit as 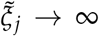. Approximating log 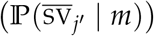 in Eq. (29) to leading order in 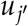 greatly simplifies the expression, yielding:

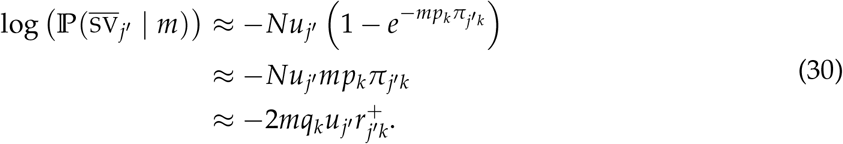

The first approximation is the leading-order expansion in 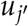 used for the continuous-type limit. The second approximation uses 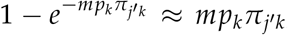, and the last line uses the same weak-establishment linearization as above,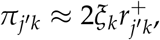 with 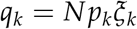.

Using this expression, we can compute the probability across all possible types, in the continuous types limit. There is no establishment from segregating mutants overall *iff* none of the types produce an ER. Each type has independent fate and the mutant clone sizes from each type *j*′ are i.i.d. draws: *m*~ *D*. Overall, taking the product of type-specific probabilities 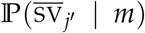 and using the approximation in Eq. (30), the sum over mutant types becomes, in the continuous-type limit,

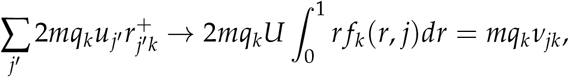

With 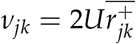. This yields:

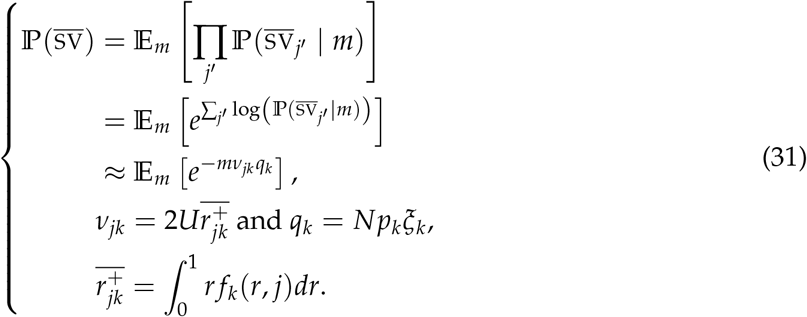

#### Luria-Delbrück model under the ‘pure burst model’

Eq. (31) states that, given the Laplace transform of the distribution of mutant clone sizes, *M*_*m*_(*z*) = E[*e*^−*zm*^], *z* ∈ ℝ^+^, the probability that no rescue comes from standing variation is: 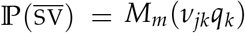. It now remains to compute this Laplace transform. To do so, we use the stochastic pure burst model proposed in Eq. (6) of (Hubbarde et al., 2007). This continuous time branching process is an extension of the classic pure birth model to allow for more than two offspring upon birth. The duration of an infectious cycle (which includes transport to infect new cells) is exponentially distributed with rate *λ*. Upon completing its infectious cycle, the virus then infects 1 + *B* new host cells where *B* ≥ 1 and the infection goes on. This model has no extinction but yields a stochastic demography. It thus corresponds to the conditions we have assumed for mature infections in source hosts (no stochastic loss), while still allowing analytical treatment.

We denote *m*_*t*_ ~ PBM(*λ, B*) (for ‘Pure Burst Model’) the process *m*_*t*_ with parameters *λ* and *B*, started in *m*_0_ = 1 copy at time *t* = 0. The PGF *P*(*z, t*) of *m*_*t*_ is (Hubbarde et al., 2007):

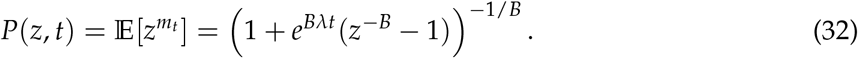

The expected process grows exponentially as E[*m*_*t*_] = *∂*_*z*_*P*(*z, t*) |_*z*=1_ = *e*^*rt*^ where *r* = *λB*. We rescale time and thus *λ* so that *r* = 1, hence *λ* = 1/*B*. The corresponding Laplace transform of the mutant clone size, in rescaled time, is:

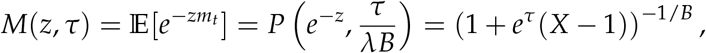

where *X* = *e*^*zB*^. From classic Luria-Delbrück theory, conditional on sampling a mutant lineage at the end of an exponentially growing parental population, the backward time to the mutation event has an exponential density. More precisely, if the parental population grows at rate *r*, the age of a randomly sampled mutant clone is *a* ~ Exp(*r*). This exponential age distribution is conditional on the no-extinction pure-burst source model used here for mutant lineages; allowing mutant death would preferentially remove older lineages and shift the distribution of surviving mutant ages toward more recent origins. This statement concerns the time at which the mutation was produced by the parent; any type-specific growth cost is then included in the subsequent growth of the mutant clone. This is the approximation used to compute the Laplace transform of *m*_*a*_ by averaging *M*(*z, τ*) over *τ* = *a*.

Assume that the mutant has some cost 0 ≤ *c <* 1, so that the mutant growth rate is 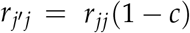. Then, in our rescaled time, the parent growth rate is *r* = 1/(1− *c*) so the ages of mutant lineages are *a* ~ Exp(1/(1− *c*)). Taking the expectation yields the Laplace transform of mutant clone sizes:

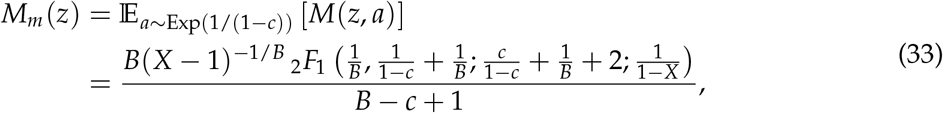

where_2_*F*_1_(.,.;.;.) is the Gaussian hypergeometric function.

As a sanity check, the model recovers the Lea-Coulson result for binary division, corresponding to *B* = 1, in the absence of a mutant cost (*c* → 0). Equation (33) then becomes

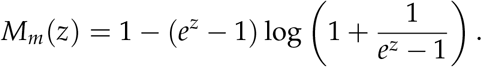

Equivalently, with *X* = *e*^*z*^,

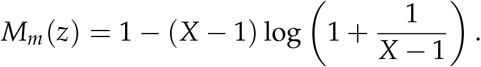

This is the Laplace transform of the classical Lea-Coulson clone-size distribution

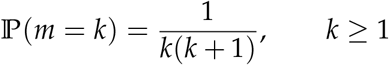

(Lea and Coulson, 1949).

From there, let us assume that the virus has a substantial burst size *B*≫ 1 and that *z* is non vanishing. Then we have *X* = *e*^*zB*^≫ 1. Taking a leading order of *M*_*m*_(*z*) in Eq. (33) as *X* → ∞ yields:

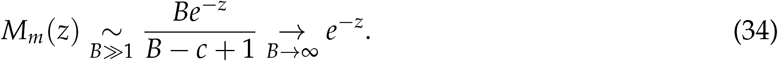

Therefore, when the burst size of the viruses is large, the Laplace transform of the mutant clone sizes converges to *M*_*m*_(*z*) = *e*^−*z*^ independently of their cost *c*. This confirms the assumption made in Section “Fate of mutants arising in the source host: P(SV)” that the distribution of *m* is independent of the mutant type *j*′. Numerical comparison with the exact expression in Eq. (33) shows that the approximation *M*_*m*_(*z*) ≈ *e*^−*z*^ is very accurate even for a mild *B* = 5 or *B* = 10 and across costs ranging from *c* = 0.001 to *c* = 0.5.

We thus insert this limiting Laplace transform in Eq. (31) to compute the probability that no mutant from the source host produces an ER:

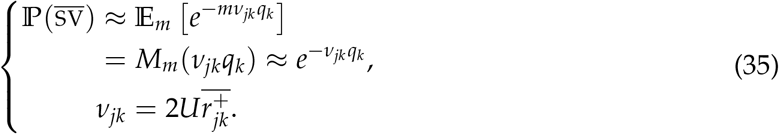

Combining Eqs. (20), (25) and (35) gives

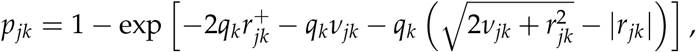

which is the successful-infection probability used in the likelihood.

## C Rescue probability in the FGM: complementary approximations and numerical validation

This appendix derives the closed-form approximation of 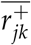, the expected positive part of the scaled growth rate in host *k* among random single-step mutants (*j*′) derived from parent strain *j*, and assesses its numerical accuracy against the exact noncentral-*χ*^2^ expression.

### Gaussian moment matching *G*_*jk*_

Recall that for a virus evolved on host *j* and inoculated on host *k*,

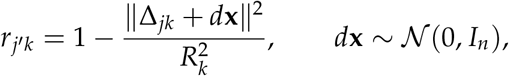

where Δ_*jk*_ = *O*_*j*_ − *O*_*k*_ and *d*_*jk*_ = ∥Δ_*jk*_∥. Let

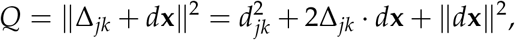

so that

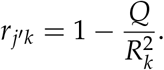

**Mean**. Since E[Δ_*jk*_ · *d***x**] = 0 and E[∥*d***x**∥^2^] = *n*,

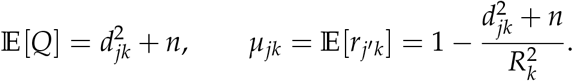

**Variance**. Write 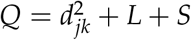 with

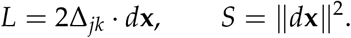

Because 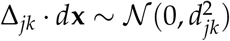,

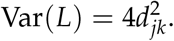

Since 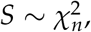,

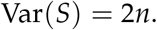

Moreover, Cov(*L, S*) = 0 by symmetry. Hence

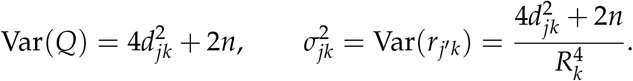

We then approximate

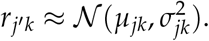

Therefore 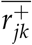 is approximated by the truncated first moment above zero. For a normal random variable *r* ~ *N* (*µ, σ*^2^),

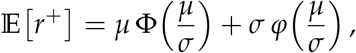

where Φ and *φ* denote the cumulative distribution function and density of the standard normal distribution. This yields the Gaussian approximation:

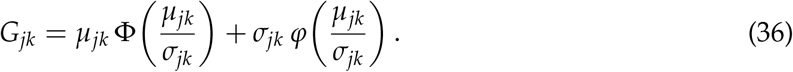

#### Laplace approximation *L*_*jk*_

For completeness, we sketch the derivation of the Laplace approximation used in the main text. A closely related derivation is given in Anciaux et al. (2018).

Recall that

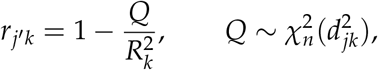

so that

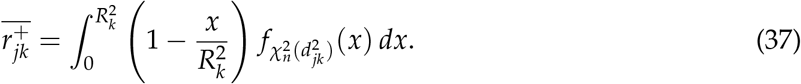

We then use the Bessel-function expression of the noncentral-*χ*^2^ density,

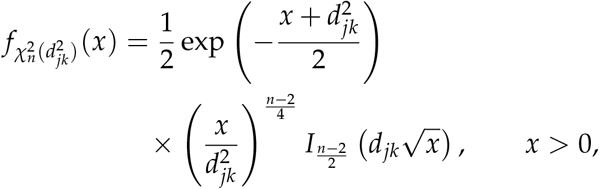

where *I*_*ν*_denotes the modified Bessel function of the first kind. If *R*_*k*_ is fixed and *d*_*jk*_ − *R*_*k*_ is large, then for any fixed *A >* 0 the contribution of the region 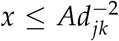 in (37) is negligible compared with the full integral. Hence the dominant contribution comes from values of for which the Bessel argument 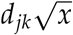 is large. For fixed order *ν*and positive real argument, its standard large-argument asymptotic expansion is

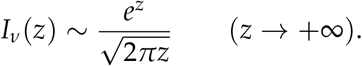

Using this asymptotic, we obtain

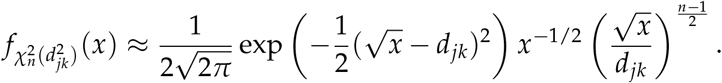

Writing *x* = (*R*_*k*_ − *t*)^2^, we obtain

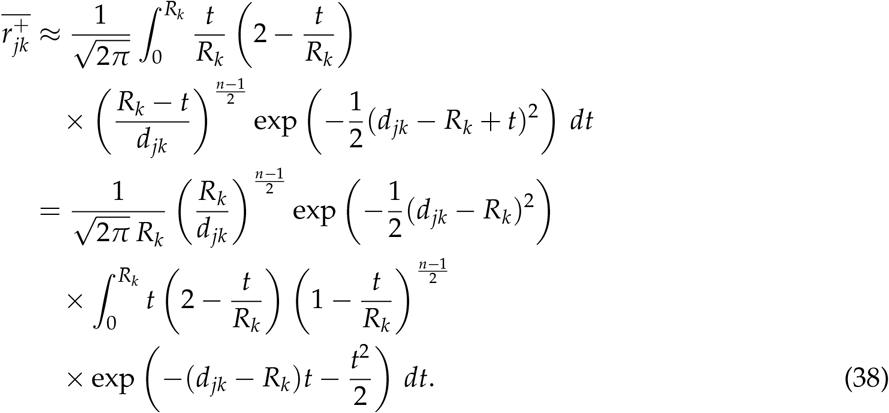

The Laplace approximation is now obtained in the same regime where *d*_*jk*_ − *R*_*k*_ is moderately large, so that the exponential factor localizes the integral near *t* = 0. We then replace the slowly varying prefactor by its first-order exponential expansion at *t* = 0 and extend the upper limit of integration to infinity. More precisely, as *t* → 0,

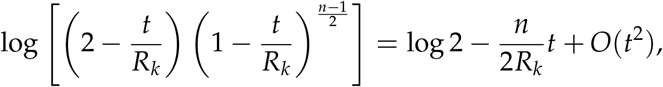

so that

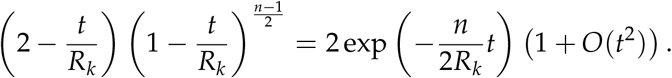

Substituting this local approximation gives

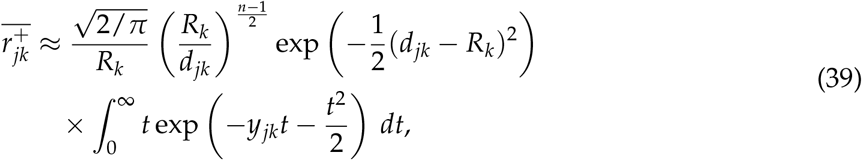

where

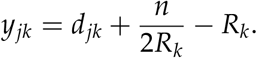

The remaining integral is explicit. Indeed,

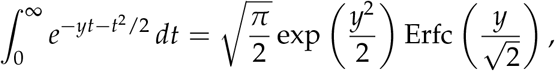

so differentiation with respect to *y* yields

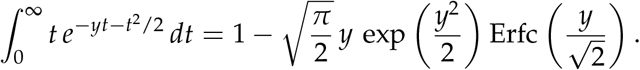

Therefore,

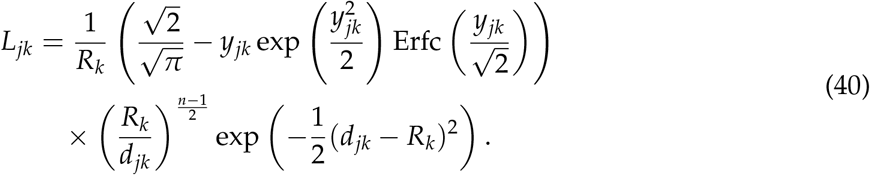

#### Numerical validation against the exact expression

We numerically compared the exact quantity 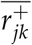 in (4) to the Gaussian and Laplace approximations *G*_*jk*_ and *L*_*jk*_ defined in (36), as well as to the matching approximation min *G*_*jk*_, *L*_*jk*_, for *n* = 1, 2, 3 and representative values *R*_*k*_ ∈{1.5, 5, 10} (Figure C1). In each panel, the exact curve was evaluated numerically from the noncentral-*χ*^2^ expression, and the dashed vertical line marks the threshold *d* = *R*_*k*_.

The two approximations are complementary. The Gaussian approximation *G*_*jk*_ is accurate for *d < R*_*k*_ but overestimates the tail when *d > R*_*k*_, whereas the Laplace approximation *L*_*jk*_ captures well the decay beyond *d* = *R*_*k*_ but is less accurate for small distances. Across all cases shown in Figure C1, the matching approximation min(*G*_*jk*_, *L*_*j*_) remains very close to the exact 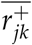 over the whole range of distances, including the transition region around *d* = *R*_*k*_.

## D Individual-based validation of branching-process approximations

The analytical model uses two distinct branching-process approximations at successive stages of the experimental protocol. During Step 1, mutations arise while the viral population grows in the source host, and the source-standing-variation calculation depends on the distribution of mutant clone sizes at the time of sampling. During Step 2, the early dynamics of each lineage in the target host are approximated by a Feller diffusion. We assessed both approximations using individual-based simulations of continuous-time burst-death processes.

Throughout this section, *B* ≥ 1 denotes the net increase in population size produced by one burst event. Thus, one particle is replaced by *B* + 1 particles and the population size changes from *n* to *n* + *B*. Binary division therefore corresponds to *B* = 1. The parameter *λ* is the per-capita burst-event rate, consistently with the pure-burst model of Appendix B, and *δ* is the per-capita death rate.

**Figure C1:**
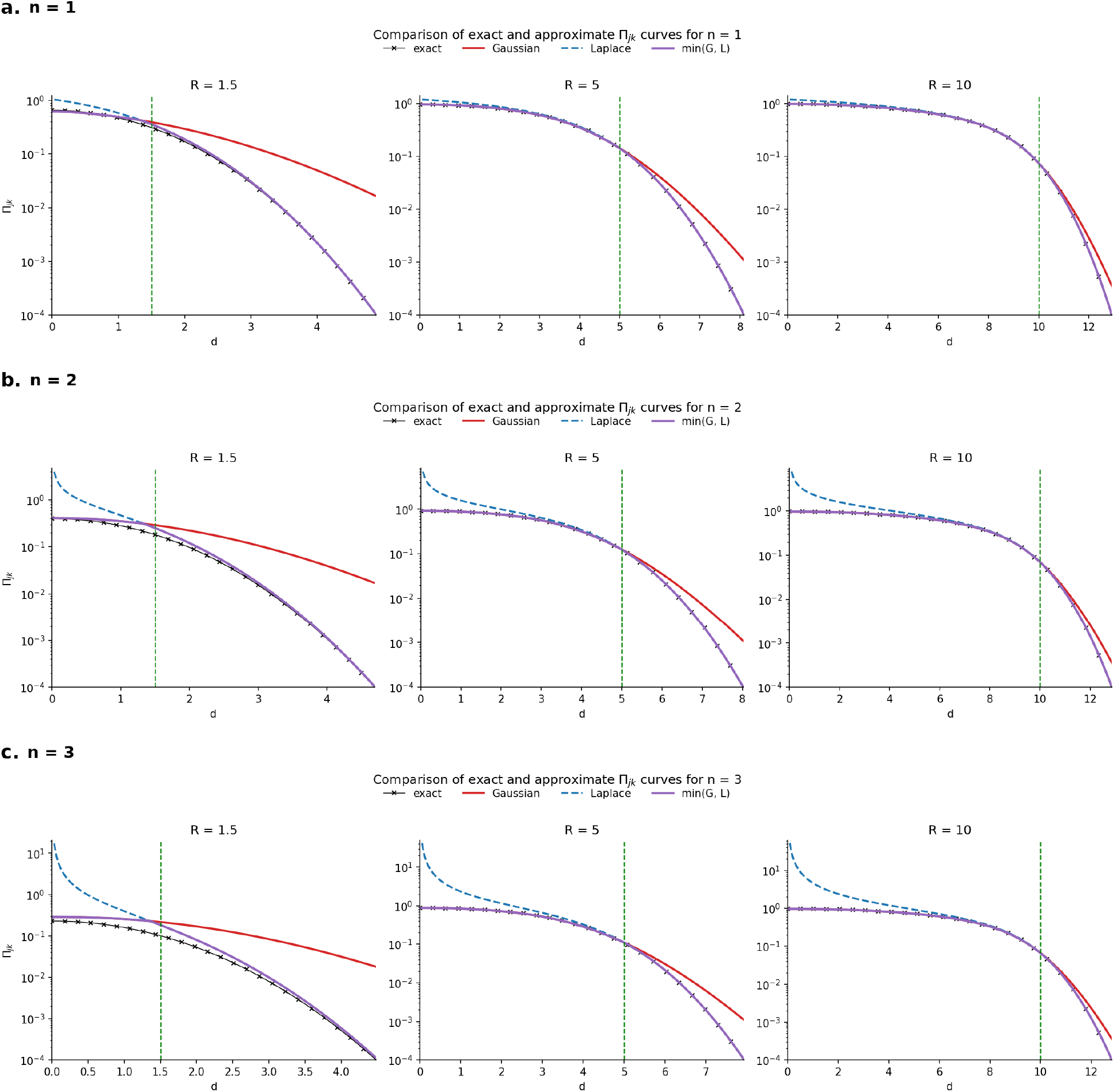
Numerical validation of the closed-form rescue approximation for *n* = 1, 2, 3. The green dashed lines correspond to *d* = *R*.

### Step 1: distribution of source-mutant clone sizes

The source-standing-variation calculation depends on the Laplace transform of the size *m* of a mutant clone sampled from the source population:

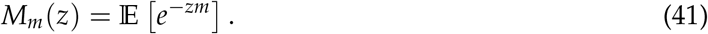

We simulated a parental type and a mutant type using an exact continuous-time Markov process. A particle of type *i* underwent a burst event at rate *λ*_*i*_ or a death event at rate *δ*_*i*_. A mutant burst produced only mutant offspring, with no back mutation. At each parental burst, the parental particle was replaced by *B* + 1 offspring, each of which independently became a mutant with probability *u*. The number of mutants generated during a parental burst was therefore distributed as Binomial (*B* + 1, *u*).

Each simulation started with ten parental particles and no mutants. It was stopped when the total population first reached or exceeded *N* = 103. We used the microscopic mutation probability *u* = 10^−4^, giving *Nu* = 0.1. Only replicates with a positive mutant count at sampling were used to calculate the empirical transform in Eq. (41). The choice *Nu* = 0.1 places the simulations in a rare-mutation regime while providing enough positive-mutant replicates to estimate *M*_*m*_(*z*).

The pure-burst calculation in Appendix B gives the exact analytical expression in Eq. (33). For a large net burst increment, this expression has the limiting form

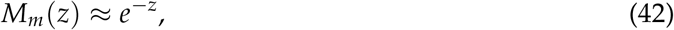

which leads to the approximation used in Eq. (35):

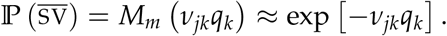

We examined three simulation settings.

#### Binary-division benchmark

We first considered ordinary binary division, corresponding to *B* = 1, with equal parental and mutant burst rates

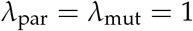

and no death:

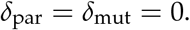

The mutant had no growth-rate cost. In this case, the model reduces to the classical setting underlying the Lea-Coulson mutant clone-size distribution (Lea and Coulson, 1949),

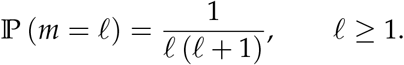

Its Laplace transform is

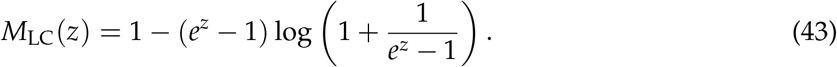

Among 7,000 simulated populations, 1,231 contained at least one mutant at sampling. Figure D1a shows that the empirical transform calculated from these positive-mutant populations coincides with both the classical Lea-Coulson result and the exact pure-burst expression evaluated at *B* = 1. This provides a benchmark for the individual-based simulation algorithm and the analytical calculation.

#### Viral burst demography

We next considered a net burst increment *B* = 20, corresponding to one particle being replaced by 21 particles, with

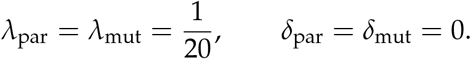

Both types therefore had growth rate

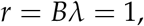

and the mutant had no cost. Among 70,000 simulated populations, 6,967 contained mutants at sampling. Figure D1b shows close agreement between the empirical transform and the exact pure-burst expression. It also shows that, already at *B* = 20, the large-burst approximation

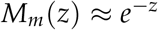

is accurate over the displayed range of *z*. In contrast, the classical binary-division Lea-Coulson transform differs markedly from the viral large-burst result.

#### Robustness to mutant cost and death

The analytical expression in Appendix B assumes a pure-burst process without particle death. We therefore considered an additional case outside this assumption. We used *B* = 50, corresponding to one particle being replaced by 51 particles, with

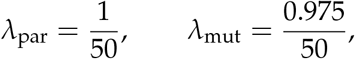

and a common death rate

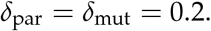

The resulting net growth rates were

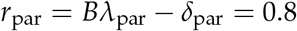

and

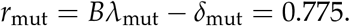

The relative mutant cost, defined from the net growth rates, was therefore

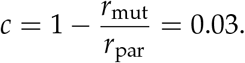

The death rate was equal to 20% of the parental burst-driven growth scale *Bλ*_par_ = 1. Among 70,000 simulated populations, 4,206 contained mutants at sampling.

Figure D1c shows that the empirical transform remains close to the analytical pure-burst expression evaluated at the corresponding relative cost, despite the addition of death to the individual-based process. It also remains close to the large-burst limit *e*^−*z*^. Thus, the approximation used for the SV term is not restricted to the exact death-free simulation setting and remains accurate in this example with moderate death and a mutant cost of approximately 3.1%.

**Figure D1:**
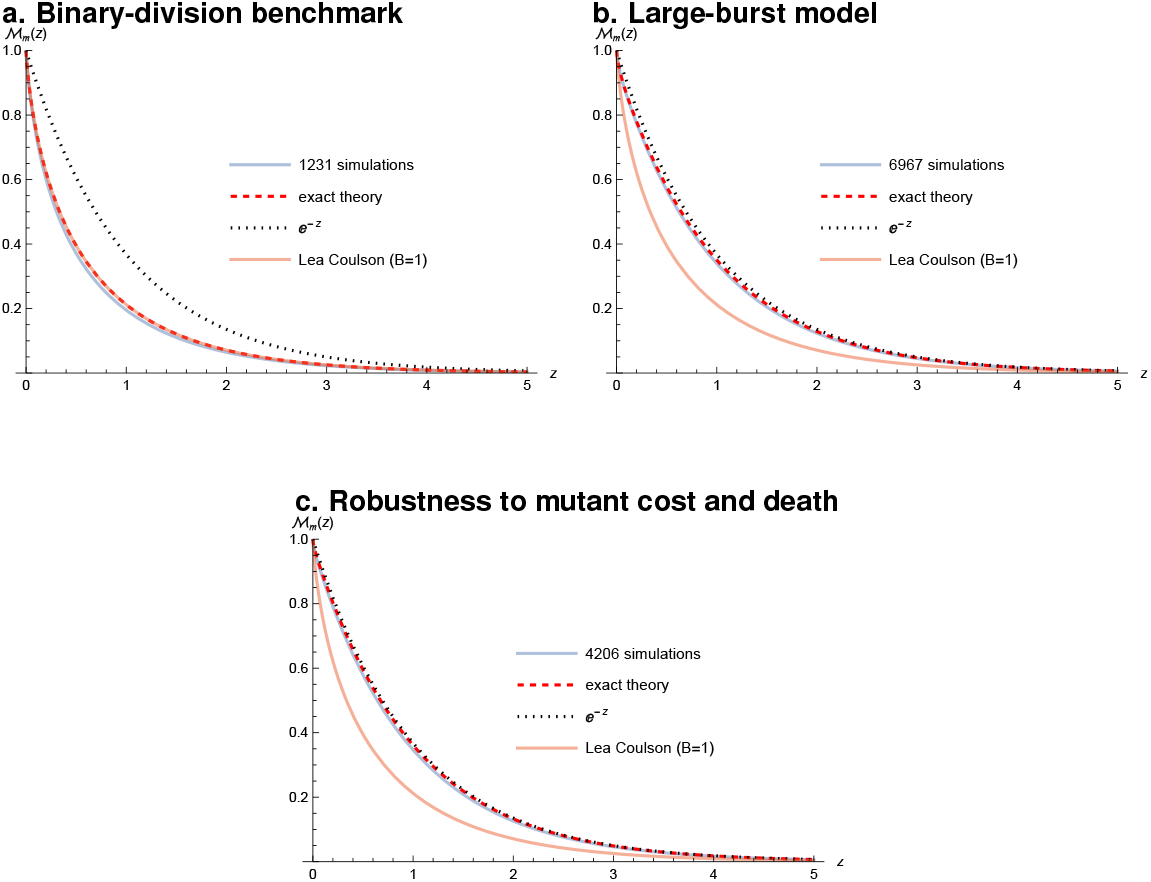
Individual-based validation of the source-mutant clone-size Laplace transform *M*_*m*_(*z*) = E[*e*^−*zm*^]. The empirical curves were calculated from simulated populations containing at least one mutant. The other curves show the exact pure-burst expression from Eq. (33), its large-burst limit *e*^−*z*^, and the classical binary-division Lea-Coulson transform in Eq. (43). All simulations started with ten parental particles, used the microscopic mutation probability *u* = 10^−4^, and stopped when the total population first reached or exceeded *N* = 10^3^, giving *Nu* = 0.1. (a) Binary division, with *B* = 1, no mutant cost, and no death. The empirical transform coincides with the exact pure-burst and Lea-Coulson expressions. (b) Net burst increment *B* = 20, corresponding to 21 particles after a burst, with no mutant cost and no death. The empirical transform agrees with the exact pure-burst expression and is already close to *e*^−*z*^. (c) Net burst increment *B* = 50, corresponding to 51 particles after a burst, with death rate *δ* = 0.2 and relative mutant cost *c* = 0.03. The empirical transform remains close to the death-free analytical expression and to *e*^−*z*^.

### Step 2: approximation of target-host burst-death dynamics by a Feller diffusion

We next tested the diffusion approximation used for the early dynamics of lineages in the target host. Consider a population of size *n*(*t*) in which each particle undergoes a burst event at rate *λ*, increasing the population by *B*, or a death event at rate *δ*, decreasing it by one. The probability generating function of this process *G*(*z, t*) = E [*z*^*n*(*t*)^] satisfies

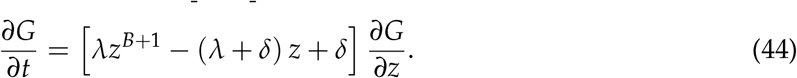

Differentiating Eq. (44) at *z* = 1 gives

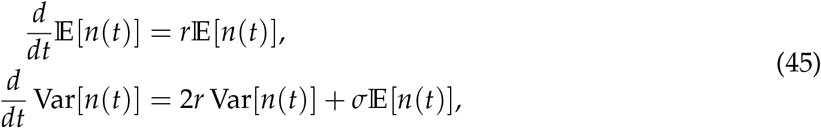

where

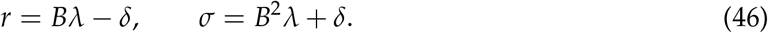

The second expression can equivalently be written as

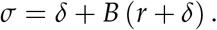

Thus, both the drift *r* and demographic-variance coefficient *σ* of the Feller diffusion equation 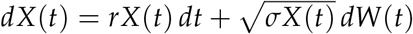 are obtained directly from the probability generating function of the discrete burst-death process.

We tested whether the Feller process approximates the complete population-size distribution of the discrete burst-death process in two settings: a supercritical benchmark relevant to growing lineages and a subcritical benchmark relevant to lineages declining toward extinction. In both cases, the Feller coefficients were determined from the microscopic burst-death parameters through Eq. (46).

#### Supercritical benchmark

We simulated 7,000 independent trajectories starting from

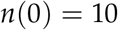

and used

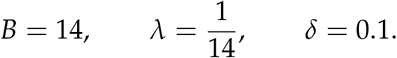

The corresponding Feller coefficients were

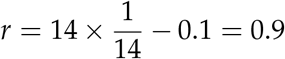

and

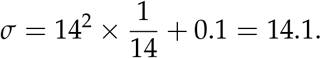

Figure D2a compares the empirical population-size distributions with the transition laws of the Feller diffusion having these theoretically derived coefficients.

The agreement across the three observation times shows that the Feller diffusion accurately approximates the expected growth of the discrete burst-death process and its population-size distribution. The close agreement across the three observation times provides a numerical check of the diffusion approximation in a supercritical regime relevant to the early growth of establishing viral lineages.

#### Subcritical benchmark

To assess the approximation for declining lineages, as required for the *de novo* rescue calculation, we also simulated 7,000 independent trajectories starting from

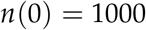

and used

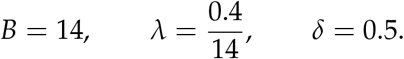

The corresponding net growth rate was negative:

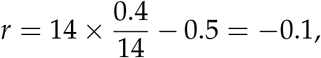

and the demographic-variance coefficient was

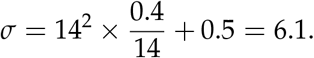

Figure D2b compares the simulated positive population-size distributions with the corresponding Feller transition laws. The agreement shows that the diffusion approximation captures the distributional dynamics of the discrete burst-death process as the population declines toward extinction.

Together, the two benchmarks show that the Feller transition law provides an accurate approximation to the positive population-size distributions of the discrete burst-death process in both growing and declining regimes. The subcritical benchmark specifically supports the use of a Feller diffusion to describe the declining parental lineages that produce *de novo* mutants before extinction.

**Figure D2:**
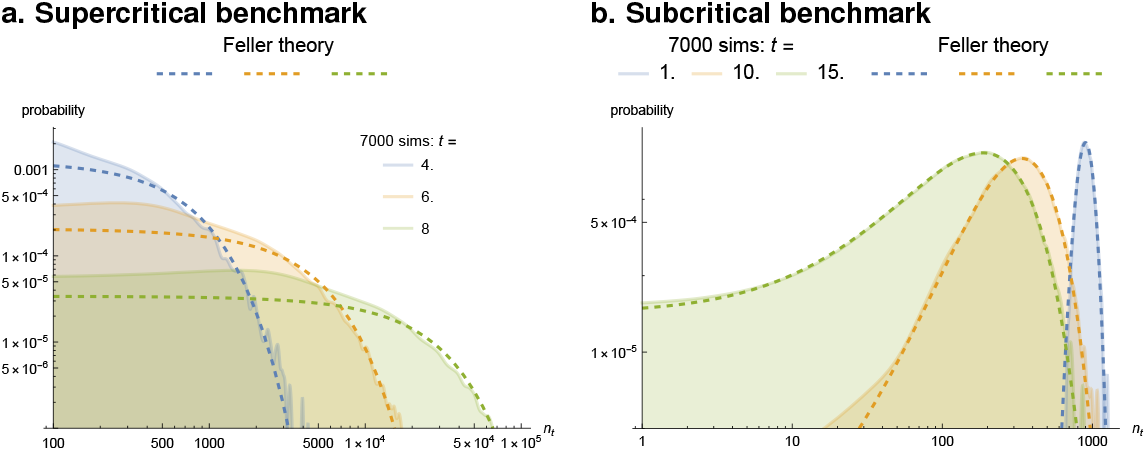
Individual-based validation of the Feller approximation for discrete burst-death processes. Filled curves show the positive population-size distributions obtained from 7,000 independent individual-based simulations. Dashed curves show the positive parts of the corresponding Feller transition laws, using coefficients derived directly from the exact probability generating function. (a) Supercritical benchmark. Simulations started from *n*(0) = 10 and used a net burst increment *B* = 14, a burst-event rate *λ* = 1/14, and a death rate *δ* = 0.1, giving *r* = 0.9 and *σ* = 14.1. Distributions are shown at *t* = 4, 6, and 8. (b) Subcritical benchmark. Simulations started from *n*(0) = 1000 and used a net burst increment *B* = 14, a burst-event rate *λ* = 0.4/14, and a death rate *δ* = 0.5, giving *r* = − 0.1 and *σ* = 6.1. Distributions are shown at *t* = 1, 10, and 15.

**Figure E1:**
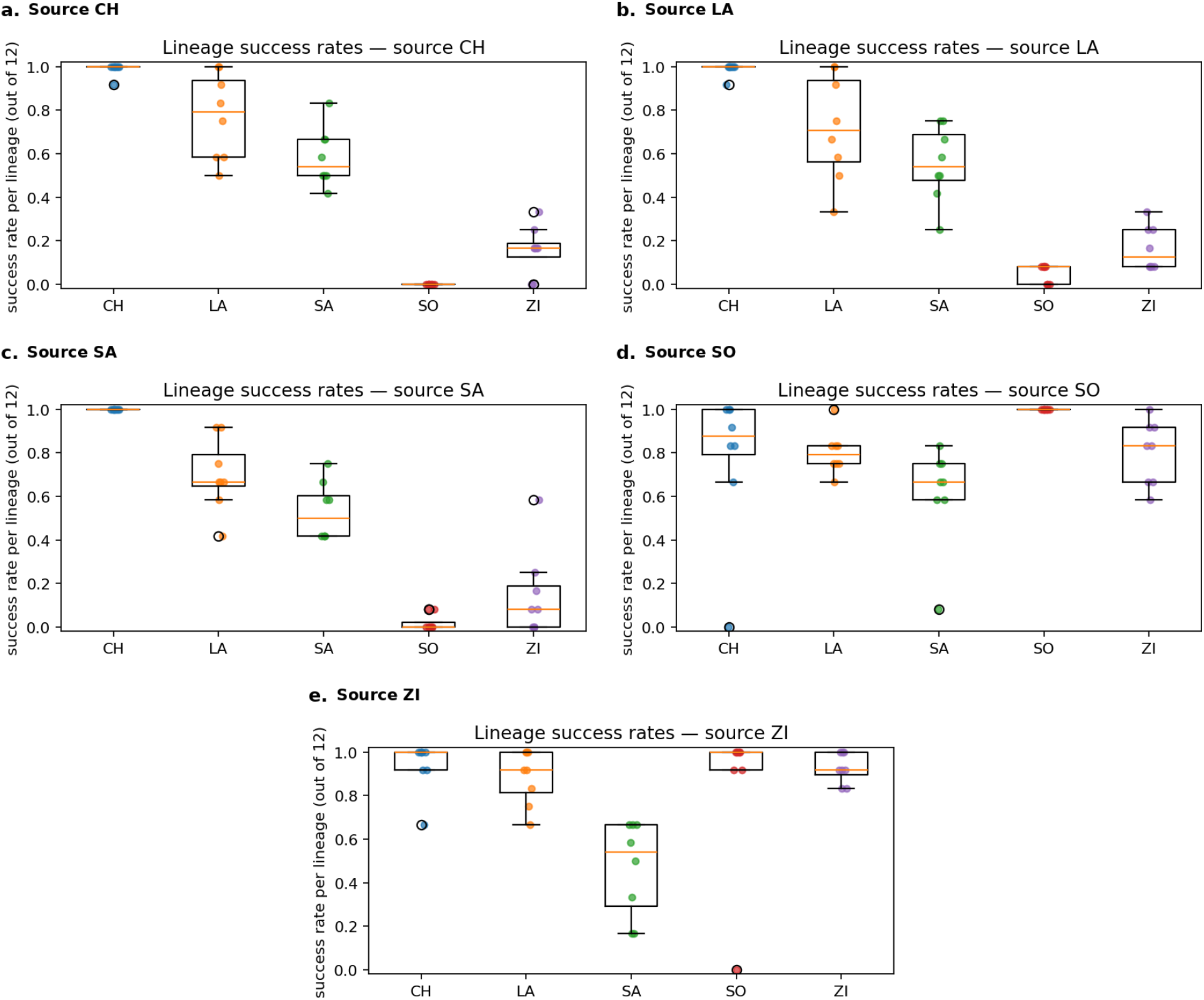
Lineage-level cross-infection outcomes. For each source host (panel title), boxplots show the distribution across the 8 evolved lineages of the number of successful infections out of 12 plants, for each target host. The spread across lineages motivates an explicit overdispersion model.

## E Additional results on the statistical analysis of successful infection outcomes

### Lineage-level variability across sources

#### Overdispersion diagnostics and estimation of the Beta-Binomial concentration

This appendix quantifies lineage-to-lineage heterogeneity in cross-infection outcomes and motivates the Beta-Binomial observation model used in Section “Observation model and likelihood“. The analysis below is purely statistical: it uses only the lineage-level counts 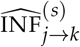 (8 lineages per evolved source host *j*, 12 plants per lineage-target pair) and does not involve the mechanistic model of Section “Mechanistic model“.

**Figure E2:**
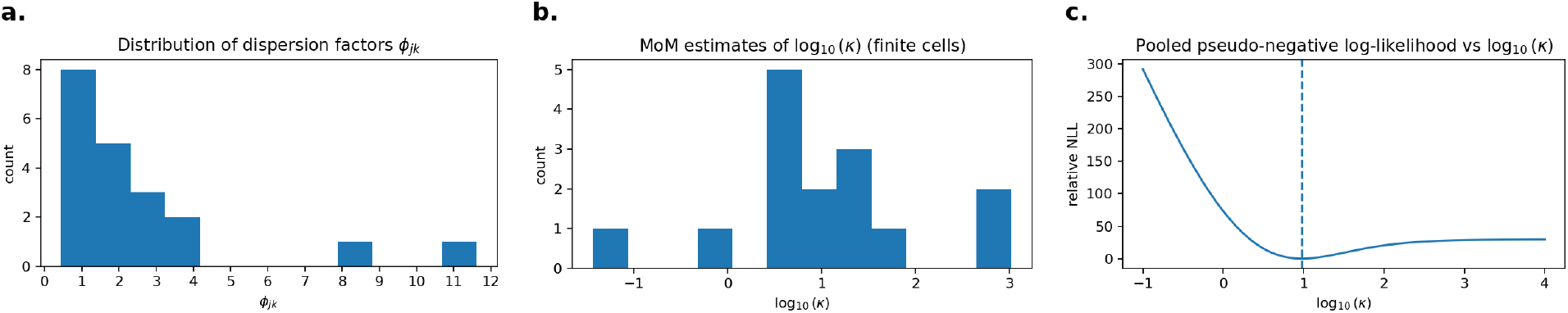
Overdispersion and Beta-Binomial concentration. (a) Distribution of per-cell dispersion factors *ϕ*_*jk*_. (b) Distribution of method-of-moments estimates of *κ* across cells with finite estimates (shown on a log_10_ scale). (c) Pooled pseudo-negative log-likelihood as a function of log_10_, with optimum at 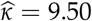.

#### Lineage-level data within a cell

For each evolved source host *j* ∈ {CH,LA,SA,SO,ZI} and target host *k* ∈ {CH,LA,SA,SO,ZI}, we observe

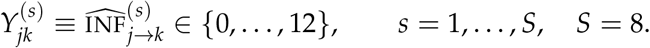

Define the within-cell empirical mean success probability (aggregated over lineages)

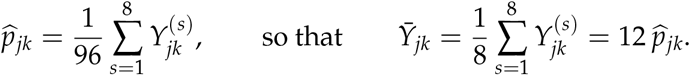

#### Dispersion factor

Under a Binomial(12, *p*_*jk*_) model with a common success probability across lineages in cell (*j, k*), the expected variance of 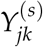 is 12 *p*_*jk*_(1 − *p*_*jk*_). We quantify extra-binomial variability by comparing the observed among-lineage variance to the Binomial variance evaluated At 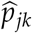. Let

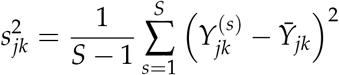

be the sample variance across lineages. The dispersion factor is

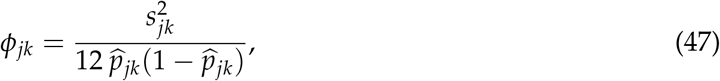

where, in practice, cells with 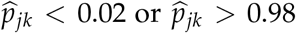 were excluded because this diagnostic is unstable near the boundaries. Values *ϕ*_*jk*_ *>* 1 indicate overdispersion relative to the Binomial model.

Panel (a) of Figure E2 shows the empirical distribution of *ϕ*_*jk*_ across the *H*^2^ = 25 evolved (source, target) cells (*H* = 5 hosts). We also report a pooled dispersion estimate obtained as the mean of the cellwise dispersion factors:

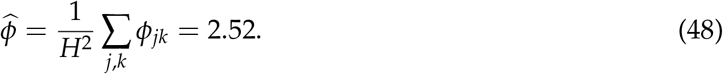

This pooled value indicates substantial extra-binomial variability at the lineage level (in practice, the average was computed only over the non-excluded pairs).

#### Beta-Binomial model

To model overdispersion, we use the hierarchical Beta-Binomial model (11):

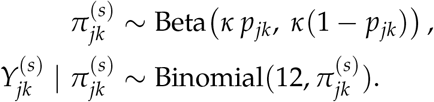

Marginally, 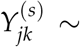 BetaBinomial(12, *κp*_*jk*_, *κ*(1 − *p*_*jk*_)). Under this model, the variance of 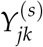 is inflated relative to Binomial(12, *p*_*jk*_) by the factor

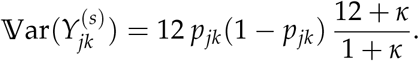

This suggests an estimator obtained by matching the empirical variance 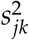 to the above expression with *p*_*jk*_ replaced by 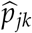. Using (47), this yields

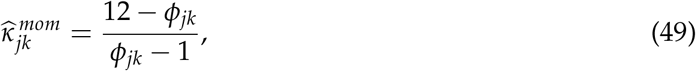

whenever 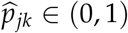 and *ϕ*_*jk*_ ∈ (1, 12) (otherwise the moment estimate is undefined or non-positive and is discarded). Panel (b) of Figure E2 shows the distribution of the finite 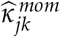 values on a log_10_scale. Across cells with finite moment estimates, the median is 7.54 and the geometric mean is 10.31.

#### Pooled pseudo-likelihood estimate of *κ*

Because the method-of-moments estimator can be unstable in small samples (only *S* = 8 lineages) and can fail when 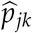 is near 0 or 1, we also estimate a single pooled *κ* by maximizing a pseudo-log-likelihood that conditions on the empirical cell means 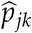. Specifically, for each cell we plug 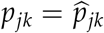 and consider the Beta-Binomial probability mass function

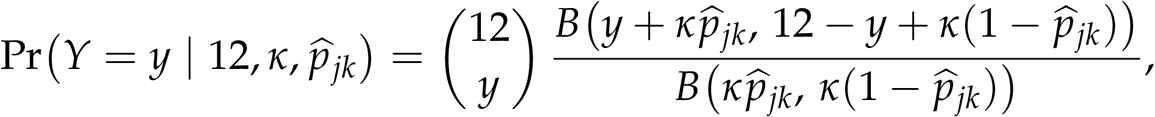

where *B*(·, ·) denotes the Beta function (for numerical stability, values 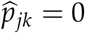 and 1 were replaced by 10^−^ and 1 − 10^−^, respectively). The pooled pseudo-log-likelihood is then

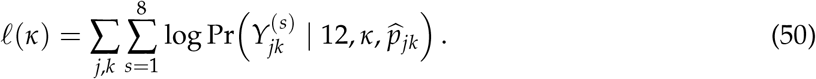

Panel (c) of Figure E2 plots the corresponding pseudo-negative log-likelihood as a function of log_10_ and shows a clear optimum at 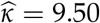.

#### Choice for mechanistic inference

The dispersion diagnostics (Figure E2) consistently indicate marked lineage-to-lineage heterogeneity, motivating the Beta-Binomial observation model (12). In the mechanistic inference, we fix *κ* = 10, close to both the pooled pseudo-likelihood estimate (9.50) and the moment estimates.

## F Descriptive analysis: MDS of the aggregated cross-inoculation profiles

As a descriptive complement to the mechanistic analysis, we represented the aggregated source profiles by classical multidimensional scaling (MDS). This analysis uses only the observed cross-inoculation matrix and does not rely on the fitness-landscape model.

For each source *j* ∈ {CL, CH, LA, SA, SO, ZI} and target *k* ∈ {CH, LA, SA, SO, ZI}, let 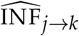 denote the observed number of successful infections and *N*_jk_ the corresponding number of inoculated deplants. We first transformed each observed infection count into a finite empirical logit,

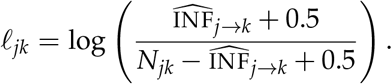

The 0.5 correction avoids infinite values when the observed infection rate is exactly 0 or 1.

Each source *j* is then represented by the five-dimensional transformed profile

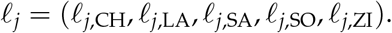

We computed Euclidean distances between source profiles,

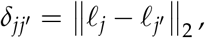

thereby obtaining a 6 × 6 descriptive distance matrix

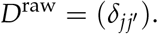

Classical MDS was then applied to this descriptive distance matrix. Specifically, with

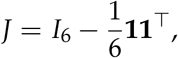

we formed the double-centered matrix

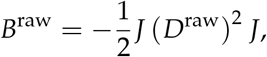

where (*D*^raw^)^2^ is the matrix of squared distances. If

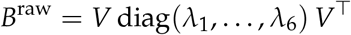

is the spectral decomposition of *B*^raw^, with eigenvalues ordered decreasingly, the MDS coordinates are obtained from the positive eigenvalues and associated eigenvectors. Specifically, for object *i*, the coordinate along axis *a* is

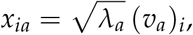

where (*v*_*a*_)_*i*_ is the *i*-th component of eigenvector *v*_*a*_. The two-dimensional representation shown in Figure F1 uses the first two positive coordinates. These two axes captured 96.9% of the total positive eigenvalue mass.

**Figure F1:**
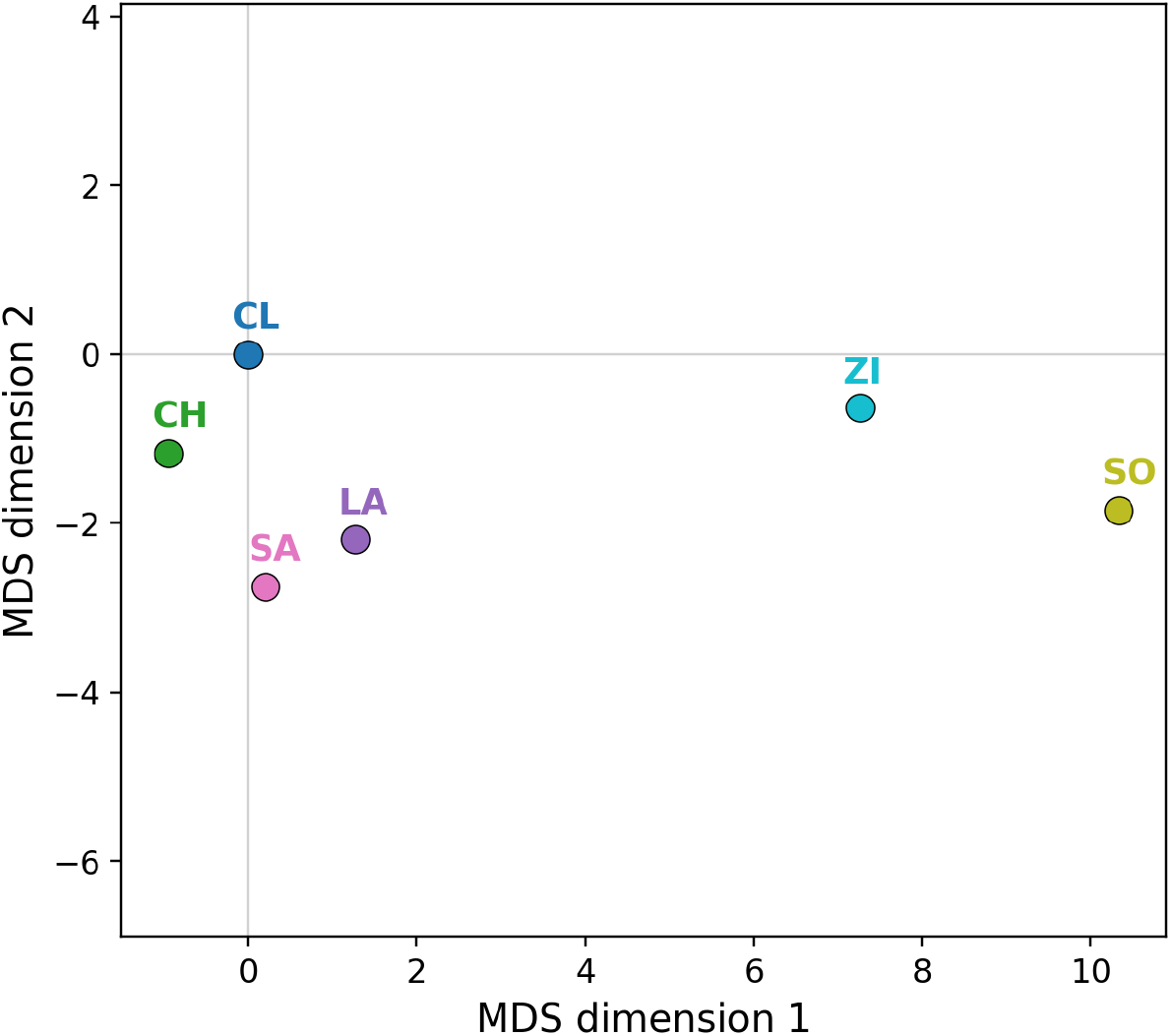
Classical MDS of the aggregated cross-inoculation profiles. Each source profile was constructed from the empirical-logit-transformed infection counts across the five target hosts, and Euclidean distances were computed between these five-dimensional profiles.

## G Predictive checks and model comparison

### Posterior predictive checks (PPC)

We assess model adequacy using posterior predictive checks based on aggregated infection success rates. Let 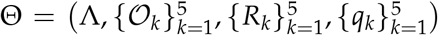 denote the parameter vector of the distance-only model (with *O*_CL_ = **0** and fixed *κ* = 10), and let *p*(Θ | data) be the posterior distribution sampled by MCMC. The posterior predictive distribution is

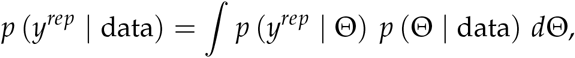

where *y*^*rep*^ denotes a replicated dataset generated under the fitted model.

### Observed aggregated rates used in the PPC

The PPC is computed on a vector of aggregated success rates that combines (i) clonal inoculations and (ii) evolved cross-inoculations aggregated over lineages:

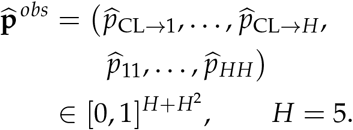

#### Clonal component

For each target host *k* ∈ {1, …, *H*},

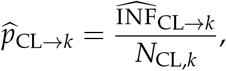

where 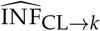 is the number of successful infections among *N*_CL,*k*_ inoculated plants.

#### Evolved component (aggregated over lineages)

For each source host *j* ∈ {1, …, *H*} and target host *k* ∈ {1, …, *H*},

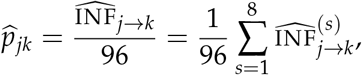

where 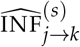 is the number of successes among 12 plants for lineage *s* evolved on host *j*, and 96 = 8× 12 is the total number of plants per (source, target) cell when aggregating over the 8 lineages.

### Replicated datasets under the fitted model

For each posterior draw Θ^(*t*)^ (for *t* = 1, …, *T*), we compute the mechanistic successful infection probabilities 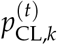 and 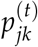 from (6), then simulate a replicated dataset using exactly the observation model of Section “Observation model and likelihood“.

#### Clonal inoculations

For each target host *k*,

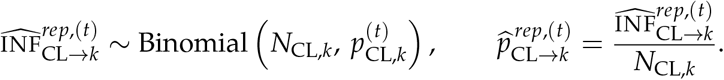

#### Lineage-level evolved inoculations

For each source host *j*, target host *k*, and lineage *s* ∈ {1, …, 8}, we draw a lineage-specific probability

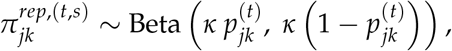

then conditional on 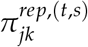 we simulate lineage-level successes

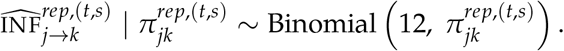

To match the aggregated rates used in the PPC, we sum over lineages and rescale:

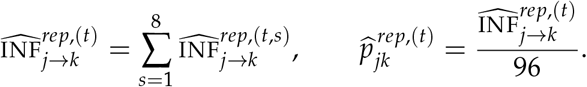

This produces a replicated aggregated-rate vector 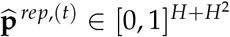 in the same order as 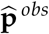.

### Posterior predictive summaries shown in the figures

For each component *i* ∈{1, …, *H* + *H*^2^} of the aggregated-rate vector, we compute the posterior predictive mean

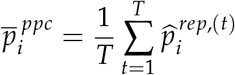

and the 95% posterior predictive interval

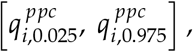

where 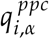 is the empirical *α*-quantile of 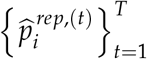.

In the PPC panel of Figure 3a, points plot 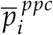 against the corresponding observed rate 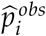,and vertical bars represent 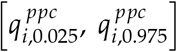. As a single summary of discrepancy on aggregated rates we report the RMSE,

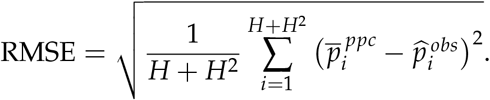

### Predictive model comparison with WAIC

To compare candidate phenotype dimensions *n* on a common predictive scale, we use the Widely Applicable Information Criterion (WAIC), a fully Bayesian criterion that approximates leave-one-out cross-validation using the pointwise log-likelihood (Watanabe, 2010). WAIC is computed from posterior draws of the log-likelihood contributions for each observation, and estimates the expected log pointwise predictive density (elpd), penalized by an effective number of parameters.

Let *y*_1_, …, *y*_*N*_ denote the *N* observational units used in the likelihood (here *N* = 205, combining *H* = 5 clonal Binomial outcomes and *H* ×*H*× 8 = 200 lineage-level Beta-Binomial outcomes). For posterior draws *θ*^(*s*)^, *s* = 1, …, *S*, define the pointwise log-likelihood

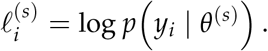

WAIC is based on two components. First, the log pointwise predictive density (lppd) is

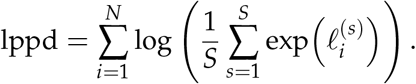

Second, the WAIC effective complexity penalty is the sum of pointwise posterior variances,

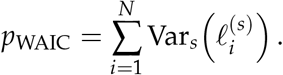

The expected log predictive density estimate is then

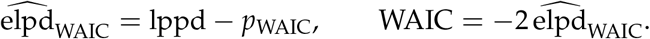

Higher 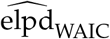 (equivalently, lower WAIC) indicates better expected out-of-sample predictive performance. Because WAIC is computed pointwise, differences in 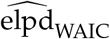 between two models can be summarized using the pointwise contributions 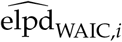 to obtain an approximate standard error:

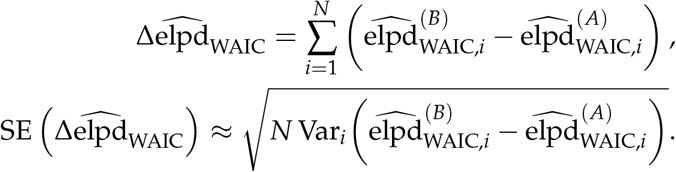

In the main text we report WAIC and 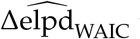 to compare phenotype dimensions *n*, using the observation-level likelihood contributions described above.

## References

Y Anciaux, L-M Chevin, O Ronce, and G Martin. Evolutionary rescue over a fitness landscape. Genetics, 209:265–279, 2018. doi: 10.1534/genetics.118.300908.

Yoann Anciaux, Amaury Lambert, Ophélie Ronce, Lionel Roques, and Guillaume Martin. Population persistence under high mutation rate: from evolutionary rescue to lethal mutagenesis. Evolution, 73(8):1517–1532, 2019. doi: 10.1111/evo.13771.

Rowan D. H. Barrett and Dolph Schluter. Adaptation from standing genetic variation. Trends in Ecology & Evolution, 23(1):38–44, 2008. doi: 10.1016/j.tree.2007.09.008.

Trevor Bedford, Marc A. Suchard, Philippe Lemey, Gytis Dudas, Victoria Gregory, Alan J. Hay, John W. McCauley, Colin A. Russell, Derek J. Smith, and Andrew Rambaut. Integrating influenza antigenic dynamics with molecular evolution. eLife, 3:e01914, 2014. doi: 10.7554/eLife.01914.

Graham Bell. Evolutionary rescue and the limits of adaptation. Philosophical Transactions of the Royal Society B: Biological Sciences, 368(1610):20120080, 2013. doi: 10.1098/rstb.2012.0080.

Bence Bognár, Réka Spohn, and Viktória Lázár. Drug combinations targeting antibiotic resistance. npj Antimicrobials and Resistance, 2(1):29, 2024. doi: 10.1038/s44259-024-00047-2. URL https://doi.org/10.1038/s44259-024-00047-2.

Tobias Bollenbach. Antimicrobial interactions: Mechanisms and implications for drug discovery and resistance evolution. Current Opinion in Microbiology, 27:1–9, 2015. doi: 10.1016/j.mib.2015.05.008. URL https://doi.org/10.1016/j.mib.2015.05.008.

Ingwer Borg and Patrick J. F. Groenen. Modern Multidimensional Scaling: Theory and Applications. Springer, New York, 2 edition, 2005.

Thierry Caquet, Chantal Gascuel, and Michèle Tixier-Boichard, editors. Agroécologie: des recherches pour la transition des filières et des territoires. Quae, 2020.

Stephanie M. Carlson, Curry J. Cunningham, and Peter A. H. Westley. Evolutionary rescue in a changing world. Trends in Ecology & Evolution, 29(9):521–530, 2014. doi: 10.1016/j.tree.2014.06.005.

F de la Iglesia, F Martínez, J Hillung, J M. Cuevas, P J. Gerrish, J-A Daròs, and SF. Elena. Luria– Delbrück Estimation of Turnip Mosaic Virus Mutation Rate In Vivo. Journal of Virology, 86: 3386–3388, 2012. doi: 10.1128/JVI.06909-11.

J. Arjan G. M. de Visser and Joachim Krug. Empirical fitness landscapes and the predictability of evolution. Nature Reviews Genetics, 15:480–490, 2014. doi: 10.1038/nrg3744.

Michael M. Desai and Daniel S. Fisher. Beneficial mutation-selection balance and the effect of linkage on positive selection. Genetics, 176(3):1759–1798, 2007. doi: 10.1534/genetics.106.067678.

William Feller. Diffusion processes in genetics. In Jerzy Neyman, editor, Proceedings of the second Berkeley symposium on mathematical statistics and probability, volume 2, pages 227–246. University of California Press, 1951.

Inês Fragata, Alexandre Blanckaert, Marco António Dias Louro, David A. Liberles, and Claudia Bank. Evolution in the light of fitness landscape theory. Trends in Ecology & Evolution, 34(1):69–82, 2019. doi: 10.1016/j.tree.2018.10.009.

Sergey Gavrilets. Fitness Landscapes and the Origin of Species, volume 41 of Monographs in Population Biology. Princeton University Press, Princeton, NJ, 2004. ISBN 9780691117584.

Gregory S. Gilbert and Campbell O. Webb. Phylogenetic signal in plant pathogen-host range. Proceedings of the National Academy of Sciences of the United States of America, 104(12):4979–4983, 2007. doi: 10.1073/pnas.0607968104.

Richard Gomulkiewicz and Robert D. Holt. When does evolution by natural selection prevent extinction? Evolution, 49(1):201–207, 1995. doi: 10.1111/j.1558-5646.1995.tb05971.x.

Richard Gomulkiewicz, Stephen M Krone, and Christopher H Remien. Evolution and the duration of a doomed population. Evolutionary applications, 10(5):471–484, 2017. doi: 10.1111/eva.12467.

John C. Gower. Generalized Procrustes analysis. Psychometrika, 40(1):33–51, 1975. doi: 10.1007/BF02291478.

Serafín Gutiérrez, Yannis Michalakis, and Stéphane Blanc. Virus population bottlenecks during within-host progression and host-to-host transmission. Current Opinion in Virology, 2(5):546–555, 2012. doi: 10.1016/j.coviro.2012.08.001.

Noémie Harmand, Romain Gallet, Guillaume Martin, and Thomas Lenormand. Evolution of bacteria specialization along an antibiotic dose gradient. Evolution Letters, 2(3):221–232, 2018. doi: 10.1002/evl3.52.

B. D. Harrison. The infectivity of extracts made from leaves at intervals after inoculation with viruses. Journal of General Microbiology, 15:210–220, 1956. doi: 10.1099/00221287-15-1-210.

Bahram Houchmandzadeh. General formulation of luria-delbrück distribution of the number of mutants. Physical Review E, 92(1):012719, 2015. doi: 10.1103/PhysRevE.92.012719.

JE Hubbarde, G Wild, and LM Wahl. Fixation probabilities when generation times are variable: The burst–death model. Genetics, 176(3):1703–1712, 2007. doi: 10.1534/genetics.107.072009.

Lejla Imamovic and Morten O. A. Sommer. Use of collateral sensitivity networks to design drug cycling protocols that avoid resistance development. Science Translational Medicine, 5 (204):204ra132, 2013. doi: 10.1126/scitranslmed.3006609. URL https://doi.org/10.1126/scitranslmed.3006609.

Thibaud Jayet, Lucie Tamisier, Marion Szadkowski, Elise Lepage, Grégory Girardot, Loup Rimbaud, Véronique Lefebvre, and Benoît Moury. The Replicative Fitness and Virulence of Potato Virus Y Evolve Differently in Pepper Lines with Different Levels of Resistance and Tolerance. Journal of General Virology, 107:002208, 2026. doi: 10.1099/jgv.0.002208.

Stuart A. Kauffman and Simon Levin. Towards a general theory of adaptive walks on rugged landscapes. Journal of Theoretical Biology, 128(1):11–45, 1987. doi: 10.1016/S0022-5193(87)80029-2.

Felicia Keesing, Robert D. Holt, and Richard S. Ostfeld. Effects of species diversity on disease risk. Ecology Letters, 9(4):485–498, 2006. doi: 10.1111/j.1461-0248.2006.00885.x.

Amaury Lambert. Population dynamics and random genealogies. Stochastic models, 24(sup1): 45–163, 2008. doi: 10.1080/15326340802437728.

Douglas E Lea and Charles Alfred Coulson. The distribution of the numbers of mutants in bacterial populations. Journal of genetics, 49(3):264–285, 1949. doi: 10.1007/BF02986080.

Ben Longdon, Michael A. Brockhurst, Colin A. Russell, John J. Welch, and Francis M. Jiggins. The evolution and genetics of virus host shifts. PLoS Pathogens, 10(11):e1004395, 2014. doi: 10.1371/journal.ppat.1004395.

Ailene MacPherson, Paul A. Hohenlohe, and Scott L. Nuismer. Trait dimensionality explains widespread variation in local adaptation. Proceedings of the Royal Society B: Biological Sciences, 282 (1802):20141570, 2015. doi: 10.1098/rspb.2014.1570.

Jennifer R Mandel, Rebecca B Dikow, Carolina M Siniscalchi, Ramhari Thapa, Linda E Watson, and Vicki A Funk. A fully resolved backbone phylogeny reveals numerous dispersals and explosive diversifications throughout the history of Asteraceae. Proceedings of the National Academy of Sciences, 116(28):14083–14088, 2019. doi: 10.1073/pnas.1903871116.

G Martin and T Lenormand. The fitness effect of mutations across environments: Fisher’s geometrical model with multiple optima. Evolution, 69(6):1433–1447, 2015. doi: 10.1111/evo.12671.

Guillaume Martin. Fisher’s geometrical model emerges as a property of complex integrated phenotypic networks. Genetics, 197(1):237–255, 2014. doi: 10.1534/genetics.113.160325.

Guillaume Martin and Lionel Roques. The nonstationary dynamics of fitness distributions: asexual model with epistasis and standing variation. Genetics, 204(4):1541–1558, 2016. doi: 10.1534/genetics.116.187385.

Guillaume Martin, Robin Aguilée, Johan Ramsayer, Oliver Kaltz, and Ophélie Ronce. The probability of evolutionary rescue: towards a quantitative comparison between theory and evolution experiments. Philosophical Transactions of the Royal Society B: Biological Sciences, 368(1610):20120088, 2013. doi: 10.1098/rstb.2012.0088.

A. McDonald and C. Linde. Pathogen population genetics, evolutionary potential, and durable resistance. Annual Review of Phytopathology, 40:349–379, 2002. doi: 10.1146/annurev.phyto.40.120501.101443. URL https://doi.org/10.1146/annurev.phyto.40.120501.101443.

Philipp W. Messer and Dmitri A. Petrov. Population genomics of rapid adaptation by soft selective sweeps. Trends in Ecology & Evolution, 28(11):659–669, 2013. doi: 10.1016/j.tree.2013.08.003. URL https://doi.org/10.1016/j.tree.2013.08.003.

Ramon Miralles, Philip J. Gerrish, Andrés Moya, and Santiago F. Elena. Clonal interference and the evolution of RNA viruses. Science, 285(5434):1745–1747, 1999. doi: 10.1126/science.285.5434.1745.

Benoît Moury, Marion Szadkowski, Catherine Wipf-Scheibel, Grégory Girardot, Julien Papaïx, Lionel Roques, Yamila Agrofoglio, Adrián A. Valli, Karine Berthier, and Cécile Desbiez. Common molecular determinants underlie potyvirus host species jumps and resistance breakdown. bioRxiv preprint, 2026. URL https://doi.org/10.64898/2026.07.09.737469. Preprint 10.64898/2026.07.09.737469.

C. C. Mundt. Durable resistance: A key to sustainable management of pathogens and pests. Infection, Genetics and Evolution, 27:446–455, 2014. doi: 10.1016/j.meegid.2014.01.011. URL https://doi.org/10.1016/j.meegid.2014.01.011.

Christopher C. Mundt. Use of multiline cultivars and cultivar mixtures for disease management. Annual Review of Phytopathology, 40:381–410, 2002. doi: 10.1146/annurev.phyto.40.011402.113723.

Daniel Nichol, Peter Jeavons, Alexander G. Fletcher, Robert A. Bonomo, Philip K. Maini, Jerome L. Paul, Robert A. Gatenby, Alexander R. A. Anderson, and Jacob G. Scott. Steering evolution with sequential therapy to prevent the emergence of bacterial antibiotic resistance. PLOS Computational Biology, 11(9):e1004493, 2015. doi: 10.1371/journal.pcbi.1004493. URL https://doi.org/10.1371/journal.pcbi.1004493.

Daniel Nichol, James Rutter, Christopher Bryant, Andrea M. Hujer, Stanley Lek, Mark D. Adams, Peter Jeavons, Alexander R. A. Anderson, Robert A. Bonomo, and Jacob G. Scott. Antibiotic collateral sensitivity is contingent on the repeatability of evolution. Nature Communications, 10(1):334, 2019. doi: 10.1038/s41467-018-08098-6. URL https://doi.org/10.1038/s41467-018-08098-6.

Matthew M Osmond, Sarah P Otto, and Guillaume Martin. Genetic paths to evolutionary rescue and the distribution of fitness effects along them. Genetics, 214(2):493–510, 2020. doi: 10.1534/genetics.119.302890.

Csaba Pál, Balázs Papp, and Viktória Lázár. Collateral sensitivity of antibiotic-resistant microbes. Trends in Microbiology, 23(7):401–407, 2015. doi: 10.1016/j.tim.2015.02.009. URL https://doi.org/10.1016/j.tim.2015.02.009.

Frank J. Poelwijk, Daniel J. Kiviet, Daniel M. Weinreich, and Sander J. Tans. Empirical fitness landscapes reveal accessible evolutionary paths. Nature, 445(7126):383–386, 2007. doi: 10.1038/nature05451.

Maija E Pollari, William WE Aspelin, Linping Wang, and Kristiina M Mäkinen. The molecular maze of potyviral and host protein interactions. Annual review of virology, 11(1):147–170, 2024. doi: 10.1146/annurev-virology-100422-034124.

Loup Rimbaud, Frédéric Fabre, Julien Papaïx, Benoît Moury, Christian Lannou, Luke G. Barrett, and Peter H. Thrall. Models of plant resistance deployment. Annual Review of Phytopathology, 59 (1):125–152, 2021. doi: 10.1146/annurev-phyto-020620-122134. URL https://doi.org/10.1146/annurev-phyto-020620-122134.

Raina S Robeva and John R Jungck. Fascination with fluctuation: Luria and Delbrück’s legacy. Axioms, 12(3):280, 2023. doi: 10.3390/axioms12030280.

Jason R. Rohr, David J. Civitello, Frank W. Halliday, Peter J. Hudson, Kevin D. Lafferty, Chelsea L. Wood, and Erin A. Mordecai. Toward common ground in the biodiversity–disease debate. Nature Ecology & Evolution, 4(1):24–33, 2020. doi: 10.1038/s41559-019-1060-6.

Roderich Römhild and Dan I. Andersson. Mechanisms and therapeutic potential of collateral sensitivity to antibiotics. PLOS Pathogens, 17(1):e1009172, 2021. doi: 10.1371/journal.ppat.1009172. URL https://doi.org/10.1371/journal.ppat.1009172.

Roderich Römhild, Tobias Bollenbach, and Dan I. Andersson. The physiology and genetics of bacterial responses to antibiotic combinations. Nature Reviews Microbiology, 20:478–490, 2022. doi: 10.1038/s41579-022-00700-5. URL https://doi.org/10.1038/s41579-022-00700-5.

Derek J. Smith, Alan S. Lapedes, Jan C. de Jong, Theo M. Bestebroer, Guus F. Rimmelzwaan, Albert D. M. E. Osterhaus, and Ron A. M. Fouchier. Mapping the antigenic and genetic evolution of influenza virus. Science, 305(5682):371–376, 2004. doi: 10.1126/science.1097211.

Erik I. Svensson and Ryan Calsbeek, editors. The Adaptive Landscape in Evolutionary Biology. Oxford University Press, Oxford, 2012. ISBN 9780199595372. doi: 10.1093/acprof:oso/9780199595372.001.0001.

Lucie Tamisier, Elsa Rousseau, Sebastien Barraillé, Ghislaine Nemouchi, Marion Szadkowski, Ludovic Mailleret, Frederic Grognard, Frederic Fabre, Benoît Moury, and Alain Palloix. Quantitative trait loci in pepper control the effective population size of two RNA viruses at inoculation. Journal of General Virology, 98(7):1923–1931, 2017. doi: 10.1099/jgv.0.000835.

O Tenaillon. The utility of Fisher’s geometric model in evolutionary genetics. Annual Review of Ecology, Evolution, and Systematics, 45:179–201, 2014. doi: 10.1146/annurev-ecolsys-120213-091846.

Mike Tyers and Gerard D. Wright. Drug combinations: a strategy to extend the life of antibiotics in the 21st century. Nature Reviews Microbiology, 17(3):141–155, 2019. doi: 10.1038/s41579-018-0141-x. URL https://doi.org/10.1038/s41579-018-0141-x.

Sumio Watanabe. Asymptotic equivalence of Bayes cross validation and widely applicable information criterion in singular learning theory. Journal of Machine Learning Research, 11:3571–3594, 2010.

Daniel M. Weinreich, Nigel F. Delaney, Mark A. DePristo, and Daniel L. Hartl. Darwinian evolution can follow only very few mutational paths to fitter proteins. Science, 312(5770):111–114, 2006. doi: 10.1126/science.1123539.

GA Whitmore and V Seshadri. A heuristic derivation of the inverse gaussian distribution. The American Statistician, 41(4):280–281, 1987. doi: 10.1080/00031305.1987.10475503.

M. S. Wolfe. The current status and prospects of multiline cultivars and variety mixtures for disease resistance. Annual Review of Phytopathology, 23:251–273, 1985. doi: 10.1146/annurev.py.23.090185.001343. URL https://doi.org/10.1146/annurev.py.23.090185.001343.

Mark E. J. Woolhouse, Louise H. Taylor, and Daniel T. Haydon. Population biology of multihost pathogens. Science, 292(5519):1109–1112, 2001. doi: 10.1126/science.1059026.

Sewall Wright. The roles of mutation, inbreeding, crossbreeding and selection in evolution. In Donald F. Jones, editor, Proceedings of the Sixth International Congress of Genetics, Ithaca, New York, 1932, volume 1, pages 356–366. Brooklyn Botanic Garden, Menasha, WI, 1932.

Pamela J. Yeh, Matthew J. Hegreness, Aviva Presser Aiden, and Roy Kishony. Drug interactions and the evolution of antibiotic resistance. Nature Reviews Microbiology, 7(6):460–466, 2009. doi: 10.1038/nrmicro2133. URL https://doi.org/10.1038/nrmicro2133.

Jiasui Zhan, Peter H. Thrall, Julien Papaïx, Lianhui Xie, and Jeremy J. Burdon. Playing on a pathogen’s weakness: Using evolution to guide sustainable plant disease control strategies. Annual Review of Phytopathology, 53(1):19–43, 2015. doi: 10.1146/annurev-phyto-080614-120040. URL https://doi.org/10.1146/annurev-phyto-080614-120040.

Youyong Zhu, Hairu Chen, Jinghua Fan, Yunyue Wang, Yan Li, Jianbing Chen, JinXiang Fan, Shisheng Yang, Lingping Hu, Hei Leung, Tom W. Mew, Paul S. Teng, Zonghua Wang, and Christopher C. Mundt. Genetic diversity and disease control in rice. Nature, 406(6797):718–722, 2000. doi: 10.1038/35021046. URL https://doi.org/10.1038/35021046.

