## Supplementary pdf file for "Inferring the multi-host fitness landscape of endive necrotic mosaic virus from cross-inoculation experiments"

### Supplementary File 1
### Inferring the multi-host fitness landscape of endive necrotic mosaic virus from cross-inoculation experiments

Lionel Roques et al.

#### S1 Alternative phenotype dimensions ($n = 1$ and $n = 3$)

To assess the effect of phenotypic dimensionality on inference, we repeated the full inference pipeline for $n = 1$ and $n = 3$ trait dimensions. Figures S1 and S2 report, for each dimension, the aggregated posterior predictive check (PPC), the posterior mean distance matrix among the host optima (and clonal strain CL), an MDS visualization of the inferred geometry after Procrustes alignment, and the marginal posterior distributions of the host-specific radii $R_k$, the target-specific parameters $q_k$, and the mutation parameter $\Lambda = \log_{10}(U)$. For $n = 1$, the MDS embedding is displayed with the second coordinate fixed at 0 for visualization, and the posterior clouds are not shown. In both dimensions, the mean radii for LA and SA are not displayed in the MDS panels because they extend beyond the plotting range; for $n = 1$, the radius for SO is additionally too small to be visible.

Note that, although the $n = 3$ model is visualized in two dimensions in Figure S2c, this representation is a projection of a three-dimensional configuration. For each posterior draw, we computed the eigenvalues $\lambda_1 \geq \lambda_2 \geq \lambda_3$ of the double-centered matrix used in the MDS and quantified the fraction of total positive eigenvalue mass captured by the first two coordinates, $(\lambda_1 + \lambda_2) / (\lambda_1 + \lambda_2 + \lambda_3)$. This fraction has posterior mean 0.879 (median 0.879, 95% CI [0.781, 0.976]), implying that the third coordinate contributes a non-negligible share of the total mass (mean 12%, median 12%, 95% CI [2%, 22%]). Thus the inferred configuration is not strictly planar, although predictive criteria do not support $n = 3$ over $n = 2$ (the main-text section "Phenotype dimension").

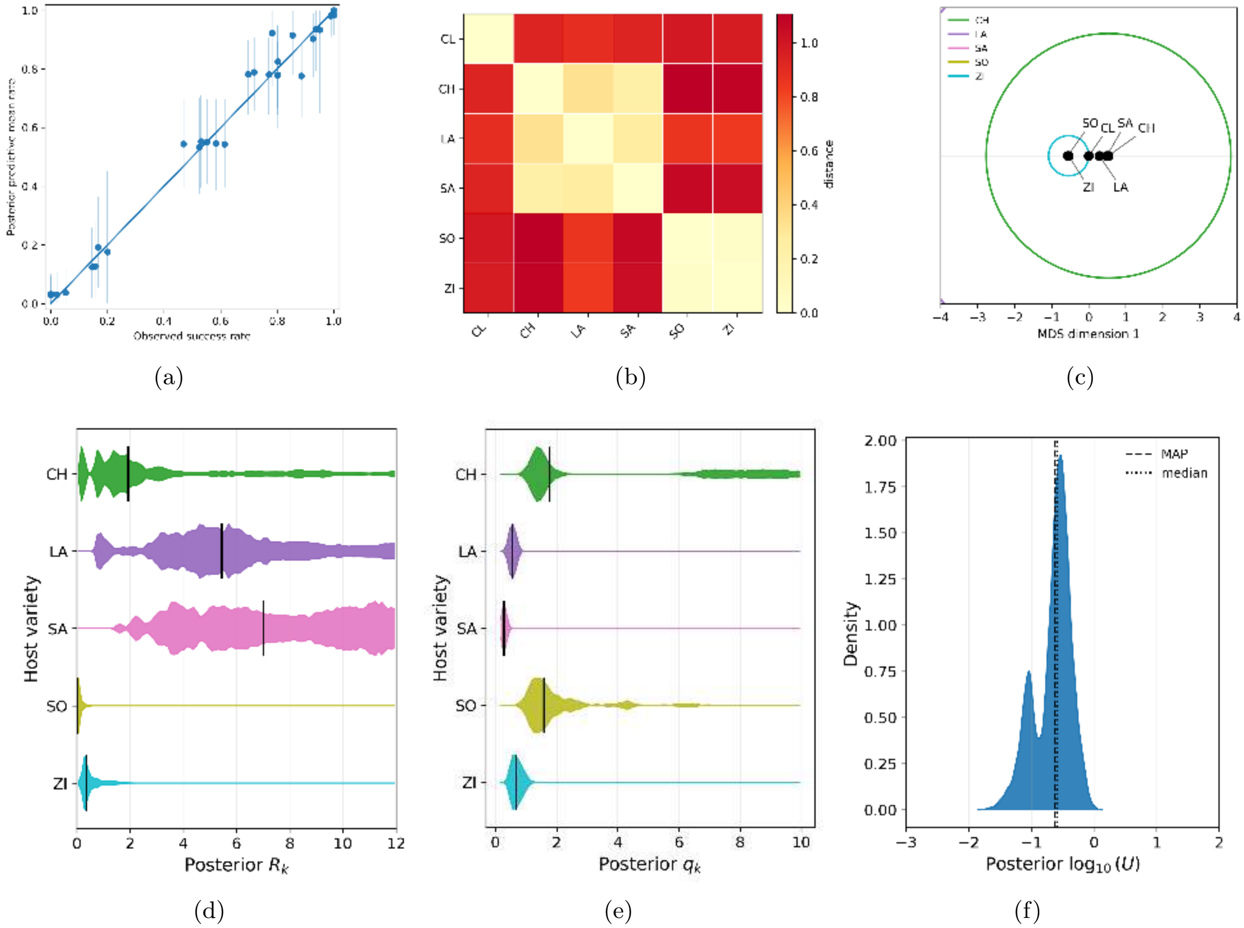


Figure S1: Posterior inference for $n = 1$ trait dimension. (a) Posterior predictive check comparing observed aggregated infection success rates with posterior predictive draws. (b) Posterior mean distance matrix among the clonal strain (CL) and the host optima. (c) MDS embedding, plotted with the second coordinate fixed at 0 for visualization, with circles representing the inferred target radii $R_k$ (posterior clouds are not shown). The mean radii for LA and SA are not displayed because they extend beyond the plotting range, while the radius for SO is too small to be visible. (d) Marginal posterior distributions of the parameters $R_k$. (e) Marginal posterior distributions of the parameters $q_k$. (f) Marginal posterior distribution of the mutation parameter $\Lambda = \log_{10}(U)$.

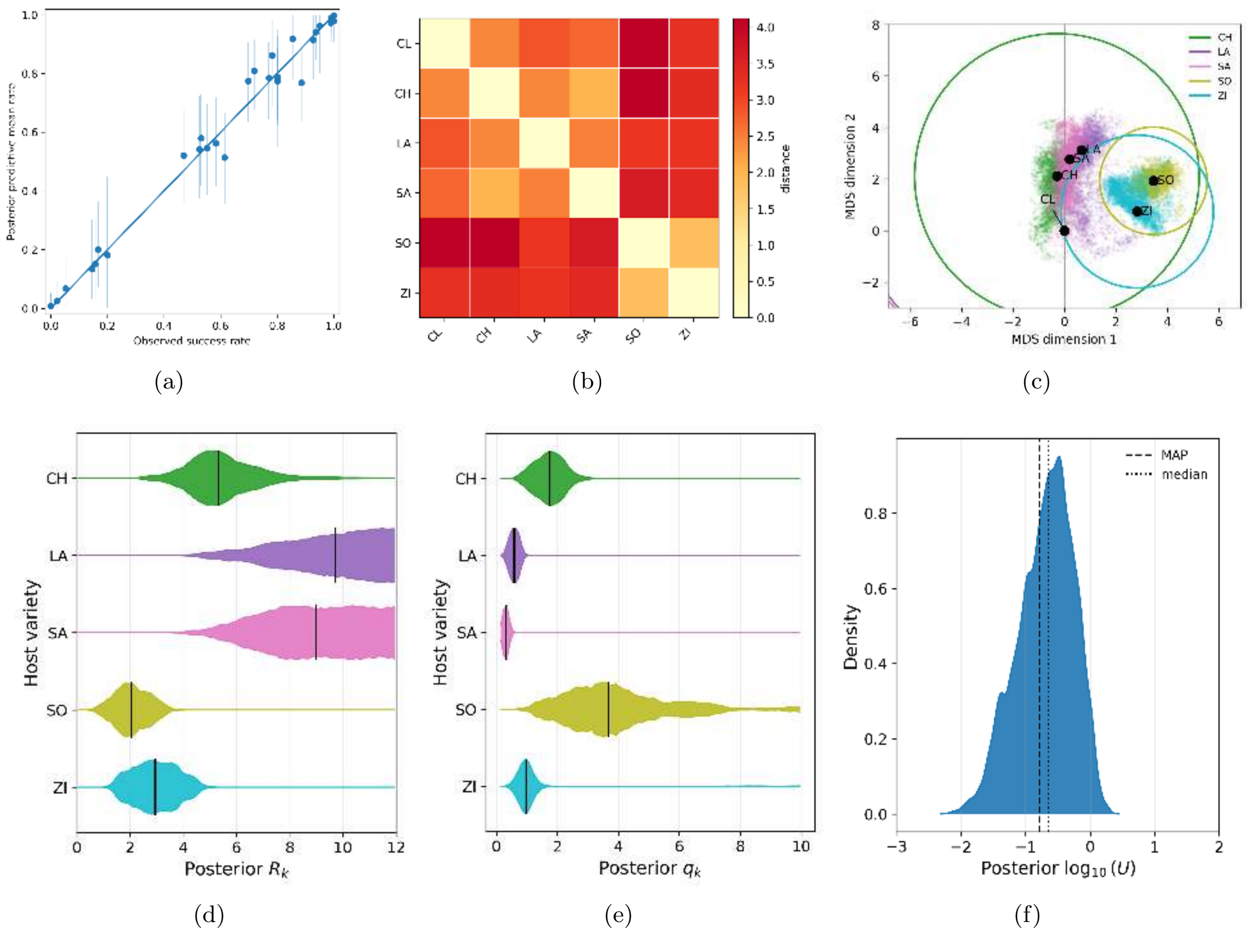


(a) (b) (c)

(d) (e) (f)

Figure S2: Posterior inference for $n = 3$ trait dimensions. (a) Posterior predictive check comparing observed aggregated infection success rates to posterior predictive draws. (b) Posterior mean distance matrix among the clonal strain (CL) and host optima. (c) MDS embedding with posterior uncertainty and circles representing inferred target radii $R_k$; the mean radii for LA and SA are not displayed because they extend beyond the plotting range. (d) Marginal posterior distributions of the parameters $R_k$. (e) Marginal posterior distributions of the parameters $q_k$. (f) Marginal posterior distribution of the mutation parameter $\Lambda = \log_{10}(U)$.

#### S2 CL-like source-inoculum model

The descriptive analysis of the raw cross-inoculation matrix showed that the CL, CH, LA and SA source rows are not detectably different once target-host effects are accounted for (the main-text section "Descriptive structure of the cross-inoculation matrix"). This raises the possibility that viral populations evolved on CH, LA and SA did not need to move to distinct adapted source phenotypes during Step 1 in order to explain their cross-inoculation profiles. The companion sequence data support this cautious interpretation rather than providing independent evidence for adaptive divergence on these three hosts: no nonsynonymous VPg substitution was detected in three randomly selected final populations from each of CH, LA, and SA, whereas recurrent nonsynonymous VPg substitutions were observed in all populations evolved on SO and ZI (Moury *et al.*, 2026).

To assess this point, we fitted an alternative model with $n = 2$ trait dimensions. In the baseline model, the effective source phenotype of an evolved inoculum from host $j$ is approximated by the corresponding source optimum, $\mathbf{x}_j \simeq \mathcal{O}_j$. In the alternative model considered here, the effective source phenotypes of the CH-, LA- and SA-evolved inocula were instead set to the clonal phenotype,

$$\mathbf{x}_j = \mathcal{O}_{\mathrm{CL}} = \mathbf{0}, \qquad j \in \{\mathrm{CH}, \mathrm{LA}, \mathrm{SA}\}.$$

Note that the hosts CH, LA and SA are still retained as distinct target environments, with their own optima $\mathcal{O}_k$, radii $R_k$, and infection-efficiency parameters $q_k$.

All other components of the inference pipeline were unchanged: the same mechanistic infection probability, priors, Binomial likelihood for the clonal inoculations, Beta-Binomial likelihood for lineage-level evolved inoculations, and fixed overdispersion parameter $\kappa = 10$ were used. The alternative model achieved essentially the same best fit as the baseline $n = 2$ model, with only a small decrease in maximum log-likelihood ($|\Delta \log L| = 1.3$). This indicates that the data do not require CH-, LA- and SA-evolved inocula to be modeled as distinct adapted source phenotypes.

Figure S3 summarizes the posterior inference under this alternative model. The posterior predictive check shows that the model reproduces the aggregated infection rates with accuracy comparable to the baseline $n = 2$ model. However, the inferred landscape is less directly interpretable than in the baseline model. In this alternative model, the CH, LA and SA source phenotypes are constrained to be identical to CL; consequently, the points labelled CH, LA and SA in the distance matrix and MDS plot should not be interpreted as source phenotypes. They represent only target-host optima, i.e. the phenotypic positions that determine how easily incoming inocula can establish in CH, LA and SA as target hosts. Distances involving these points therefore describe target-side constraints, not similarity among evolved source inocula. This loss of source-side interpretation also weakens the geometric constraints on their positions, which explains the broader posterior uncertainty for several target optima. In particular, target hosts with broad viable regions can be fit by a wide range of optimum positions once the corresponding source phenotypes are fixed to CL.

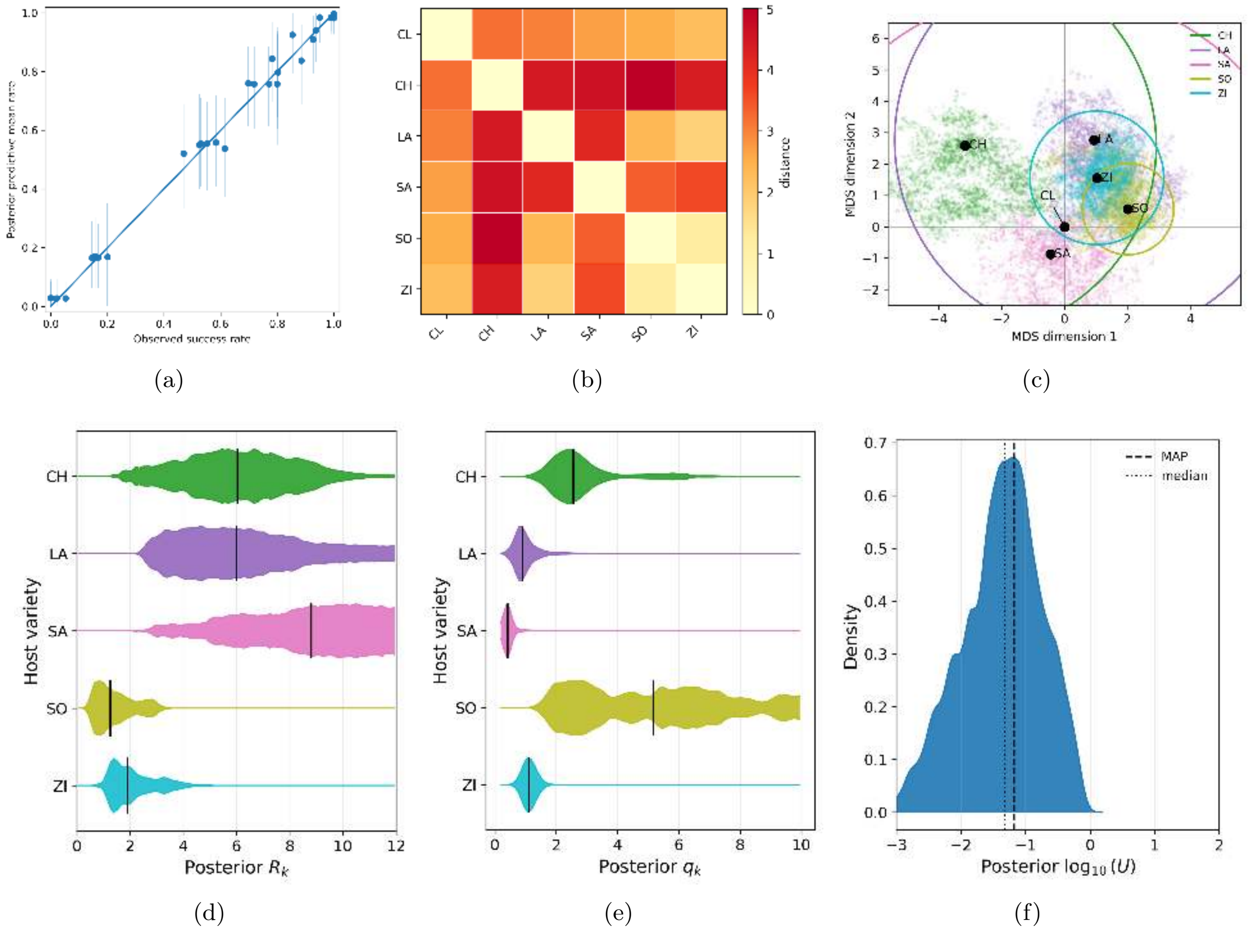


Figure S3: Posterior inference under the alternative CL-like source-inoculum model with $n = 2$ trait dimensions. In this model, the effective source phenotypes of CH-, LA- and SA-evolved inocula are fixed to the clonal phenotype, while CH, LA and SA remain distinct target hosts with their own optima, radii and infection-efficiency parameters. (a) Posterior predictive check comparing observed aggregated infection success rates with posterior predictive draws. (b) Posterior mean distance matrix among the clonal strain (CL) and host optima. (c) MDS embedding with posterior uncertainty and circles representing inferred target radii $R_k$; broad posterior clouds indicate weakly identified target positions under this alternative inoculum assumption. (d) Marginal posterior distributions of the host-specific radii $R_k$. (e) Marginal posterior distributions of the target-specific infection-efficiency parameters $q_k$. (f) Marginal posterior distribution of the mutation parameter $\Lambda = \log_{10}(U)$.

#### S3 Robustness to the source-standing-variation term

The derivation of the SV term assumes that mutant lineages generated during source-population growth contribute effectively to the sampled inoculum. This is a simplifying approximation. Source-derived mutant lineages may be lost stochastically before sampling, remain at very low abundance, or fail to be transmitted to the target plant.

To assess how strongly the inferred landscape depends on the effective magnitude of this term, we introduced a fixed attenuation factor $\alpha_{\mathrm{SV}}$ in the probability that no source-standing-variation lineage establishes:

$$\mathbb{P}\left(\overline{\mathrm{SV}};\alpha_{\mathrm{SV}}\right) = \exp\left[-\alpha_{\mathrm{SV}} q_k \nu_{jk}\right]. \tag{S1}$$

The direct-establishment and *de novo* rescue failure probabilities retain their baseline expressions:

$$\mathbb{P}\left(\overline{\mathrm{EST}}\right) = \exp\left[-2q_k r_{jk}^{+}\right]$$

and

$$\mathbb{P}\left(\overline{\mathrm{DN}} \mid \overline{\mathrm{EST}}\right) = \exp\left[-q_k\left(\sqrt{2\nu_{jk}+r_{jk}^2} - |r_{jk}|\right)\right].$$

The resulting total infection probability is therefore

$$\begin{aligned} p_{jk}\left(\alpha_{\mathrm{SV}}\right) &= 1 - \mathbb{P}\left(\overline{\mathrm{EST}}\right) \\ &\quad \times \mathbb{P}\left(\overline{\mathrm{DN}} \mid \overline{\mathrm{EST}}\right) \\ &\quad \times \mathbb{P}\left(\overline{\mathrm{SV}};\alpha_{\mathrm{SV}}\right) \\ &= 1 - \exp\Bigg[-2q_k r_{jk}^{+} - \alpha_{\mathrm{SV}} q_k \nu_{jk} \\ &\qquad\qquad - q_k\left(\sqrt{2\nu_{jk}+r_{jk}^2} - |r_{jk}|\right)\Bigg]. \end{aligned} \tag{S2}$$

The baseline model corresponds to $\alpha_{\mathrm{SV}} = 1$. We refitted the complete $n = 2$ model for two additional fixed values, $\alpha_{\mathrm{SV}} = 0.5$ and $\alpha_{\mathrm{SV}} = 0$. The value $\alpha_{\mathrm{SV}} = 0.5$ halves the term $q_k \nu_{jk}$ in the failure exponent, whereas $\alpha_{\mathrm{SV}} = 0$ gives

$$\mathbb{P}\left(\overline{\mathrm{SV}};\alpha_{\mathrm{SV}}\right) = 1$$

and therefore removes the SV route.

The alternative fits gave maximum log-likelihoods very close to that of the baseline model. The baseline model had maximum log-likelihood −320.47, whereas the models with $\alpha_{\mathrm{SV}} = 0.5$ and $\alpha_{\mathrm{SV}} = 0$ had maximum log-likelihoods −320.72 and −320.57, respectively. These small differences indicate that the binary cross-inoculation data discriminate only weakly among these fixed levels of the effective source-standing-variation term.

Figures S4 and S5 show the corresponding posterior summaries. In both alternative fits, the posterior predictive checks remain very similar to those of the baseline model, indicating that the observed aggregated infection rates are still well reproduced. The inferred distance matrices and MDS maps also preserve the main geometric structure: SO and ZI remain separated from the CL/Cichorieae group, whereas CH, LA, and SA occupy a more compact region of the inferred landscape. Likewise, the target radii retain the same qualitative interpretation, with SO remaining the most restrictive target and LA and SA remaining broad, permissive targets.

Attenuating or removing the SV term is partly compensated by changes in other inferred parameters, especially the effective mutation-supply parameter $U$ and the target-specific parameters

$q_k$ and $R_k$. This is expected because the binary endpoint data constrain the total infection probabilities more directly than they determine how successful infections should be attributed among the three routes. Nevertheless, the main conclusions are preserved: infection of SO from CL-, CH-, LA-, and SA-derived inocula remains strongly constrained, whereas SO- and ZI-derived inocula retain broad predicted infectivity. Thus, the assumed magnitude of the effective SV term affects the fitted parameters and the conditional route-attribution probabilities, but it does not drive the qualitative host-landscape structure inferred from the cross-inoculation data.

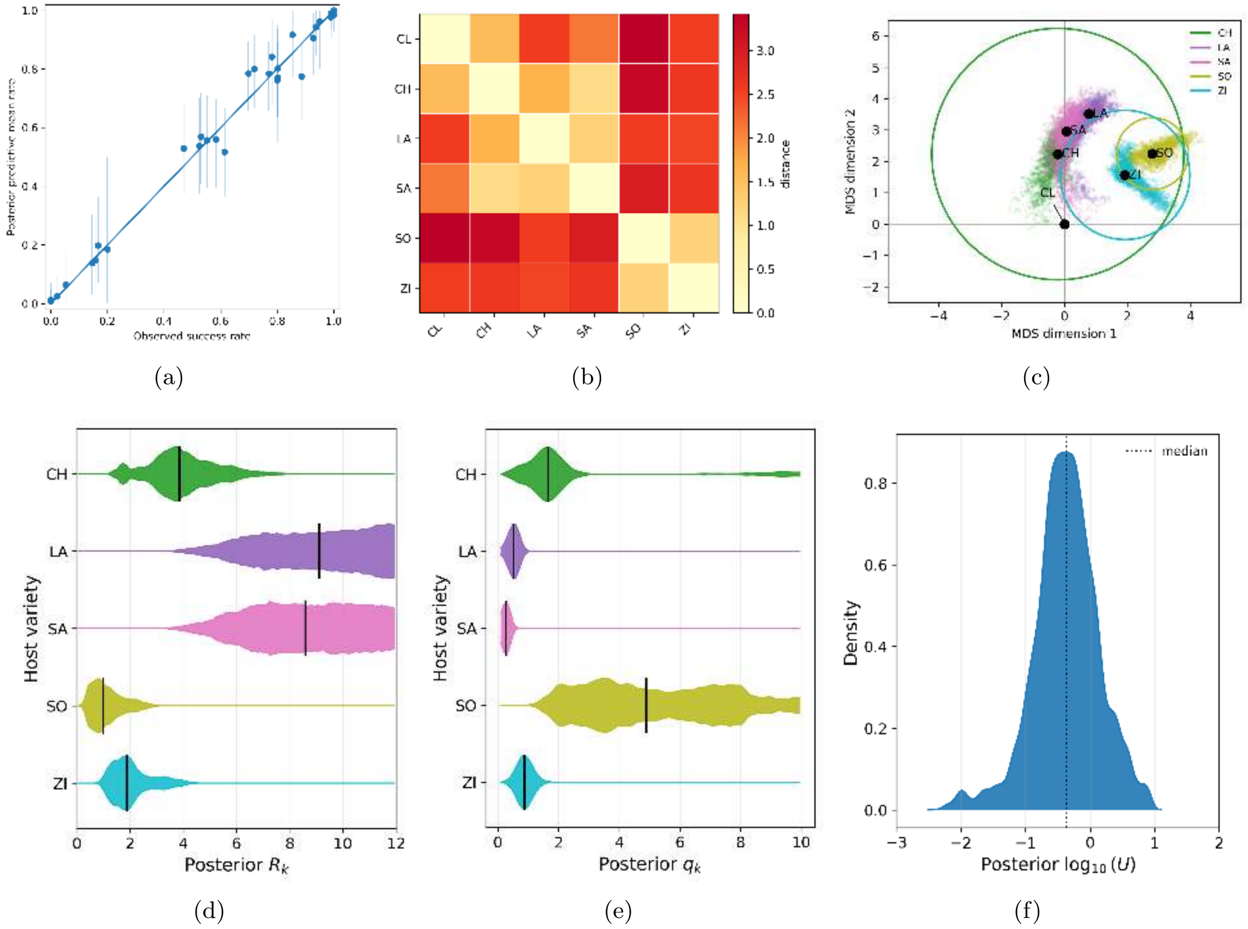


Figure S4: Robustness analysis with an attenuated source-standing-variation term, $\alpha_{\mathrm{SV}} = 0.5$, for the $n = 2$ model. The term $q_k \nu_{jk}$ is multiplied by $\alpha_{\mathrm{SV}}$ in $\mathbb{P}(\overline{\mathrm{SV}}; \alpha_{\mathrm{SV}})$, whereas the direct-establishment and *de novo* rescue failure probabilities retain their baseline expressions. (a) Posterior predictive check comparing observed aggregated infection success rates with posterior predictive draws. (b) Posterior mean distance matrix among the clonal strain (CL) and host optima. (c) MDS embedding with posterior uncertainty and circles representing inferred target radii $R_k$. (d) Marginal posterior distributions of the host-specific radii $R_k$. (e) Marginal posterior distributions of the target-specific infection-efficiency parameters $q_k$. (f) Marginal posterior distribution of the mutation parameter $\Lambda = \log_{10}(U)$.

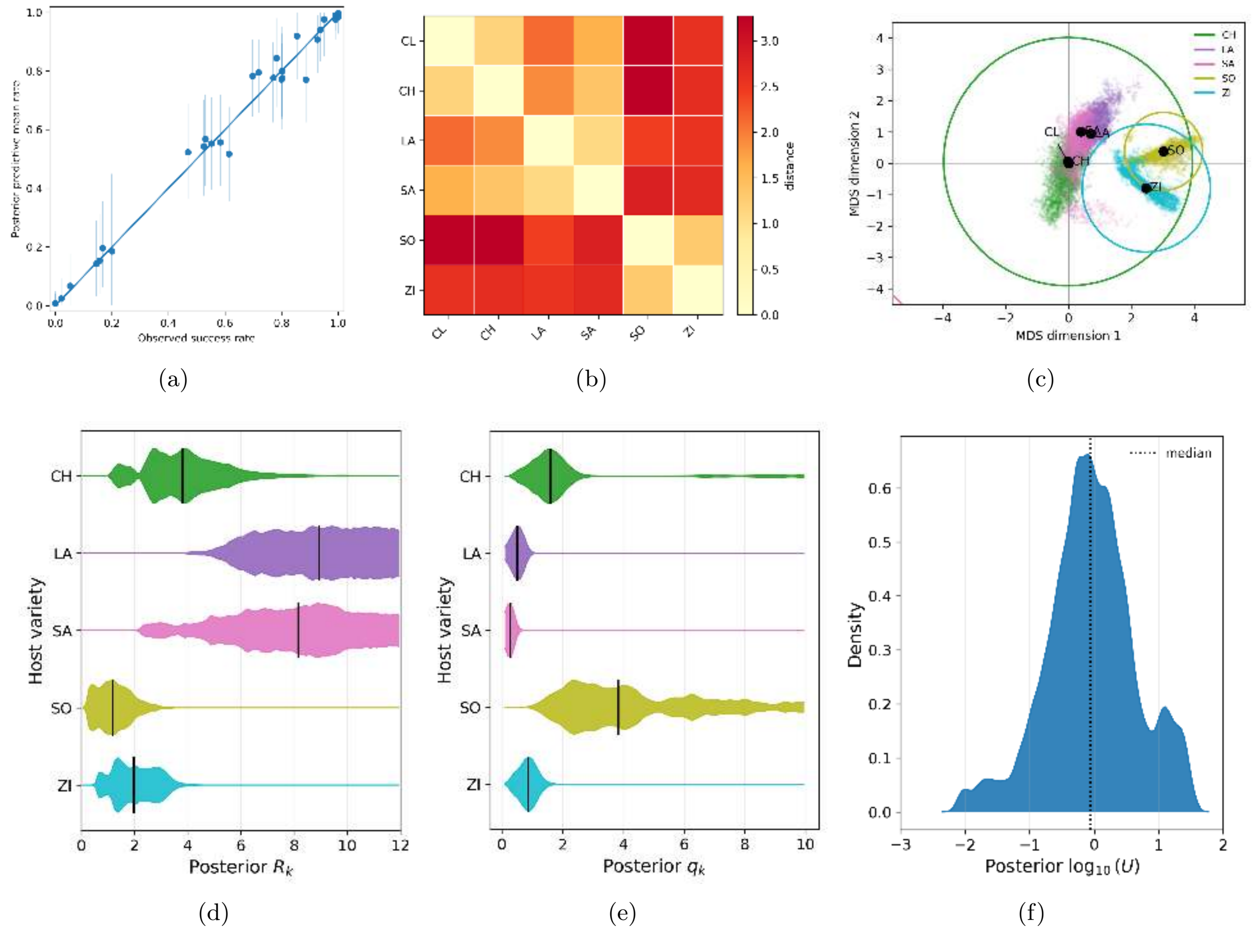


Figure S5: Robustness analysis with source-standing-variation rescue removed, $\alpha_{\text{SV}} = 0$, for the $n = 2$ model. In this case, $\mathbb{P}(\overline{\text{SV}}; \alpha_{\text{SV}}) = 1$, whereas the direct-establishment and *de novo* rescue failure probabilities retain their baseline expressions. (a) Posterior predictive check comparing observed aggregated infection success rates with posterior predictive draws. (b) Posterior mean distance matrix among the clonal strain (CL) and host optima. (c) MDS embedding with posterior uncertainty and circles representing inferred target radii $R_k$. (d) Marginal posterior distributions of the host-specific radii $R_k$. (e) Marginal posterior distributions of the target-specific infection-efficiency parameters $q_k$. (f) Marginal posterior distribution of the mutation parameter $\Lambda = \log_{10}(U)$.

### References


Moury B, Szadkowski M, Wipf-Scheibel C, Girardot G, Papaïx J *et al*. 2026. Common molecular determinants underlie potyvirus host species jumps and resistance breakdown. bioRxiv. pp. 2026–07.
